# Gene Novelties in Amphioxus Illuminate the Early Evolution of Cephalochordates

**DOI:** 10.1101/2022.05.18.492404

**Authors:** Qing Xiong, Kevin Yi Yang, Xi Zeng, Mingqiang Wang, Patrick Kwok-Shing Ng, Jun-Wei Zhou, Judy Kin-Wing Ng, Cherie Tsz-Yiu Law, Qiao Du, Kejin Xu, Bingyu Mao, Stephen Kwok-Wing Tsui

## Abstract

Amphioxus is considered the best-known living proxy to the chordate ancestor and an irreplaceable model organism for evolutionary studies of chordates and deuterostomes. In this study, a high-quality genome of the Beihai amphioxus, *Branchiostoma belcheri* beihai, was *de novo* assembled and annotated. Within four amphioxus genomes, twenty-eight groups of gene novelties were identified, revealing new genes that lack homologs in non-deuterostome metazoa, but share unexpectedly high similarities with those from non-metazoan species. These gene innovation events have played roles in amphioxus adaptations, including innate immunity responses, glycolysis, and regulation of calcium balance. The gene novelties related to innate immunity, such as a group of lipoxygenases and a DEAD-box helicase, boosted amphioxus immune responses. The novel genes for alcohol dehydrogenase and ferredoxin could aid in the glycolysis of amphioxus. A proximally arrayed cluster of EF-hand calcium-binding protein genes were identified to resemble those of bacteria. The copy number of this gene cluster was negatively correlated to the sea salinity of the collection region, suggesting that it may enhance their survival at different calcium concentrations. This comprehensive study collectively reveals insights into adaptive evolution of cephalochordates and provides valuable resources for research on early evolution of deuterostomes.

## Background

Amphioxus, a group of non-vertebrate chordates within Cephalochordata, diverged from the common ancestor of other two chordate groups, urochordates (tunicates) and vertebrates, around 550 million years ago (Bertrand and Escriva, 2011; Bourlat et al., 2006; Delsuc et al., 2006). Compared with tunicates, which were considered as a rapidly-evolving group (Bourlat et al., 2006), amphioxus shares a basic chordate body plan, undergoes slower molecular evolution (Benito-Gutiérrez et al., 2021; Putnam et al., 2008). Therefore, amphioxus is considered an irreplaceable model organism for evolutionary studies. With the advent of high-throughput sequencing, genome-based investigation has revolutionized the study of amphioxus evolution and greatly expanded our knowledge of the origin and evolution of vertebrates (Acemel et al., 2016; Brasó-Vives et al., 2022a; Holland et al., 2008; Huang et al., 2014; Huang et al., 2023; Marlétaz et al., 2018; Putnam et al., 2008; Simakov et al., 2020).

Whole-genome comparison of the three chordate groups provided evidence supporting the 2R hypothesis, which posits that two rounds of whole-genome duplication occurred in the early vertebrate lineage (Holland et al., 2008; Putnam et al., 2008). The application of functional genomics to amphioxus has provided a wealth of information on the origins of the complex gene regulatory landscapes that are characteristic of vertebrates, shedding new light on the evolutionary history of this important group of animals (Marlétaz et al., 2018).

In this study, we successfully constructed a high-quality genome of the Beihai amphioxus, *Branchiostoma* (*B.*) *belcheri* beihai, which was collected from the nearshore area of South China. Although amphioxus is widely regarded as the most basally divergent group of chordates and lacks the specializations and genome innovations of vertebrates (Escriva, 2018; Holland, 2017; Kassahn et al., 2009; Louis et al., 2012), we performed a comparative genomic analysis of amphioxus and identified a number of gene novelties, where the genes involved markedly differ from those found in other non-deuterostome metazoa, but surprisingly exhibit similarities to those found in non-metazoan species such as bacteria, fungi, and plants. Our analysis suggests that these novel genes may play crucial roles in a variety of amphioxus adaptations, including innate immunity, glycolysis, and calcium balance. Overall, our comprehensive genomic analysis of the gene novelties in amphioxus provided insights into the genetic mechanisms that underlie the adaptations and diversification of chordates and deuterostomes.

## Methods

### Sample collection

The adults of the Beihai amphioxus, *B. belcheri* beihai, were obtained from the sea near Dianbai District, Maoming City, Guangdong Province, China, cultured at 24–28 °C with air-pumped circulating artificial seawater in Beihai Marine Station of Nanjing University in Beihai City, Guangxi Province, China, and fed with seawater and sea alga (Chen et al., 2009; Du et al., 2023; Yang et al., 2016). This *B. belcheri* beihai amphioxus population has been sustained for several years to attain a low degree of polymorphism.

### Genomic DNA extraction and sequencing

To obtain the genome information of *B. belcheri* Beihai, genomic DNA was extracted from a pool of amphioxus samples and subjected to both PacBio long-read and Illumina short-read sequencing. Genomic DNAs were extracted using the Qiagen Blood & Cell Culture DNA Maxi Kit (Qiagen, Germany). Firstly, adult bodies of the Beihai amphioxus were washed twice with phosphate-buffered saline (PBS, pH 7.4), immediately frozen in liquid nitrogen, and then homogenized into powder form using a mortar and pestle with liquid nitrogen to maintain a low temperature. The homogenized samples were then incubated at 50 °C for proteolysis following the manufacturer’s protocol. Genomic DNA was bound to the column, washed, and eluted by washing buffer and elution buffer respectively, and precipitated in 70% ethanol. Finally, air-dry pellets of genomic DNA were dissolved overnight in UltraPure™ DNase/RNase-free distilled water (Thermo Fisher Scientific, USA) overnight at room temperature. The integrity of genomic DNA was determined by electrophoresis in 0.5% agarose gel and analyzed by Agilent 2100 Bioanalyzer (Agilent Technologies, USA), and the quantity was detected by Nanodrop 2000 spectrophotometer (Thermo Fisher Scientific, USA) and Qubit Fluorometer (Thermo Fisher Scientific, USA). The genomic DNA of *B. belcheri* beihai was sequenced by Illumina HiSeq 2000 in 500-bp and 3,000-bp libraries to generate NGS reads and PacBio RS system for third-generation long reads.

### De novo genome assembly and annotation

The initial genome assembly of *B. belcheri* beihai was constructed with PacBio long reads using Canu v1.8 (Koren et al., 2017). Then, scaffolding was performed by SSPACE Basic v2.0 (Boetzer et al., 2011) with paired-end Illumina short reads (Bentley et al., 2008) and by SSPACE LongRead v1.1 (Boetzer and Pirovano, 2014) with PacBio long reads (Eid et al., 2009). The final sequence polishing was finished by Pilon v1.22 (Walker et al., 2014) with all Illumina short reads to generate the final genome assembly. The genome assembly was assessed by QUAST v5.0.2 (Gurevich et al., 2013) in continuity and the completeness of genome assembly was assessed by BUSCO v3.1.0 (Simão et al., 2015) with database metazoa_odb9.

With genome assembled, genome annotation was performed. Firstly, repeat masking was performed by *de novo* prediction with RepeatModeler v2.0.1 (Flynn et al., 2020) and masking with RepeatMasker v4.0.8 (RepBase edition 20181026) (Tarailo-Graovac and Chen, 2009).

In the *de novo* prediction with RepeatModeler v2.0.1 (Flynn et al., 2020), RECON v1.05 (Bao and Eddy, 2002) and RepeatScout v1.0.6 (Price et al., 2005) were used for the prediction of repeat family in the genome. Then, genome annotation was performed by Maker pipeline v2.31 (Cantarel et al., 2008). In the Maker pipeline, transcriptome assemblies and homologous proteins were used as evidence for alignment by Exonerate v2.4.0 (Slater and Birney, 2005), whilst gene prediction was completed by SNAP (lib v2017-03-01) (Korf, 2004), GeneMark v4.38 (Borodovsky and Lomsadze, 2011) and Augustus v3.3.1 (Hoff and Stanke, 2019). The quality of genome annotation was assessed by BUSCO v3.1.0 (Simão et al., 2015) with database metazoa_odb9.

### Data collection and access

Besides of the *de novo* assembled and annotated genome of *B. belcheri* beihai, genome assemblies and annotations of other three amphioxus were downloaded from NCBI database (NCBI Assembly accession GCF_001625305.1 for *B. belcheri* xiamen, GCF_000003815.2 for *B. floridae* and GCA_900088365.1 for *B. lanceolatum*). The genome sequencing data, assembly, and annotation of *B. belcheri* beihai have been uploaded to NCBI database under the BioProject accession: PRJNA804338. The transcriptome data of *B. belcheri* beihai were from the BioProject accession: PRJNA310680.

Despite the recent reports of several high-quality *B. lanceolatum* genomes (Brasó-Vives et al., 2022b), the absence of transcriptome data hindered our ability to use the genome in manual curation. We did not include other chromosome-level amphioxus genomes produced from hybrid breeding due to concerns about the phasing step and subsequent transcriptome analysis (Huang et al., 2023).

### Phylogenetic analysis of deuterostomes

The genome sequences of deuterostome species were phylogenetically analyzed with those of other species. The overlapped single and complete BUSCO protein sequences were extracted by BUSCO v3.1.0 (Simão et al., 2015) with database metazoa_odb9, then aligned by MAFFT (Katoh et al., 2002) and edited in Gblocks (Castresana, 2000) with the option ‘-t=p’ to generate sequence alignment of conserved amino-acid residues. Finally, the sequence alignment was used to construct the phylogenetic tree in maximum likelihood algorithm and 100 bootstrap replicates by RAxML v8.2.12 (Stamatakis, 2014) with the options ‘-m PROTCATWAG -f a -# 100’. The phylogenetic tree was edited by the online tool Interactive Tree of Life (iTOL) (Letunic and Bork, 2021).

To construct a ultrametric time tree, the alignment of codon sequences of conserved amino-acid residues were extracted and used for estimating divergence time by the MCMCTree program from the PAML package v4.9j (Yang, 2007). The ultrametric time tree was final edited by the online tool Interactive Tree of Life (iTOL) (Letunic and Bork, 2021).

### Phylogenomic orthology analysis

Phylogenomic orthology and gene gain/loss analysis (Hahn et al., 2005) was performed among protein sequences in 21 annotated genomes by OrthoFinder v2.5.4 (Emms and Kelly, 2015, 2019) and CAFÉ v4.2 (De Bie et al., 2006) based on the ultrametric time tree. All the protein sequences were assigned into orthogroups according to the protein similarities on sequence level. These orthogroups were also considered as gene families. Specific orthogroups were summarized and further analyzed. Functional clusters of orthologous groups (COGs) of genes in specific orthogroups were performed by eggnog-mapper v2.1.5 (Cantalapiedra et al., 2021; Huerta-Cepas et al., 2019).

### Identification of gene novelties

Our study primarily focuses on exploring the gene novelties of amphioxus within the framework of deuterostomes. Specifically, we concentrated on genes that exhibit notable differences from those present in other non-deuterostome metazoans, while also sharing similarities with genes from non-metazoan species, including bacteria, fungi, plants, and other eukaryotes. The whole workflow can be seen in Fig. S3.

To identify gene novelties in amphioxus, we compared the best hits of all the annotated proteins of *B. belcheri* beihai to those from different taxonomic groups in UniRef50 database (UniProt, last updated on Dec. 15^th^, 2020) (Suzek et al., 2015) as follows. Although amphioxus specific orthogroups have been extracted, we input all the annotated proteins of *B. belcheri* beihai, to avoid missing important gene novelties.

1. In UniRef50 database, collecting the proteins of different taxonomic groups according to the taxon identifiers, namely non-deuterostome metazoa, bacteria, fungi, plants, and other eukaryotes (excluding fungi, plants, and metazoa), and building BLAST databases.
2. Searching the best hits of 44,745 annotated proteins of *B. belcheri* beihai in the five BLAST databases built in Step 1, using BLASTP v2.9.0 (McGinnis and Madden, 2004) with the options ‘-evalue 1e-6 -max_hsps 1 -max_target_seqs 1 -outfmt 6’.
3. The best hits in different taxonomic groups were compared and identified two groups of candidates of gene novelties. Firstly, when one protein had hit in the databases of bacteria, fungi, plants, or other eukaryotes, but not that of non-deuterostome metazoan, this gene was identified as group I candidates of gene novelties. Secondly, when the gene novelty index of a protein was calculated as dividing the bit score of the best hit in the databases of bacteria, fungi, plants, and other eukaryotes, by that of non-deuterostome metazoan; if the index ≥ 1.5, this gene was identified as group II candidates. This method has been adapted from the one employed to identify horizontal gene transfer events (Xiong et al., 2022).
4. Collecting protein sequences of two groups of candidates, performing BLASTP online search in the NCBI non-redundant protein (NR) database (last updated on Nov. 1^st^, 2021), and then manually checking the top 100 hits. Except those from Amphioxiformes (including all amphioxus), if the best hits were from unusual taxonomic groups like bacteria, fungi, plants, and other eukaryotes (excluding fungi, plants, and metazoa), this gene was confirmed as an amphioxus specific gene; if some of the best hits were from other deuterostome groups except jawed vertebrates, this gene was considered as a novel gene conserved with other deuterostomes. Additionally, only the DBH and ADH genes had homologs from the soft coral *Dendronephthya gigantea* and the sponge *Amphimedon queenslandica*, respectively (Table 1).

Therefore, the dominantly abundant and well annotated proteins of jawed vertebrates were used to filter out the candidate genes possibly introduced at stem chordates and deuterostomes, to avoid repeated study in the previous report on in stem chordates (Holland et al., 2008) and deuterostomes (Simakov et al., 2015). The absence of the candidate genes in jawed vertebrates was double confirmed by searching using TBLASTN in the genomic DNA sequences and BLASTP in the annotated proteins at E-value cutoff of 1E-6. Those candidates supported by poor hits (e.g., low bit scores or repeated domains) were discarded either.

5. Once a candidate gene was confirmed, homologous genes of four amphioxus were searched by TBLASTN in the genomic DNA sequences and BLASTP in the annotated proteins at E-value cutoff of 1E-6. After again checking those homologous genes at Step 4 and filtering out false-positive genes; manual checking and curating based on transcriptome data, especially in splice sites (locations of exon–intron boundaries) using the program Integrative Genomics Viewer (IGV) v2.9.4 (Thorvaldsdóttir et al., 2013); finally, all homologous genes among four amphioxus (Table 1) were collected and further analyzed.

**Table 1.**
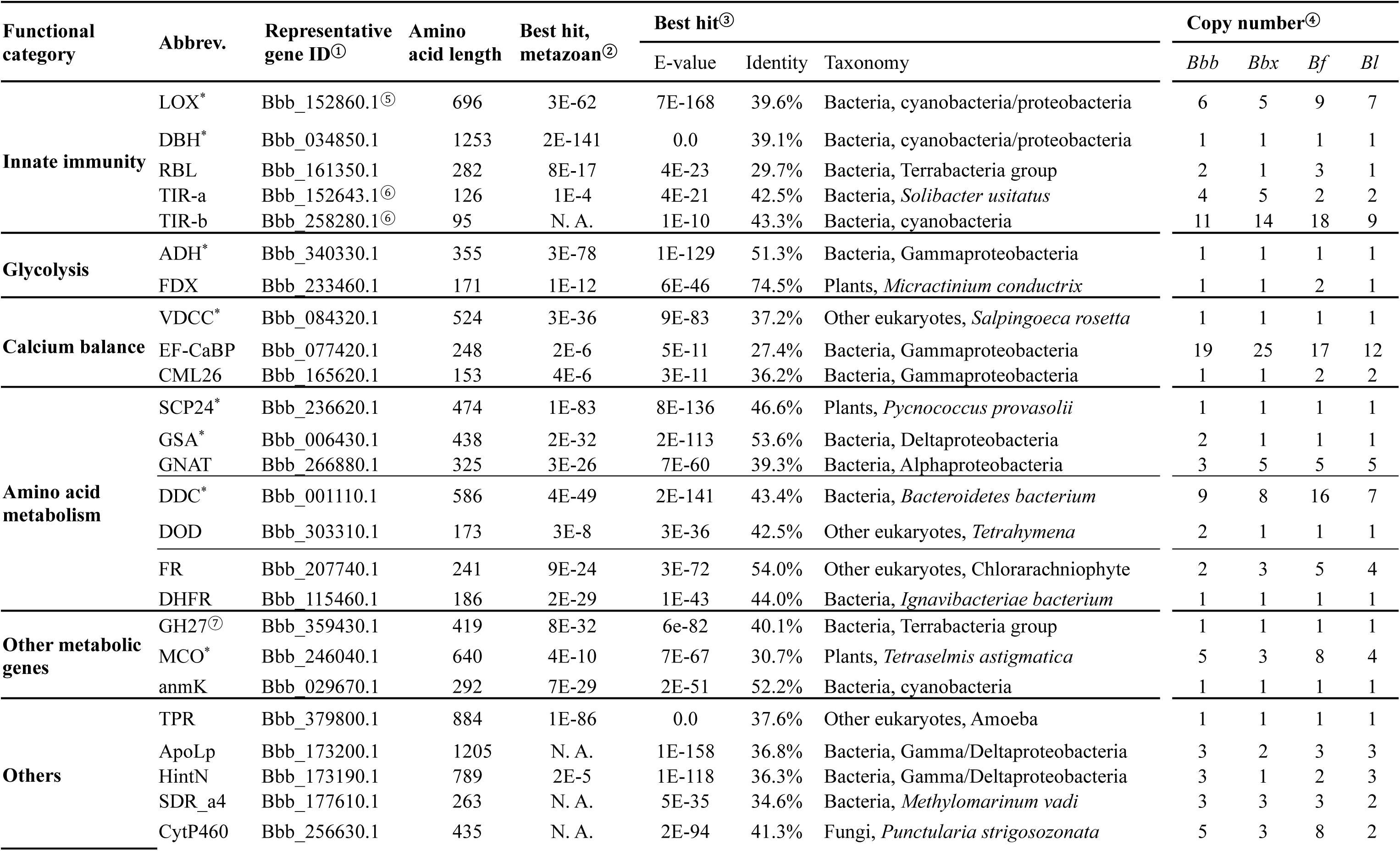
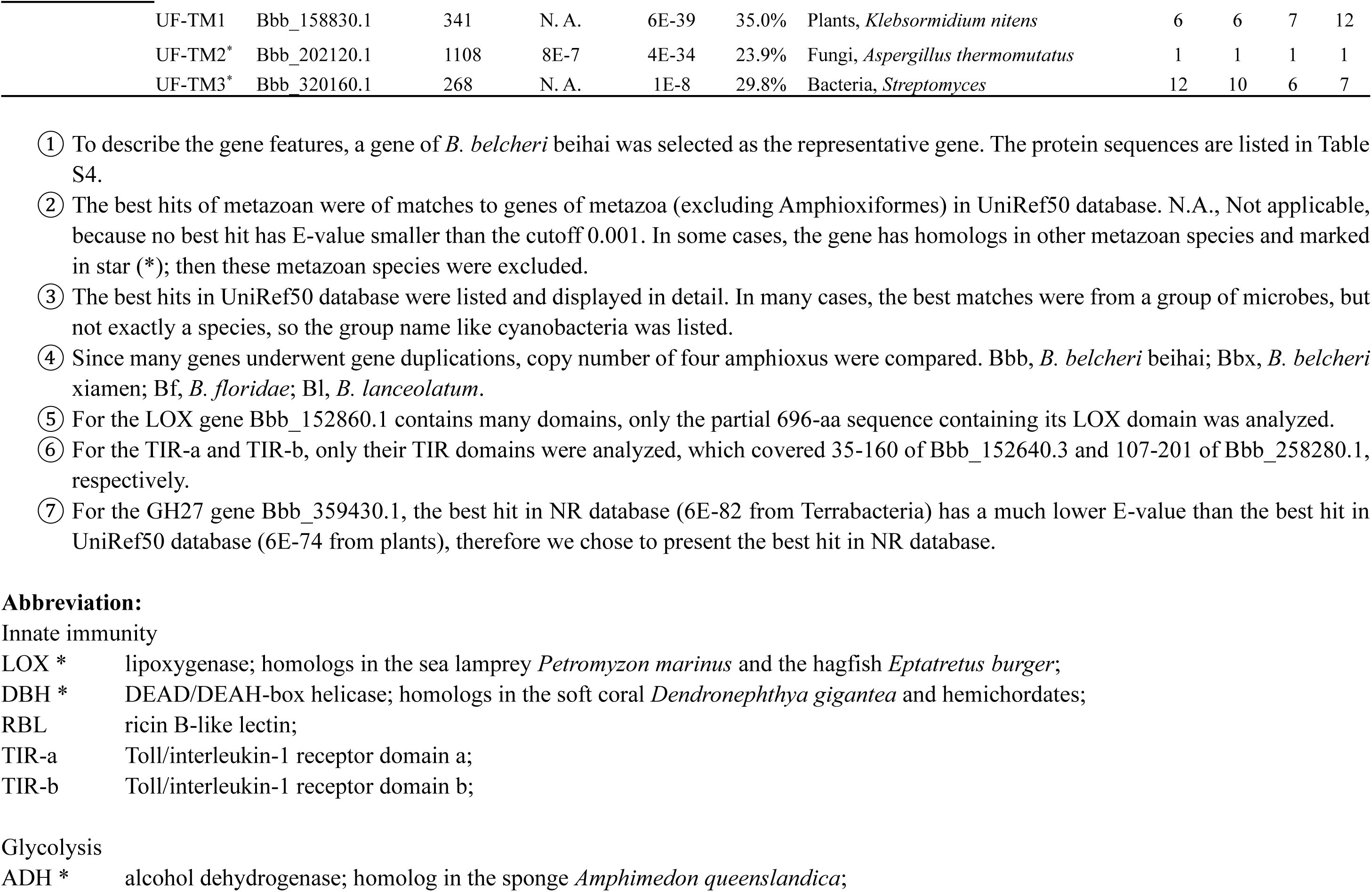
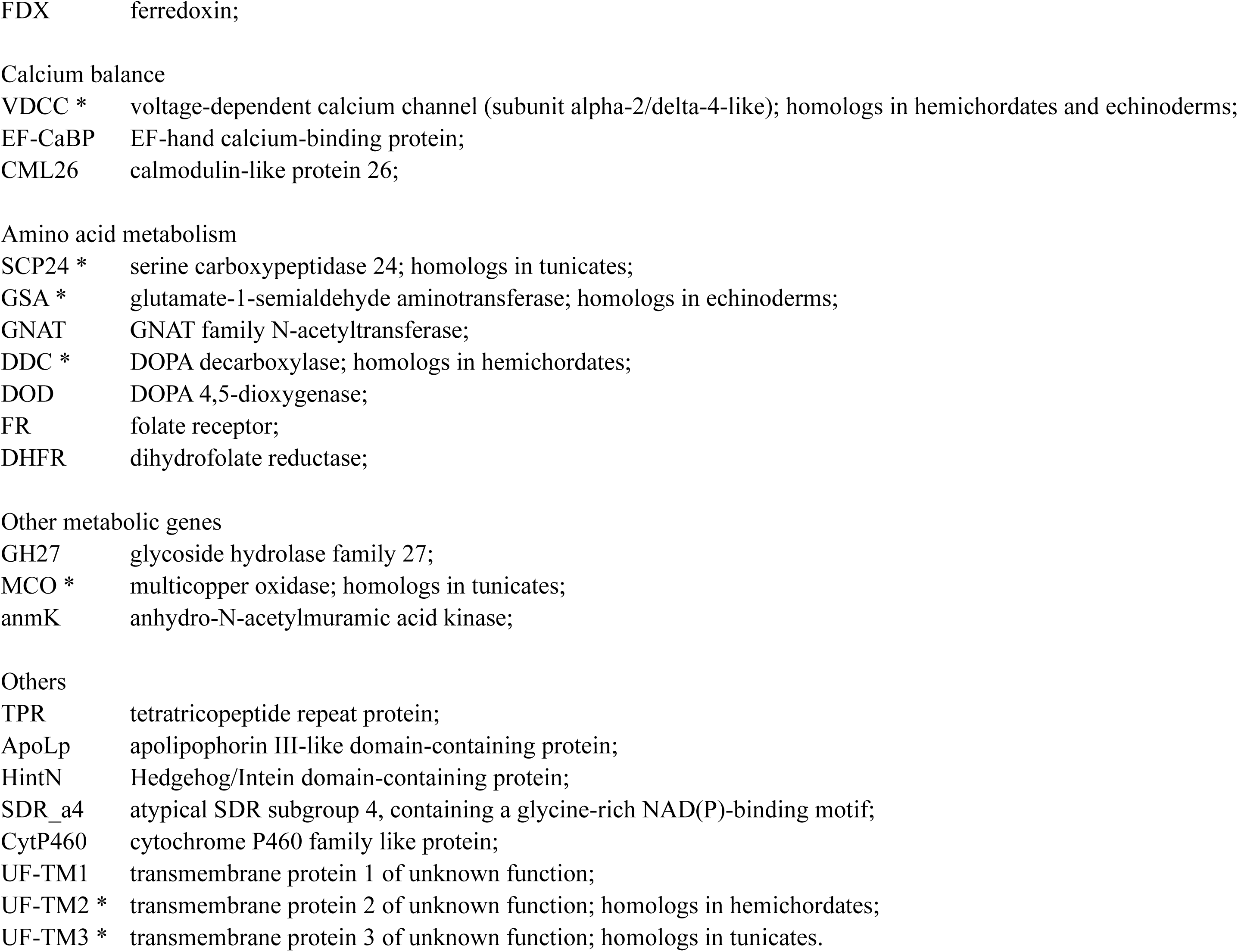
Twenty-eight groups of gene novelties identified in amphioxus.

Most candidates were filtered out and the final 28 groups of gene novelties were summarized in Table 1. Since those involved genes were conserved among up to deuterostome groups (except the VDCC gene among three groups, Table S3) and often had highly similar genes from unusual taxonomic groups (Table 1), we excluded the possibility of assembly contamination and false-positiveness as implausible. For better reproducibility, the protein sequences coded by these representative genes can be found in Table S4. Some gene novelties were further validated by phylogenetic analysis.

### Transcriptome and gene expression level

To explore the expression level of target genes, we collected the transcriptome data of different tissues of *B. lanceolatum* as follows. The SRA accessions of the transcriptome data in the NCBI database are epidermis: SRR6246024; cirri: SRR6246021; gill 1: SRR6246026; gill 2: SRR6246027; gut 1: SRR6246028; gut 2: SRR6246029; hepatic diverticulum 1: SRR6246030; hepatic diverticulum 2: SRR6246031; male gonad 1: SRR6246032; male gonad 2: SRR6246033; female gonad: SRR6246025; muscle 1: SRR6246034; muscle 2: SRR6246035; neural tube 1: SRR6246036; neural tube 2: SRR6246037. The transcript sequences of all the target genes and Glyceraldehyde 3-phosphate dehydrogenase (*GAPDH*) or elongation factor 1 alpha (*EF1A*) of *B. lanceolatum* were built as index and calculated in transcripts per million (TPM) values by the program Salmon v0.12.0 with the options ‘--type quasi -k 31’. The values of log2 (1+ TPM) appeared to be disorganized, likely due to varying transcriptome data sizes among tissues. This suggests the need for data normalization. To generate normalized TPM values, the TPM values of all target genes of *B. lanceolatum* were divided by that of the reference genes, *GAPDH* or *EF1A (Zhang et al., 2016)*, and then the log2 (1+ normalized TPM) values were calculated and presented in the heatmaps generated by the program GraphPad Prism v9.0.0.

### Comparative analysis of genes

All protein sequences of target genes were searched TBLASTN in the genomic DNA sequences and BLASTP in the annotated proteins of the four amphioxus genomes at E-value cutoff of 1E-6, to acquire all the homologs. After manual check and validation based on transcriptome data, all protein sequences of target genes were collected in four amphioxus.

The sequence alignment of genes was performed by the online tool Clustal Omega (Sievers and Higgins, 2014). The gene synteny and splice sites (locations of exon–intron boundaries) of target genes were identified and checked in the program Integrative Genomics Viewer (IGV) v2.9.4 (Thorvaldsdóttir et al., 2013). The conserved sites were defined with at least one of two flanking amino acids identical among protein sequences. The domain structure of target genes was manually drawn based on the online analysis of BLASTP with the NCBI NR database, InterProScan (Jones et al., 2014) and Pfam search (Mistry et al., 2021). Gene synteny alignment was manually drawn in PowerPoint 2021, in which the gene distance below 5 Kb defined as tandemly arrayed and below 20 Kb as proximally arrayed.

In basic phylogenetic analysis of genes, protein sequences were aligned by CLUSTAL W (Thompson et al., 1994) and MUSCLE (Edgar, 2004), and phylogenetic trees were constructed by MEGA v10.2.2 (Kumar et al., 2018) with maximum likelihood (ML) algorithm in the JTT (Jones-Taylor-Thornton) model, 80% site coverage and 100 bootstrap replicates, and then edited by online tool Interactive Tree of Life (iTOL) (Letunic and Bork, 2006).

To validate the novelty of these genes, another phylogenetic method was adopted as follows. The protein sequences of the target genes were subjected to phylogenetic analysis with the best-hit protein sequences of other taxonomical groups, including bacteria, fungi, other eukaryotes (excluding metazoa, plants, and fungi) and other metazoa (excluding Amphioxiformes), in UniRef50 database and some sequences in the NCBI GenBank database. In addition, protein sequences of the multi-copy genes of amphioxus were phylogenetically analyzed to explore their duplication and divergence. All protein sequences were aligned by MAFFT (Katoh et al., 2002) and edited in Gblocks (Castresana, 2000) with the option ‘-b4=5 -b5=h’ to generate sequence alignment of conserved amino-acid residues. Then, the sequence alignment was used to construct the phylogenetic trees in maximum likelihood algorithm by IQ-TREE v2.2.0 (Minh et al., 2020) was used with options “-seqtype AA -T AUTO --merge -rclusterf 10 -m MFP -alrt 1000 -bb 1000 -safe” to generate phylogenetic trees dually tested by Shimodaira-Hasegawa approximate likelihood-ratio test (SH-aLRT) and ultrafast bootstrap values. For poorly supported branches with SH-aLRT or ultrafast bootstrap values below 50 that may cause misinterpretation to the gene origin, the branches were discarded and manually converted into polytomies. Phylogenetic noise like long branch attraction effect (Townsend et al., 2012) was carefully excluded using the classic SAW method (Siddall and Whiting, 1999). The phylogenetic tree was edited by the online tool Interactive Tree of Life (iTOL) (Letunic and Bork, 2021).

To perform selection pressure analysis for one-to-one orthologs, we utilized the wgd pipeline (Zwaenepoel and Van de Peer, 2018) and the CodeML program from the PAML package v4.9j (Yang, 2007) to determine the nonsynonymous-to-synonymous substitution ratio (*Ka*/*Ks*). The annotated proteins of the genomes of *B. belcheri xiamen* and *B. lanceolatum* were selected for the analysis, to enable more one-to-one orthologs of the genes (Table 1).

## Data availability

The dataset of *B. belcheri* beihai supporting the results of this article is available in the NCBI BioProject repository, with the accession PRJNA804338.

## Results and Discussion

### Genome assembly and annotation

A high-quality genome of the Beihai amphioxus, *B. belcheri* beihai (Du et al., 2023; Yang et al., 2016), was *de novo* assembled, annotated, and compared with three other genome assemblies of the Xiamen amphioxus (*B. belcheri* xiamen), Florida amphioxus (*B. floridae*) and European amphioxus (*B. lanceolatum*) (Supplementary Note 1.1, Table S1, Fig. S1). For the genome assembly of *B. belcheri* beihai, the scaffold N50 length (4,185,906 bp) was second only to that of *B. floridae* (25,441,410 bp), the gap content was the lowest among the four amphioxus and the BUSCO completeness was one of the highest (97.2%) (Supplementary Note 1.2, Table S1). In the genome annotation of *B. belcheri* beihai, 44,745 protein-coding genes were annotated with 95.2% BUSCO completeness. The genome of *B. lanceolatum* was annotated with 38,190 protein-coding genes with 91.9% completeness (Supplementary Note 1.3, Table S1).

### Phylogeny and comparative genomics

Along with other genome assemblies (Supplementary Note 2.1, Table S2), a phylogenetic analysis was performed, with the sponge *Amphimedon queenslandica* used as the outgroup. The deuterostome phylogeny of this organism was consistent with current opinions (Kapli et al., 2021; Satoh et al., 2014). The phylogeny supported the monophylies of both chordates and deuterostomes, and amphioxus was suggested as the outermost or early branching chordate group (Fig. 1).

**Fig. 1.**
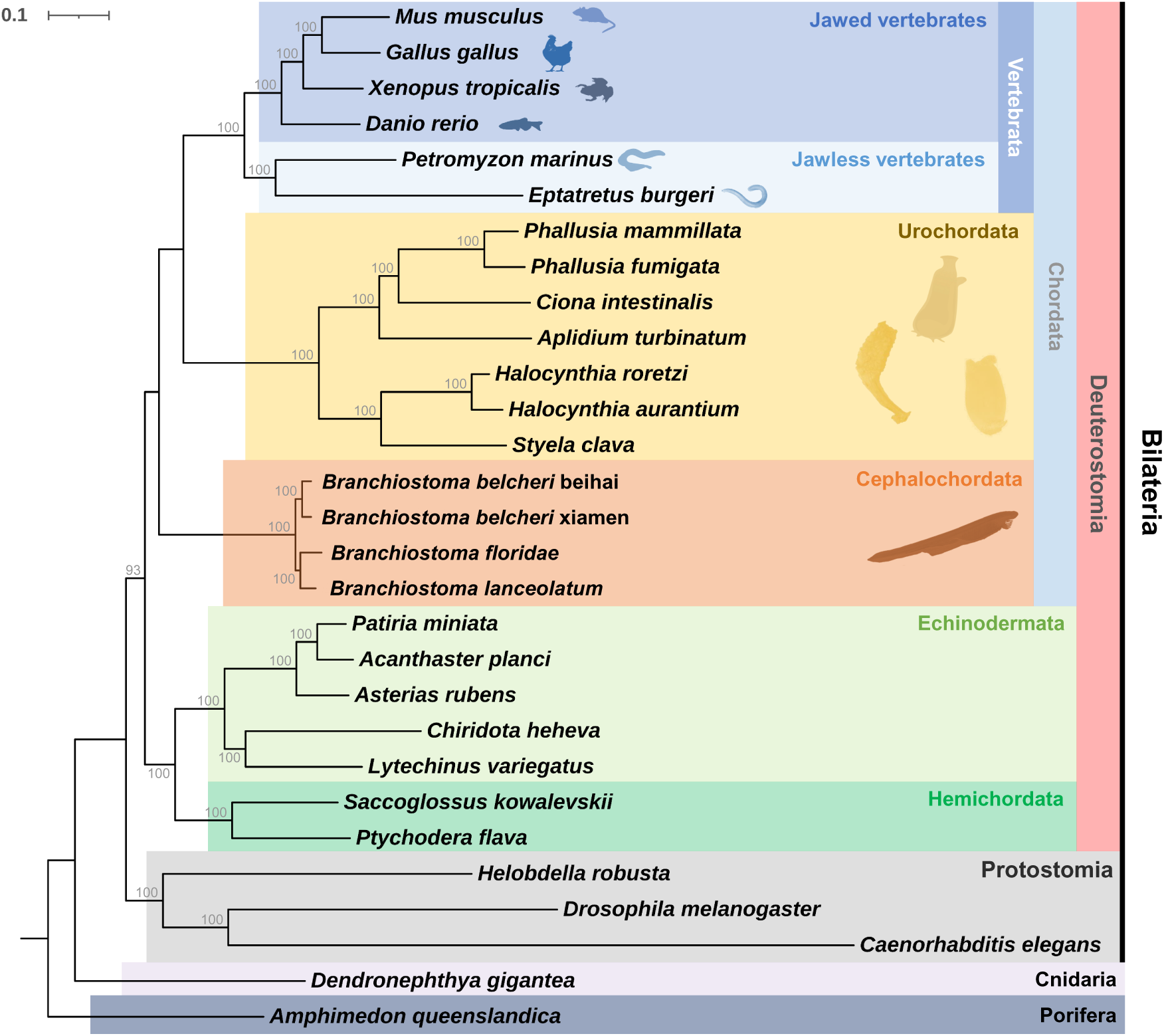
Phylogenetic analysis of major deuterostome groups. The completeness of 29 genome assemblies (Table S2) was assessed by Benchmarking Universal Single-Copy Orthologs (BUSCO) using database metazoa_odb9. 112 single-copy BUSCOs overlapped by 29 genome assemblies were extracted in protein sequences and aligned by MAFFT for the phylogenetic analysis by RAxML with 100 bootstrap replicates (Supplementary Note 2.2).

To gain insights into the genome evolution of amphioxus, we performed a comprehensive comparative genomics analysis of annotated animal genomes (Table S2). Based on the ultrametric time tree and phylogenomic orthology analysis (Supplementary Note 2.3), we observed significantly more assigned orthogroups in the four amphioxus genomes compared to other animal genomes, including over 2,000 amphioxus-specific orthogroups (Fig. S2). This observation piqued our interest in exploring the gene novelties that underlie the genome evolution of amphioxus. While it would be also interesting to explore the divergence of these amphioxus genomes, these fall outside the scope of this study, which focuses on gene novelties.

### Gene novelties

Within four annotated amphioxus genomes (Table S1), we identified a number of gene novelties conserved among them (Table 1), in which the involved genes resemble those from bacteria, fungi, plants or other eukaryotes, and markedly differ from those of other non-deuterostome metazoa (Fig. S3). In addition to cautious filtering and rigorous validation to avoid false positivity in assembled sequences, the high conservation of those genes among four independently collected, sequenced, and assembled genomes of amphioxus (Table 1) precluded the occurrence of contamination. Previous studies have identified a number of gene novelties in stem chordates (Holland et al., 2008) and deuterostomes (Simakov et al., 2015). To avoid repeating these efforts and to focus on novel gene features in amphioxus, we excluded candidate novelties that are also conserved in jawed vertebrates, including the sialidase gene, which may have arisen in the deuterostome stem (Fig. S4, Supplementary Note 3.1). In the scope of this study, we also excluded novel genes that may originate from extreme sequence variation, protein domain recombination or shuffling events.

A gene novelty could generate many novel genes via gene duplications; for example, in *B. belcheri* beihai, 28 groups of gene novelties generated 108 new genes (Table 1). According to their biological functions, these involved genes were classified into six categories, namely, innate immunity, glycolysis, calcium balance, amino acid metabolism, other metabolic genes, and others without explicit functional domains (Table 1). The expression level of these genes was analyzed in different tissues of *B. lanceolatum* because of its high-quality transcriptome data, and all the genes were expressed in at least one tissue, although some gene expression levels are quite low (Fig. S5). Regardless of the inconsistent gene expression in tissue duplicates of muscle and male gonads, the most marked expression feature was that the genes coding for calcium-binding proteins were highly expressed in the guts and hepatic diverticula (Fig. S5, Supplementary Note 3.2). Subsequently, we explored these novel genes in three important functional groups, namely innate immunity, glycolysis, and calcium balance. More gene novelties were discussed in Supplementary Note 3.

### Innate immunity

Innate immunity is the first-line nonspecific immune system against infections, and this immune system is observed in jawed and jawless vertebrates as well as invertebrates (Bartl et al., 2003). Successful immunity evolution confers survival traits to a species. In the absence of an adaptative immune system in amphioxus (Bartl et al., 2003), innate immunity is the most important weapon of amphioxus in response to various pathogens.

Lipoxygenases (LOXs) are important in arachidonic acid metabolism, participate in inflammatory lipid signaling and further regulate inflammatory and allergic responses (Yuan and Xu, 2016; Yuan et al., 2014). A group of duplicated LOX genes of amphioxus were identified similar to those of cyanobacteria or proteobacteria (Table 1). Notably, these LOX genes have unexpected homologs from only two living groups of jawless vertebrates (Cyclostomata) (Janvier, 2006), the sea lamprey *Petromyzon marinus* and the hagfish *Eptatretus burger* (Fig. S6); however, such homologs are not observed in other metazoa. In the phylogenetic tree, these LOX genes of amphioxus are close to those of sea lamprey, bacteria and other eukaryotes (Fig. 2A). Three splice sites (locations of exon–intron boundaries) were conserved among the LOX genes of four amphioxus and the sea lamprey (Fig. S6). In the absence of genome annotation, two fragments of the LOX gene were recovered from the hagfish genome, and three other conserved splice sites were identified between those of the two jawless vertebrates (Fig. S6).

**Fig. 2.**
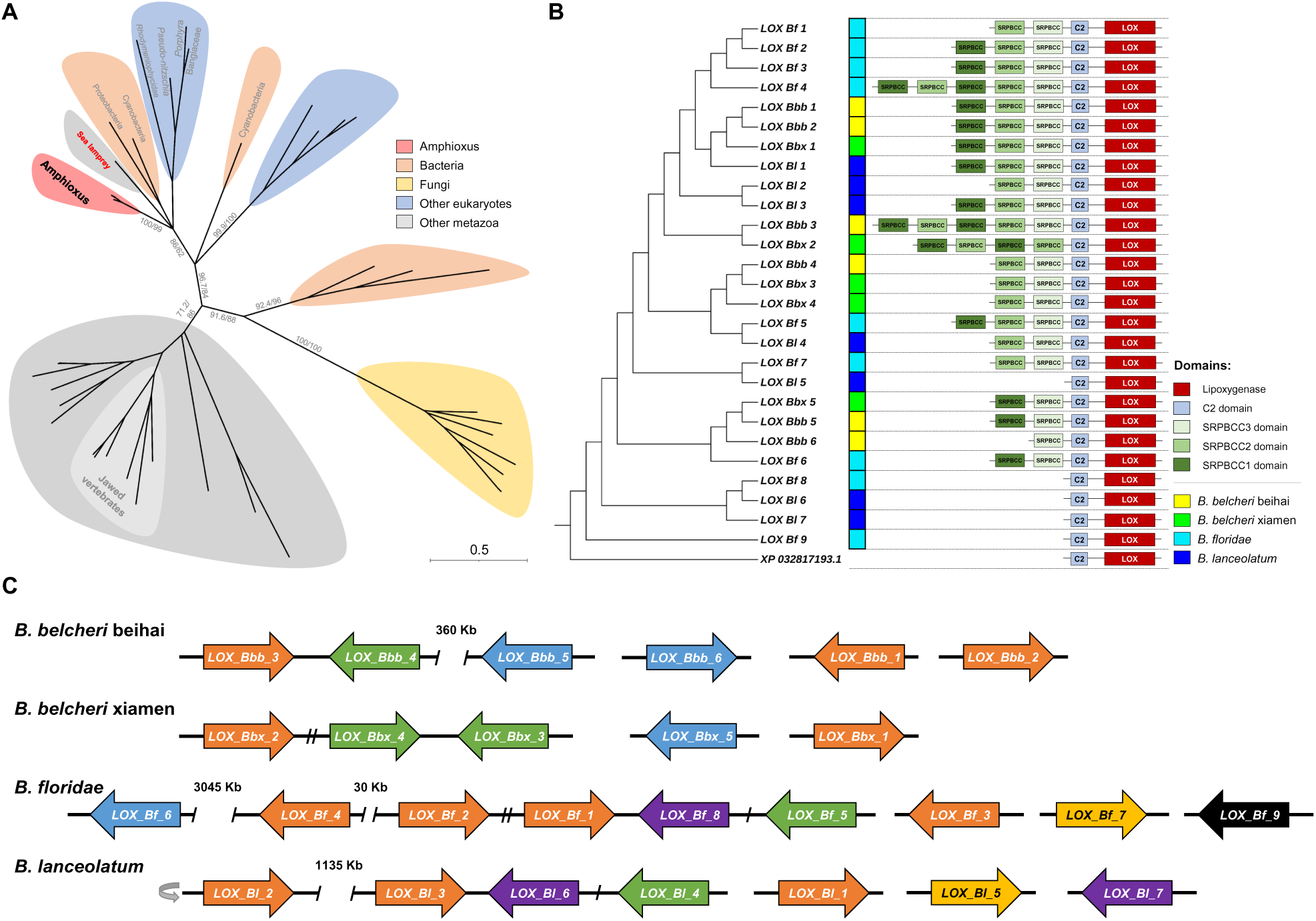
A group of lipoxygenase (LOX) genes in amphioxus. **(A)** The amphioxus LOX genes and their closest genes in the UniRef50 database from other taxonomic categories, including bacteria, fungi, other eukaryotes (excluding metazoa, plants, and fungi) and other metazoa (excluding Amphioxiformes) were collected for phylogenetic analysis. In this maximum likelihood phylogenomic tree, SH-aLRT and ultrafast bootstrap support values are given in that order on the branches. **(B)** Phylogenetic analysis of all LOX genes of four amphioxus and their domain architecture. The LOX gene of the sea lamprey *Petromyzon marinus* (GenBank accession: XP_032817193.1) was used as outgroup. C2, C2-like domain (universal calcium-binding domain); SRPBCC, PYR/PYL/RCAR family domain in SRPBCC superfamily. **(C)** Gene synteny alignment of the LOX genes of four amphioxus. The distance precision was set as 5 Kb, e.g., 30 Kb represents for 30-35 Kb. If not noted, a slash (/) between two genes represents 5 Kb, e.g., one slash represents 5-10 Kb. Except the outlier (*LOX_Bf_9*), LOXs located on clades supported by high bootstrap value (>95) in the phylogenetic tree were painted in the same color. The suffixes stand for four amphioxus, *Bbb* for *B. belcheri* beihai, *Bbx* for *B. beicheri* xiamen, *Bf* for *B. floridae* and *Bl* for *B. lanceolatum*. The grey turnover arrow means the gene alignment is after reverse complement.

The phylogenetic analysis revealed that these LOX genes were duplicated and diversified among the four amphioxus and *LOX_Bf_9* was an outlier gene (Fig. 2B). All the LOXs contain a C2-like domain (Fig. 2B). Except for the five LOXs of *B. floridae* and *B. lanceolatum*, the LOXs possessed one to five domains of the PYR (pyrabactin resistance)/PYL (PYR1-like)/RCAR (regulatory components of ABA receptor) family in the SRPBCC superfamily (SRPBCC domains, Fig. 2B) and these SRPBCC domains could be further divided into three groups (Fig. S7). In Fig. 2C, the gene synteny alignment revealed that many LOXs (12 of 27 genes) were proximally located (distance <20 Kb) and possibly generated by tandem gene duplications. *LOX_Bl_5* had an apparently wide expression in the gills, gut, and hepatic diverticula (Fig. S5).

These unusual LOX genes containing a C2-like domain were reported in amphioxus (Yuan and Xu, 2016; Yuan et al., 2014) and first in jawless vertebrates, although the SRPBCC domains were not identified. Since SRPBCC superfamily proteins contain a deep hydrophobic ligand-binding pocket and can bind diverse ligands (Chen et al., 2015), we consider that these SRPBCC domains may also function as receptor domains in these LOX genes in response to various signals. A multifunctional LOX containing a SRPBCC domain was reported in the red alga *Pyropia haitanensis* and presented substrate flexibility, oxygenation position flexibility, and functional versatility (Chen et al., 2015). These LOX genes may enrich the inflammatory lipid signaling pathway of amphioxus.

DEAD-box helicases (DBHs) are known as sensors for nucleic acids and mediators of antiviral innate immunity (Taschuk and Cherry, 2020). A DBH gene was identified as a single-copy gene that resembles those from bacteria, and it was conserved among four amphioxus (Table 1). In the phylogenetic tree (Fig. 3A), this DBH gene of amphioxus is close to those of the charophyte alga *Klebsormidium* (*K.*) *nitens*, soft coral *Dendronephthya gigantea*, hemichordate *Saccoglossus kowalevskii*, and a group of bacteria. Intriguingly, the DBH genes of amphioxus shared three conserved splice sites with those of charophyte alga, soft coral, and hemichordate (Fig. 3B and S7), which suggested that the DBH genes of four amphioxus, soft coral and hemichordate shared a common source with that of the charophyte alga. A homologous DBH gene was also identified in the genome of the other hemichordate, *Ptychodera flava*, in the absence of annotation. This DBH gene was widely expressed at low levels in the tissues of *B. lanceolatum* (Fig. S5). The single-copy DBH gene may serve as a novel sensor in the antiviral innate immunity of amphioxus. Another DBH gene was reported as a horizontally transferred gene in coral species (Bhattacharya et al., 2016).

**Fig. 3.**
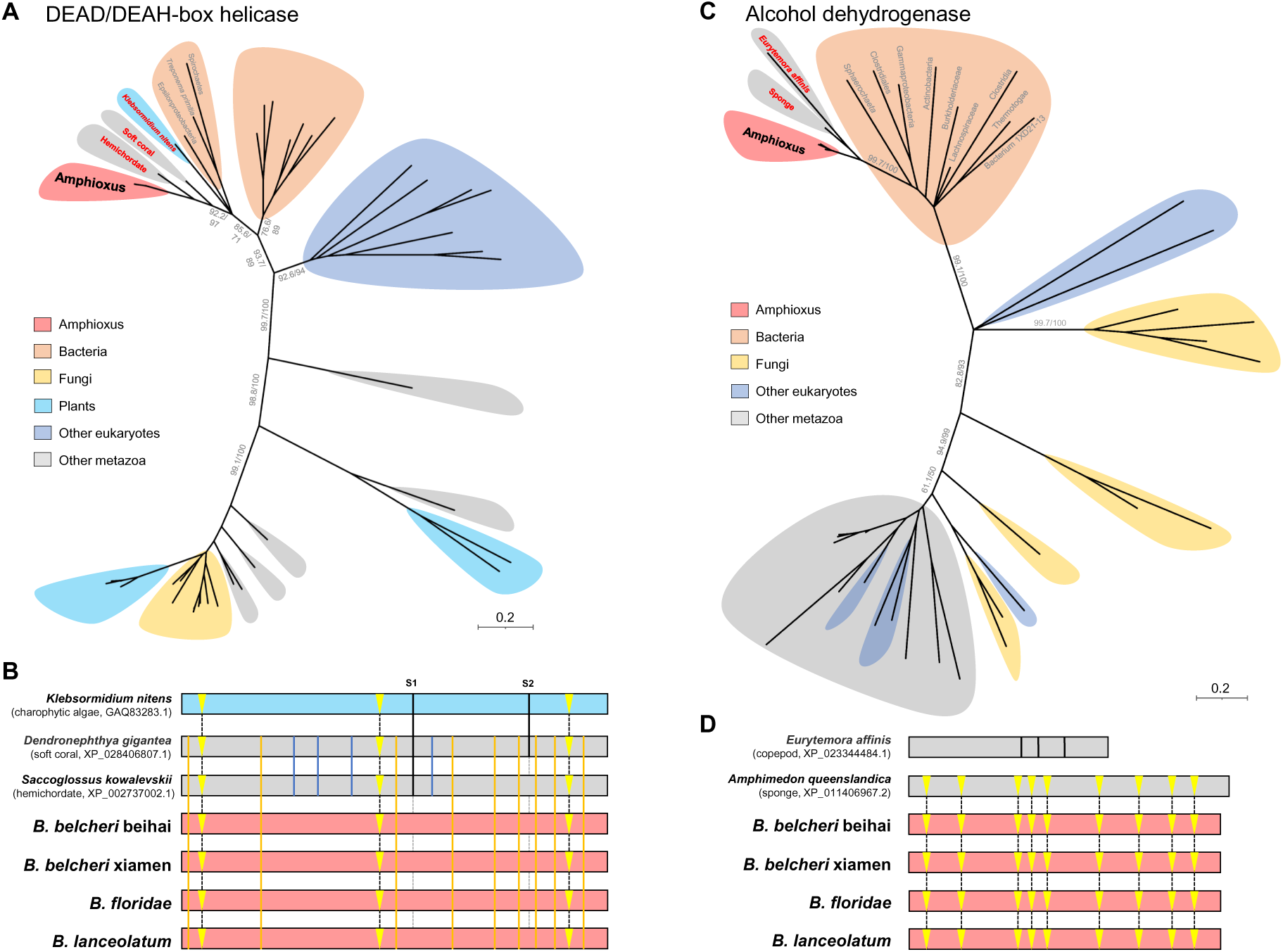
Comparative analysis of two gene novelties in amphioxus. **(A)** The DEAD/DEAH-box helicase (DBH) genes of amphioxus and their closest genes in the UniRef50 database from other taxonomic categories, including bacteria, fungi, other eukaryotes (excluding metazoa, plants, and fungi) and other metazoa (excluding Amphioxiformes) were collected for phylogenetic analysis. **(B)** Alignment of splice sites of the DBH genes of amphioxus, the charophytic algae *K. nitens* (GenBank accession: GAQ83283.1), the soft coral *Dendronephthya gigantea* (GenBank accession: XP_028406807.1) and the hemichordate *Saccoglossus kowalevskii* (GenBank accession: XP_002737002.1). Conserved splice sites were marked by vertical lines (Fig. S8), and three conserved splice sites of all seven sequences were highlighted in yellow triangles. Two unusual splice sites were labeled as S1 and S2. **(C)** The alcohol dehydrogenase (ADH) genes of amphioxus and their closest genes in the UniRef50 database from other taxonomic categories were collected for phylogenetic analysis. **(D)** Alignment of splice sites of the ADH genes of amphioxus, the sponge *Amphimedon queenslandica* (GenBank accession: XP_011406967.2) and the copepod *Eurytemora affinis* (UniProt accession: UPI000C762174) (Fig. S11). In these maximum likelihood phylogenomic trees, SH-aLRT and ultrafast bootstrap support values are given in that order on the branches.

In addition, another nine conserved splice sites in orange lines (Fig. 3B and S7) indicated close phylogenetic relationships among the DBHs of the four amphioxus, soft coral and hemichordate; another four conserved splice sites in blue lines (Fig. 3B and S7) suggested that the DBHs of the soft coral and hemichordate originated from the same ancestral gene, while two unusually conserved splice sites marked with black lines (S1 and S2, Fig. 3B and S7) could be explained by intron loss during gene evolution, which indicated S1 loss in ancestral amphioxus but S2 loss in both the ancestral amphioxus and hemichordate. Additional gene novelties related to innate immunity, such as Ricin B-like lectins, can be found in Supplementary Note 3.3.

### Glycolysis

Two novel genes, Alcohol Dehydrogenases (ADHs) and Ferredoxin (FDX), have been identified as being involved in the glycolysis pathway of amphioxus (Table 1). ADHs facilitate the oxidation of alcohols into aldehydes and ketones during the glycolysis process (Di et al., 2021). In the phylogenetic tree (Fig. 3C), an ADH gene containing zinc-binding sites and belonging to the conserved medium-chain dehydrogenase/reductase (MDR) family was much closer to those of the sponge *Amphimedon queenslandica*, the copepod *Eurytemora affinis*, and a bacterial group than to those of other metazoa. The splice sites of the ADH gene of amphioxus are highly conserved with those of the sponge but not the copepod (Fig. 3D and S10), which suggested that the ADH genes of amphioxus and *Amphimedon queenslandica* have the same eukaryotic origin. Among the 100 best hits of this ADH gene in the NCBI NR database, we noticed over ten hits from anaerobic bacteria and the ADH gene of *B. belcheri* beihai shared 64% identity with that of *Anaerolineae bacterium* (GenBank accession: MCD6286762.1).

FDX, which contains an iron-sulfur cluster, is recognized as a mediator of electron transfer in numerous metabolic pathways, including glycolysis (Meyer, 1988). The FDX gene of amphioxus was identified to be highly conserved with the genes in certain plants, bacteria, or other eukaryotes (identities over 68%) but not with that of other metazoa (excluding Amphioxiformes, Table 1). This FDX gene was tandemly duplicated into two copies on opposite strands in *B. floridae*. All these FDX genes contain a Ubl_FUBI (Ubiquitin-like protein FUBI) domain, which may protect the FDX gene from ubiquitination and subsequent degradation (Rossman et al., 2003).

### Calcium balance

Calcium balance and osmoregulation of fishes are important for their physiological functions (Guerreiro, 2007). A voltage-dependent calcium channel (VDCC), EF-hand calcium-binding protein (EF-CaBP) group and calcium-binding protein (CML26) were identified as gene novelties in amphioxus (Table 1).

VDCC is a complex of several different subunits (Dolphin, 2006); however, whether this VDCC gene functions independently as a calcium channel deserves further investigation. The VDCC genes of amphioxus were more similar to those of other eukaryotes than to those of other metazoans (Table 1). Additionally, the VDCC genes of amphioxus have homologs in the hemichordate *Saccoglossus kowalevskii* and three starfish species, all of which share three conserved splice sites (Fig. S12). This VDCC gene was proposed to be related to electrophysiological function and was widely expressed among tissues, but no apparent tissue-specific expression (Fig. S5).

EF-CaBPs may function in maintaining a high calcium concentration in the endoplasmic reticulum (Camacho et al., 2003) and could be involved in bone formation (Allen and Hall, 1978). Of the 73 newly identified EF-CaBP genes in four amphioxus, the bit scores of the best hits to bacteria were significantly higher than those to other metazoa (Fig. 4A and S12).

**Fig. 4.**
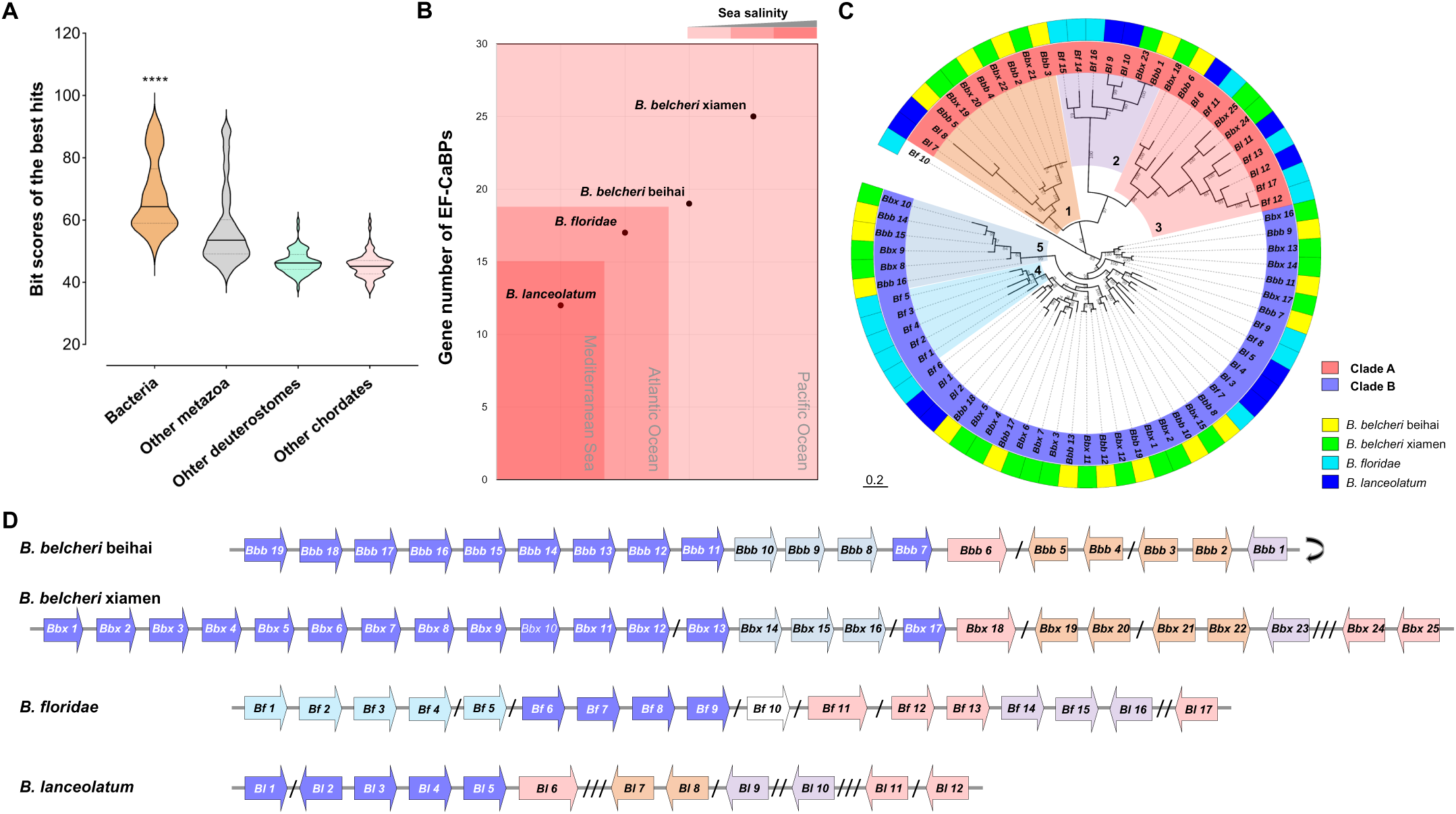
A group of proximally arrayed EF-hand calcium-binding proteins (EF-CaBPs) **(A)** Of a group of EF-CaBP genes of amphioxus, bit scores of the best hits in the UniRef50 database to other metazoa, other deuterostomes and other chordates (excluding Amphioxiformes respectively) were compared with those to bacteria in violin plot with quartile lines. **** *p* <0.0001, when comparing the best hits to bacteria with other three groups, respectively. **(B)** Correlation of the gene number and the sea surface salinity of collection locations of four amphioxus. A darker color indicates higher sea salinity. **(C)** Phylogenetic analysis of these EF-CaBPs of four amphioxus. Two large clades, Clade A and B, were highlighted and five specific clusters were marked. **(D)** Gene synteny alignment of the proximally arrayed EF-CaBP genes. A slash (/) represents 5 Kb, e.g., one slash represents 5-10 Kb, and three slashes represent 15-20 Kb, while none represents below 5 Kb. The grey turnover arrow means after reverse complement.

This group of EF-CaBP genes was proximally arrayed in four amphioxus and presented high copy number variation. *B. belcheri* xiamen has 25 EF-CaBP genes, *B. belcheri* beihai has 19 genes, *B. floridae* has 17 genes, and *B. lanceolatum* has only 12 genes. Marine salinity is linearly correlated with the calcium ion concentration (He et al., 2020). Therefore, we further explored the relationship of the gene number variation of these proximally arrayed EF-CaBP genes and the sea surface salinity of the collection locations (Fig. 4B). The Mediterranean Sea has a much higher than average seawater salinity (approximately 35 parts per thousand) (Millero et al., 2008; Millot and Taupier-Letage, 2005), and the Atlantic Ocean is globally saltier than the Pacific Ocean (Jones and Cessi, 2017). Although high-quality salinity data for the nearshore regions where the four amphioxus were collected is lacking, a negative correlation between seawater salinity and the number of EF-CaBP genes can be observed. *B. lanceolatum*, collected from the Mediterranean Sea, has the fewest genes, while the two *B. belcheri* from the Pacific Ocean have a higher number of genes (Fig. 4B).

In the phylogenetic tree (Fig. 4C), except for the outlier gene *Bf_10*, the EF-CaBP genes were classified into two large groups, Clade A and Clade B. Clade A was further classified into three clusters, Cluster 1, Cluster 2, and Cluster 3. In Cluster 1, no genes of *B. floridae* were identified while eight genes of two *B. belcheri* were assigned to four pairs, and they included four highly conserved genes, *Bbb_2*, *Bbb_3*, *Bbx_21* and *Bbx_22*. However, three and two tandemly arrayed genes in Cluster 2 were identified in *B. floridae* and *B. lanceolatum*, respectively, and variable duplications were observed in Cluster 3. The phylogenetic analysis indicated that at least three genes in Clade A appeared in the ancestral amphioxus species. For Clade B, five relatively unique genes of *B. floridae*, *Bf_1-5* in Cluster 4, were proximally arrayed, and many paired genes were identified in two *B. belcheri*, such as three pairs in Cluster 5. All 73 EF-CaBPs were proximally arrayed in four amphioxus (Fig. 4D) and all spaces between genes were below 20 Kb. Notably, four longer genes, *Bbb_6*, *Bbx_18*, *Bf_11* and *Bl_6*, contained more EF-hand motifs (Cluster 3, Fig. 4C).

Except for the large group of proximally arrayed EF-CaBPs above, the CML26 gene was a calcium-binding gene with a shorter length (Table 1). This CML26 gene was tandemly duplicated in *B. floridae* and *B. lanceolatum* (Table 1). Initially, we considered the CML26 gene (Table 1) to belong to another event, but the conserved splice sites (Fig. S14) suggested that the CML26 gene and the large group of EF-CaBPs possibly originated from a common ancestor.

Along with the two CML26 genes, these EF-CaBP genes were apparently highly expressed in the gut and hepatic diverticulum, which are both considered digestive organs for calcium absorption (Fig. S5). Thus, the expansion of these EF-CaBP genes was proposed to occur in response to marine salinity and calcium concentration changes. The *EF-CaBP_Bl_11* and the two CML26 genes had the highest expression levels (Fig. S5).

## Conclusions

As an important group of invertebrate chordates, amphioxus play a pivotal role in the evolutionary study of cephalochordates, chordates, and deuterostomes. In this study, the genome of the Beihai amphioxus, *B. belcheri* beihai, was *de novo* assembled and annotated (Table S1). Combined with the three amphioxus genomes (Table S1), we performed a phylogenetic analysis of the deuterostome genomes (Fig. 1), which supported the monophylies of deuterostomes and chordates.

We identified a number of gene novelties conserved among the four amphioxus genomes (Table 1, Supplementary Note 3.1). Among these gene novelties, except for the TIR domains of amphioxus, which was mentioned in a previous report (Huang et al., 2008), all other genes were newly identified and reported. Certain involved genes had homologs from other deuterostome groups except for the jawed vertebrate group (Table 1 and S3). That is, the LOX gene had homologs from two jawless vertebrates; 8 genes had homologs from either tunicates or echinoderms or hemichordates; and only the VDCC gene had homologs from both echinoderms and hemichordates. The conservation of these genes can be well explained by frequent gene gain and loss in the early and rapid evolution of deuterostomes (Table S3) (Kapli et al., 2021). For example, the LOX gene was suggested to be a novel gene that was introduced at the deuterostome stem but decayed in tunicates and jawed vertebrates. Those gene novelties could be well dated according to their conservation among different deuterostome groups, such as the three genes conserved with urochordates (SCP24, MCO and UF-TM3) were proposed as results of events occurred at the chordate stem, while their absence in vertebrates is due to gene decay. While it is true that certain genes, such as DBH and ADH, have homologs in ancient metazoa like coral or sponge, we cannot definitively conclude that these genes originated in the common ancestor of all metazoa. This is because the homology was observed only in specific coral or sponge species, suggesting independent acquisition events rather than inheritance from a common ancestor. For example, while the alignment of splice sites within the DBH gene (Fig. 3B) may suggest a close relationship among these genes, it cannot support that this gene was introduced in a common ancestor before the emergence of the ancient soft coral. Although these genes with restricted distribution are not exclusive to the amphioxus lineage, their surprisingly high similarity to non-metazoan genes is interesting and deserves further investigation. Additionally, the absence of high-quality genomes from the other amphioxus genera, including *Epigonichthys* and *Asymmetron*, hinders our comprehensive investigation of more cephalochordate species.

The gene novelties without any homologs in deuterostomes were suggested to be acquired by amphioxus lineage-specific horizontal gene transfer (HGT) from non-metazoan species. A number of adaptation innovations were enabled by these genes, such as innate immunity-related events that contributed more complexity to the immune system in amphioxus, especially for amphioxus has no adaptative immune system. Although the function of these novel genes has not been validated in any experimental model, either the gene duplications (Table 1) or the negative selection indicated by their low *Ka*/*Ks* values (Fig. S25) suggest that they may indeed be functional essential genes.

### Proposed mechanisms

Although it is difficult to rule out gene loss in other groups like protostomes and/or extreme divergence, those gene novelties, especially the genes with phylogenetic supports, can be explained as results of HGT from cross-kingdom origins such as marine bacteria, considering many genes shared high similarity with those from unusual taxonomic groups. Notably, reports have documented similar HGT events of the RAG transposon and the F-type lectins in amphioxus (Bishnoi et al., 2015; Carmona and Schatz, 2017).

The HGT of amphioxus could be explained by the ‘you are what you eat hypothesis’ (Doolittle, 1998). This hypothesis suggests that the HGT genes in amphioxus originate from its food sources, which is particularly plausible considering that amphioxus can perform phagocytic intercellular digestion in the diverticulum lumen (He et al., 2018). All modern deuterostomes were considered to have evolved from a filter-feeding ancestor (Simakov et al., 2015). Along with other deuterostomes, such as tunicates, echinoderms, and hemichordates, amphioxus is known as a filter feeder (Peterson and Eernisse, 2016; Riisgård and Svane, 1999) and may obtain genes from marine microbes, including bacteria and algae, which are potential foods. For example, the charophyte alga *K. nitens* frequently appeared in the analysis of the HGT genes, including the DBH, SCP24 and UF-TM1 genes (Table 1). The DBH gene was unusually identified in the charophyte alga *K. nitens* (Fig. 3A). The highly conserved splice sites among the DBH genes of the charophyte algae, soft coral, hemichordate, and amphioxus (Fig. 3B and S7) suggested that their DBH genes originated from a common ancient eukaryote, such as some marine algae. Although the charophyte alga *K. nitens* is considered a freshwater/terrestrial and nonmarine species (Hori et al., 2014), we proposed that the ancestor of *K. nitens* was possibly distributed in marine environments and served as food for these marine animals, especially because of the sister marine species *K. marinum* (no available genome sequences). Similarly, the SCP24 gene (Fig. S15) and UF-TM1 gene (Table 1) possibly originated from the marine ancestor of *K. nitens*.

### New gene generation

The origin of new genes is a fundamental process of genome evolution (Long et al., 2003). In this study, the comparative analysis of the gene novelties revealed insights into the new gene generation. Regardless of whether common gene duplication and diversification occurred, many genes of amphioxus underwent possible gene fusion or fission to generate new genes. Three novel models of new gene generation were summarized, including gene fusion (Fig. S26A), gene fission (Fig. S26B) and partial duplication or deletion models (Fig. S26C).

In the gene fusion model (Fig. S26A), we revealed that two distinct TIR domains fused with a number of endogenous genes (or domains) with immune functions (Fig. S10) and generated a series of new immune genes. Both gene fission and partial duplication or deletion were observed in the LOX family of amphioxus (Fig. 2B). Since we found that the LOX genes of amphioxus were mostly similar to bacterial LOX genes containing up to three SRPBCC domains and three groups of SRPBCC domains could be clustered with three SRPBCC domains of the cyanobacterium *Coleofasciculus sp. FACHB-125* (GenBank accession: WP_190414990.1) (Fig. S7), we considered that the original LOX gene of amphioxus contains three SRPBCC domains. Then, they were split into LOX genes with only C2 domains and genes with only SRPBCC domain via gene fission (Fig. S26B) and generated a series of LOX genes with variable numbers of SRPBCC domains (Fig. 2B) via partial gene duplication or deletion (Fig. S26C). The investigation of the origins of the functional domains of these novel genes through phylogenetic analysis would enhance the credibility of the findings. However, the substantial challenges posed by the relatively low sequence similarities resulting from extensive evolutionary divergence, coupled with the lack of comparable outgroup genes from other taxonomic groups, complicate this analysis significantly.

### Limitations of this study

During the identification of gene novelties, we ignored those candidates with homologous genes in jawed vertebrates. Although this avoided repeating efforts on the gene novelties of chordates (Holland et al., 2008) and deuterostomes (Simakov et al., 2015), we may miss some important genes. Meanwhile, the filter standard of gene novelties may be too high to achieve highly comprehensive results, and many genes may be missed during filtration.

The lack of animal model experiments impeded our validation of the metabolic functions of some interesting genes. The exploration of genomic evolution following these gene novelties presents an intriguing direction, except the gene duplications, such as LOXs and EF-CaBPs. To fully explore this direction, delineating the functional pathways and gene networks in amphioxus would be necessary, but represents a substantial challenge that falls beyond the scope of this study. In the case of the ADH gene, three intrinsic ADH like genes in *B. lanceolatum*, which share homology with those in other metazoan species, were identified with high similarity to the novel ADH gene, exhibiting low E-values below 1E-30 in BLASTP. In muscle tissue, the novel ADH gene could be expressed at levels over the combined expression of the three intrinsic genes, indicating its functional importance and warranting further investigation into its role within the related gene network.

Despite the methodological challenges and difficulties in pinpointing the precise origins of gene novelties, the phylogenetic trees support our conclusion that these notable genes are more closely related to those of non-metazoan species and represent gene novelties in amphioxus. However, the analysis included only a limited number of genes, as many were excluded due to relatively low similarities resulting from long-term evolution or the absence of comparable outgroup genes from other taxonomic groups.

## Appendix A. Supplementary Materials

Supplementary Notes 1–3 Tables S1–4

Figures S1–26

## Acknowledgements

This work was supported by the General Research Fund from Research Grants Council (Reference numbers: 464710, 475113, 14119219, 14119420, 14175617) and Health and Medical Research Fund from Food and Health Bureau (Reference numbers: 06171016, 07181266) of Hong Kong. We would like to thank Prof. Jun-Yuan Chen from the State Key Laboratory of Palaeobiology and Stratigraphy, China, for providing the Beihai amphioxus samples and for reviewing our manuscript. Additionally, we extend our gratitude to Dr. Laura Jane Falkenberg from the Simon FS Li Marine Science Laboratory, The Chinese University of Hong Kong, Hong Kong, for her valuable comments on the adaptation of amphioxus to ocean environments.

