## Supplemental Materials for "Gene Novelties in Amphioxus Illuminate the Early Evolution of Cephalochordates"

for

**This PDF file includes:**

Supplementary Notes 1–3

Tables S1–4

Figures S1–26

References

### Contents


Supplementary Note 1. Genome assembly and annotation 1

*1.1 Four amphioxus genomes* 1

*1.2 Genome assembly* 1

*1.3 Genome annotation* 2

Supplementary Note 2. Deuterostome phylogeny and comparative genomics 4

*2.1 Collection of genome assemblies* 4

*2.2 Phylogenetic analysis* 4

*2.3 Comparative genomics* 4

Supplementary Note 3. Gene novelties in amphioxus 6

*3.1 Brief summary of gene novelties* 6

*3.2 Gene expression in amphioxus tissues* 6

*3.3 Other gene novelties related to innate immunity* 7

*3.4 Gene novelties related to amino acid metabolism* 8

*3.5 Gene novelties related to other metabolic functions* 10

*3.6 Gene novelties without apparent functional domains* 11

Supplementary tables 13

Supplementary figures 18

References 44

#### Supplementary Note 1. Genome assembly and annotation

##### *1.1 Four amphioxus genomes*

In this study, four amphioxus genomes were combined for phylogenomic analysis, including two subspecies of *B. belcheri*, the Florida amphioxus (*B. floridae*) and the European amphioxus (*B. lanceolatum*). The Beihai amphioxus and the Xiamen amphioxus are suggested as two *B. belcheri* subspecies and tentatively named as *B. belcheri* beihai and *B. belcheri* xiamen respectively.

The Beihai amphioxus, *B. belcheri* beihai, were obtained from the sea near Dianbai District, Maoming City, Guangdong Province, China, while the Xiamen amphioxus, *B. belcheri* xiamen was collected from Xiamen Bay, Fujian Province, China (Huang et al., 2014). The Florida amphioxus, *B. floridae* was collected from Old Tampa Bay, Florida, USA (Putnam et al., 2008) and the European amphioxus, *B. lanceolatum* was collected in western Mediterranean (Banyuls-sur-mer, Pyrénées-Orientales, France) (Marlétaz et al., 2018).

##### *1.2 Genome assembly*

The genomes of the Xiamen amphioxus, the Florida amphioxus and the European amphioxus were directly downloaded from NCBI GenBank accessions: GCA_001625305.1, GCA_000003815.2 and GCA_900088365.1, respectively.

The adult bodies of the Beihai amphioxus, *B. belcheri* beihai were used for genomic DNA extraction and then sequenced by Illumina HiSeq 2000 in 500-bp and 3,000-bp libraries to generate paired-end NGS reads and PacBio RS system for third-generation long reads. To obtain a high-quality genome assembly with the best continuity, different assembly methods were tried. Finally, the genome assembly was first constructed with the long reads using Canu and the short reads were used for scaffolding and polishing. The *de novo* assembled genome of *B. belcheri* beihai was deposited under the NCBI GenBank accession GCA_025402995.1.

The genome assembly and annotation were summarized in Table S1. The genome assembly sizes of four amphioxus ranged from 426,123,519 bp of *B. belcheri* xiamen to 513,460,931 bp of *B. floridae*.

The scaffold numbers ranged from 432 of *B. floridae* to 10,247 of *B. lanceolatum*, and the scaffold N50 lengths ranged from 1,295,816 bp of *B. lanceolatum* to 25,441,410 bp of *B. floridae*. The superlong scaffold N50 length of *B. floridae* genome was because HiRiSE scaffolding (Koch, 2016) was involved in the genome assembly.

The gap contents of *B. floridae* and *B. lanceolatum* were as high as 5.128% and 4.120% respectively, while those of *B. belcheri* beihai and *B. belcheri* xiamen were as low as 0.156% and 1.297% respectively. Regardless of the gaps, the ungapped lengths of *B. belcheri* beihai, *B. floridae* and *B. lanceolatum* ranged from 474,945,525 bp of *B. lanceolatum* to 487,128,935 bp of *B. floridae*, while that of *B. belcheri* xiamen was only 420,596,363 bp.

As for the completeness, genome assemblies of all four amphioxus were higher than 95.7%. Based on the BUSCO completeness results of genome assemblies, we assessed the similarity among four amphioxus. The *de novo* assembled genome of *B. belcheri* beihai shared 97.4% identity and 97.86% similarity with that of *B. belcheri* xiamen respectively, which supported that *B. belcheri* beihai and *B. belcheri* xiamen are two *B. belcheri* subspecies.

##### *1.3 Genome annotation*

The genomes of the Xiamen amphioxus and the Florida amphioxus were directly downloaded from NCBI GenBank accessions GCA_001625305.1 and GCA_000003815.2 respectively. The genome assemblies of *B. belcheri* beihai and *B. lanceolatum* were annotated in this study. After repeat masking performed by *de novo* prediction with RepeatModeler (Flynn et al., 2020) and then masking with RepeatMasker, genome annotation was performed with both evidence-based annotation and gene prediction. The annotated protein sequences of the Xiamen amphioxus and the Florida amphioxus (NCBI GenBank accessions GCA_001625305.1 and GCA_000003815.2 respectively) were downloaded as protein homologs. The transcriptome data deposited under the NCBI BioProject accession: PRJNA310680 and PRJNA416866 were used for the annotations of *B. belcheri* beihai and *B. lanceolatum* respectively (Yang et al., 2016).

The two published genome annotations of *B. belcheri* xiamen and *B. floridae* contained 23,855 and 26,676 protein-coding genes respectively, while the two newly annotations of *B. belcheri* beihai and *B. lanceolatum* had 44,745 and 38,190 protein-coding genes respectively. The huge gene number difference was possibly caused by the different annotation pipelines among the four amphioxus genomes. The completeness of genome annotations ranged from 91.9% of *B. lanceolatum* to 98.7% of *B. belcheri* xiamen. The repeat contents of four amphioxus genomes ranged from 33.69% of *B. lanceolatum* to 37.21% of *B. belcheri* beihai, of which most repeats are interspersed repeats.

**Supplementary Note 2. Deuterostome phylogeny and comparative genomics**

*2.1 Collection of genome assemblies*

Besides the four amphioxus genomes above, genome assemblies of six vertebrates, seven urochordates, five echinoderms, two hemichordates, three protostomes, a cnidarian and a poriferan were downloaded from NCBI GenBank database (Table S2). The two non-bilaterian species including the cnidarian *Dendronephthya gigantea* and the poriferan *Amphimedon queenslandica* were selected as outgroups in phylogenetic analysis.

*2.2 Phylogenetic analysis*

To perform phylogenetic analysis of deuterostomes, the genome assemblies were assessed by BUSCO v3.1.0 (Simão et al., 2015) with database metazoa_odb9 and then the overlapped single-copy BUSCOs were extracted in protein sequences and generated sequence alignments.

The phylogenetic tree was constructed based on 29 genome assemblies. Among these genome assemblies, 112 single-copy BUSCOs were overlapped, then aligned by MAFFT (Katoh et al., 2002) and edited in Gblocks (Castresana, 2000) to generate sequence alignment of 34,094 conserved amino-acid residues. Based on the sequence alignment, the phylogenetic tree was constructed by RAxML v8.2.12 (Stamatakis, 2014) with the options '-m PROTCATWAG -f a -# 100' in maximum likelihood algorithm and 100 bootstrap replicates, and finally edited by the online tool Interactive Tree of Life (iTOL) (Letunic and Bork, 2021).

The ultrametric time tree was constructed by MCMCTree program in PAML v4.9j (Yang, 2007), with the alignment of codon sequences of conserved amino-acid residues. In the ultrametric time tree, amphioxus diverged from other chordates around 574 million years ago (Ma); then only around 29 million years later, tunicates branched out from vertebrates (Fig. S2A).

*2.3 Comparative genomics*

Phylogenomic orthology comparative analysis was performed for annotated genomes of 21 species.

In the orthogroup assignment, the genomes of four amphioxus, *B. belcheri* beihai, *B. belcheri* xiamen, *B. floridae* and *B. lanceolatum*, had the highest orthogroup numbers, 15,840, 13,924, 13,857 and 14,208, respectively (Fig. S2A). To avoid the bias caused by dense sampling of four amphioxus genomes, we repeated the orthogroup assignment of only *B. belcheri* beihai and observed the same result that amphioxus still had the highest orthogroup number. 1,438 and 615 species-specific orthogroups were identified in *B. belcheri* beihai and *B. lanceolatum* respectively, but none were in *B. belcheri* xiamen and *B. floridae*, which could be caused by the new annotation pipeline of *B. belcheri* beihai and *B. lanceolatum* that identified many new genes. In addition, high numbers of deuterostome specific, chordate specific and amphioxus specific orthogroups were observed in four amphioxus genomes. In the gene family history, over 10,000 orthogroups contracted at stem jawed vertebrates (*Mus musculus*, *Gallus gallus*, *Xenopus tropicalis* and *Danio rerio*), stem tunicates (*Ciona intestinalis* and *Styela clava*), and stem Ambulacraria (Hemichordata and Echinodermata), while only 3,750 orthogroups contracted at the root of four amphioxus (Fig. S2A).

There were 2,241 amphioxus-specific orthogroups that existed in all the four amphioxus genomes (Fig. S2B). Within these amphioxus-specific orthogroups, 3,711, 2,828, 2,916 and 3,436 genes were identified in the four amphioxus genomes of *B. belcheri* beihai, *B. belcheri* xiamen, *B. floridae* and *B. lanceolatum*, respectively (Fig. S2B). To further understand these amphioxus-specific orthogroups, the 3,711 *B. belcheri* beihai genes were assigned into 2,794 functional COGs by eggnog-mapper v2.1.5 (Fig. S2C). The top three COGs were "Signal transduction mechanisms" (22.76%), "Posttranslational modification, protein turnover, chaperones" (10.45%) and "Transcription" (5.55%).

#### Supplementary Note 3. Gene novelties in amphioxus

##### *3.1 Brief summary of gene novelties*

In this study, we discarded all low-confidence candidate gene novelties and mainly accepted those candidates that shared high similarity with genes from unusual taxonomic groups (Table 1, Fig. S3). Notably, those candidate genes conserved among all deuterostome groups, especially jawed vertebrates, were filtered out in this study, such as the sialidase gene (Fig. S4), to avoid repeated the gene novelties of chordates (Holland et al., 2008) and deuterostomes (Simakov et al., 2015). The single copy sialidases of four amphioxus were unusually clustered with bacterial genes (bootstrap value 72) and shared higher similarities with those from hemichordates and echinoderms than with those from vertebrates (Fig. S4).

A total of 28 gene novelties were identified and summarized in Table 1. With gene duplications, one ancient event could finally generate a group of genes, such as the tandem arrayed EF-CaBPs. Among all 28 gene novelties, 17 ones underwent gene duplications in *B. belcheri* beihai and only 11 ones were still single-copy genes (Table 1). According to the annotated gene functions, we assigned these 28 gene novelties into 6 categories, namely innate immunity, glycolysis, calcium balance, amino acid metabolism, other metabolic genes, and others without explicit functional domains (Table 1). For better reproducibility, we have attached the protein sequences in Table S4, and gene locus tags could also be found in the genome annotation deposited in the NCBI database (NCBI GenBank accession GCA_025402995.1).

Except those fully discussed in the main text, other gene novelties were further discussed in these supplementary notes, including other three related to innate immunity, seven related to amino acid metabolism, three with metabolic functions and seven without explicit functional domains (Table 1).

##### *3.2 Gene expression in amphioxus tissues*

Of the four amphioxus genomes, the European amphioxus, *B. lanceolatum*, had a set of high-quality transcriptome data from different tissues (Fig. S5). Therefore, to explore expression levels of these novel genes, we collected the transcriptome data of *B. lanceolatum* from the NCBI SRA database as follows. The SRA accessions of the transcriptome data in the NCBI database are epidermis: SRR6246024; cirri: SRR6246021; gill 1: SRR6246026; gill 2: SRR6246027; gut 1: SRR6246028; gut 2: SRR6246029; hepatic diverticulum 1: SRR6246030; hepatic diverticulum 2: SRR6246031; male gonad 1: SRR6246032; male gonad 2: SRR6246033; female gonad: SRR6246025; muscle 1: SRR6246034; muscle 2: SRR6246035; neural tube 1: SRR6246036; neural tube 2: SRR6246037.

The relative gene expression levels normalized by two references genes *GAPDH* and *EF1A* were presented in Fig. S5. Notably, all those novel genes had expression in at least one tissue of *B. lanceolatum*, although some gene expression levels are quite low (Fig. S5). The most obvious gene expression features in both two normalization results showed the calcium-binding proteins, including a group of EF-CaBPs and the CML26 genes, were highly expressed in guts and hepatic diverticula. Besides, two muscle tissues presented different gene expressions, especially one muscle tissue highly expressed the calcium-binding proteins like the guts and hepatic diverticula, which suggested possible muscle tissue differentiation. Although *EF1A* was suggested a stably expression gene among tissues *(Zhang et al., 2016)*, the two male gonad tissues showed totally different expression levels between two normalization. Because we identified many duplicated beta actin genes in amphioxus, we did not consider using beta actin as a reference gene.

*3.3 Other gene novelties related to innate immunity*

Ricin B-like lectins (RBLs) are a family of carbohydrate-binding proteins that were reported to function in antifungal immunity of the Colorado potato beetle (Rotskaya et al., 2021). An RBL gene was identified to be similar to bacterial genes (Table 1). Three proximally arrayed RBL genes were identified in *B. floridae* (Fig. S9A). Except for *RBL_Bl_1* (Fig. S9A) of *B. lanceolatum*, other RBL genes encode two tandem lectin domains with low similarity. Their N- and C- terminal domains are two distinct ricin B-like lectins (Fig. S9B) that share only 42% identity (in *RBL_Bbb_1*), although the conserved splice sites supporting two domains were generated via duplication of one ancestral gene (Fig. S9C).

Toll/interleukin-1 receptor (TIR) domains are protein interaction domains responsible for interactions among TIR-containing proteins in innate immunity (Pawson and Nash, 2003). Two groups of TIR domains (TIR-a and TIR-b) were bacterial-TIR-like and assembled with various immune-related functional domains. TIR-a is a smaller group than TIR-b (Table 1), and all TIR-a-containing genes are proximally arrayed (distance <20 Kb) in all four amphioxus (Fig. S10A). Four TIR-a-containing genes of *B. belcheri* beihai, five genes of *B. belcheri* xiamen, and two genes of *B. floridae* and *B. lanceolatum* were identified, and some of these genes were combined with two DEATH domains or a DED domain (Fig. S10A). Compared with TIR-a-containing genes, TIR-b-containing genes are much more complicated. A number of TIR-b-containing genes were analyzed to determine the domain structure (Fig. S10B) and a group of Myd88-like genes that contain a CARD domain, a group of RIG-I/MDA5-like genes containing two helicase domains and a group of LRR-containing genes were identified in the domain structure (Fig. S10B). None of the Myd88-like and RIG-I/MDA5-like genes containing the TIR-b domain were identified in *B. lanceolatum* (Fig. S10B). Compared with the other three amphioxus, *B. lanceolatum* has fewer and less diverse gene models containing these two TIR domains. Toll-like receptor (TLR) models in the amphioxus genome were reported to be much more complex than those in the human genome (Dishaw et al., 2012). These TIR domain-containing genes are widely expressed and relatively highly expressed in gills, which are important structures in the amphioxus immune system (Fig. S5) (Han et al., 2010).

*3.4 Gene novelties related to amino acid metabolism*

Seven gene novelties related to amino acid metabolism were identified in amphioxus (Table 1), including two related to DOPA metabolism and two involved in folate metabolism.

Serine carboxypeptidase (SCP)-like proteins have roles in stress response, growth, development and pathogen defense in plants (Xu et al., 2021). The SCP24 gene of amphioxus was identified as a gene novelty (Table 1). In the phylogenetic tree (Fig. S15A), the SCP24 genes of amphioxus are clustered with those of the tunicate *Ciona intestinalis*, plants (including *Klebsormidium nitens*) and other eukaryotes, but not other metazoa (excluding Amphioxiformes). The SCP24 gene of amphioxus shared one conserved splice site with those of the tunicate (GenBank accession: XP_002131445.1) and the charophyte alga *Klebsormidium nitens* (GenBank accession: GAQ88391.1) (Fig. S15B).

Glutamate-1-semialdehyde aminotransferase belongs to pyridoxal-5'-phosphate-dependent enzyme and participates in porphyrin synthesis (Grimm et al., 1991). A glutamate-1-semialdehyde aminotransferase (GSA) gene highly conserved among four amphioxus was identified as a gene novelty possibly originated from Deltaproteobacteria (Table 1) and only duplicated in *B. belcheri* beihai. In phylogenetic tree, the GSA genes of amphioxus clustered with that of two starfishes (*Acanthaster planci* and *Patiria miniate*) and deltaproteobacteria (Fig. S16A). In gene sequence alignment, this GSA gene was highly conserved among four amphioxus and two starfishes, in which two conserved splice sites were highlighted (Fig. S16B).

In addition, a group of GNAT family N-acetyltransferase (GNAT) genes (Table 1) that catalyze acetylation of various substrates were identified to resemble those from Alphaproteobacteria or cyanobacteria. Horizontal gene transfer (HGT) of GNATs was reported in a soybean cyst nematode (Noon and Baum, 2016). An aralkylamine N-acetyltransferase (AANAT in the GNAT superfamily) of *B. floridae* was reported as a possible HGT gene (Coon and Klein, 2006), but it is not one of the GNAT genes identified in this study.

DOPA decarboxylase (DDC) also belongs to pyridoxal-5'-phosphate-dependent enzyme and is responsible for the synthesis of dopamine. A group of DDC genes were identified as novel genes of amphioxus and the hemichordate *Saccoglossus kowalevskii* (Fig. S17A) with conserved splice sites (Fig. S17B). These DDC genes duplicated in amphioxus and were classified into 5 clusters (Fig. S17A). In Cluster 1 (Fig. S17A), *B. floridae* and *B. lanceolatum*, but not two *B. belcheri*, had a unique subcluster. In addition, DOPA 4,5-dioxygenase (DOD), a key enzyme of betalain biosynthesis was also identified as a novel gene of four amphioxus. The HGT of DOD gene in the domestic silkworm, *Bombyx mori* was reported as contributing to its dopa metabolism and antimicrobial activity (Wang et al., 2019).

Folate receptors (FR) could not only bind folates and folate conjugates with high affinity, but also mediates intracellular delivery of tetrahydrofolate (Wibowo et al., 2013). Dihydrofolate reductase (DHFR) plays a central role in folate metabolism and could enzymatically reduce dihydrofolic acid to tetrahydrofolic acid (Schnell et al., 2004). Two novel genes were identified as folate metabolism related (Table 1), including a folate receptor and a dihydrofolate reductase, both of which are important in folate-mediated one carbon metabolism (Fox and Stover, 2008). The FR genes were identified from chlorarachniophytes, a group of unicellular green protists, while the single-copy DHFR gene were identified from bacteria (Table 1). Four copies of the FR gene were identified in *B. lanceolatum*, in which *FR_Bl_1* was relatively highly expressed in epidermis, cirri and gills, while *FR_Bl_4* was highly expressed in guts and hepatic diverticula (Fig. S5).

*3.5 Gene novelties related to other metabolic functions*

Besides the four functional categories above, other gene novelties involved apparent metabolic functional genes were listed in Table 1.

Glycoside hydrolase family 27 (GH27) is a widespread family of glycoside hydrolases (Fernández-Leiro et al., 2010). A single-copy GH27 gene was identified as a novel gene in amphioxus. The best hit of these GH27 genes of amphioxus in the NCBI NR database was from Terrabacteria group, while the best hit in the UniRef50 database was from plant (Table 1). In the phylogenetic tree, the GH27 genes of amphioxus were clustered with those of other eukaryotes (mainly algae), plants and bacteria, but far away from those of other metazoa (excluding Amphioxiformes) (Fig. S18). Because we did not identify any conserved splice sites of the GH27 genes with other eukaryotic genes, we cannot conclude the origin of this GH27 gene.

Multicopper oxidases (MCOs) is a group of enzymes oxidizing diverse substrates using $Cu^{2+}$ as cofactors (Komori and Higuchi, 2015). A group of MCO genes of amphioxus that had the best hit from plants, could be classified into four clusters and have homologs from two tunicates, but no conserved splice sites were identified (Fig. S19). Anhydro-N-acetylmuramic acid kinase (anmK) is a key gene involved in recycle of bacterial cell wall (Uehara et al., 2005). It is interesting that a single copy anmK gene was identified to share high similarity with bacterial genes (Table 1). We cannot predict its function in amphioxus adaptation.

##### *3.6 Gene novelties without apparent functional domains*

Many other novel genes without apparent function were identified, such as a single copy tetratricopeptide repeat protein (TPR) gene of amphioxus was identified as a novel gene that shares high similarity to that of amoeba (Table 1).

Based on the Pfam search results (Fig. S20A), two large novel genes of amphioxus were tentatively named as apolipophorin III-like protein (ApoLp) and Hedgehog/Intein domain-containing protein (HintN) (1205 and 789 aa respectively) (Table 1). These two genes, ApoLps and HintNs were both highly similar to those of gammaproteobacteria and deltaproteobacteria respectively. Since the ApoLp and HintN genes, *Bbb_173200.1* and *Bbb_173190.1* adjacently located on the genome of *B. belcheri* beihai (Fig. S20A), we considered that they were possibly horizontally transferred in a single event and then diverged in four amphioxus genomes. These two genes are tandemly arrayed in the Gammaproteobacterium *Microbulbifer taiwanensis* (Fig. S20B). Three pairs of opposing ApoLp and HintN genes were identified in both *B. belcheri* beihai and *B. lanceolatum*, but only one pair in *B. beicheri* xiamen and two pairs in *B. floridae* were identified (Fig. S20B).

A group of atypical short-chain dehydrogenase/reductase subgroup 4 (SDR_a4) genes were identified mostly proximally arrayed (distance <20 Kb) and divided into two clusters in four amphioxus (Fig. S21). The SDR_a4 gene was tandemly duplicated in all four amphioxus (Fig. S21A) and could be divided into two clusters in the phylogenetic tree (Fig. S21B).

A group of novel genes were characterized as cytochrome P460-like genes (CytP460s, Fig. S22A). Tandem duplications were observed among these CytP460 genes (Fig. S22B). One of the best matches in NCBI GenBank database was from *Cadophora malorum* (GenBank accession: KAG4419885.1) and the hit E-value of the CytP460 genes from amphioxus was as low as 2.73E-107. The phylogenetic analysis divided those CytP460 genes into two large clusters, in which the cluster could be further grouped into two clusters and none of *B. lanceolatum* was identified in Cluster 2 (Fig. S22C).

In addition, three transmembrane proteins of unknown function (UF-TM1, 2 and 3) were identified as novel genes (Table 1), in which UF-TM1 has the best match from the charophyte alga *Klebsormidium nitens*, UF-TM2 has homolog from the hemichordate *Saccoglossus kowalevskii* (Fig. S23), UF-TM3 has homolog from tunicates (Fig. S24). The UF-TM1 had 12 copies in *B. lanceolatum*. In NCBI NR database, the best hit of UF-TM1 (using the representative gene Bbb_158830.1, Table 1) was from the charophyte alga *Klebsormidium nitens* (GenBank accession: GAQ78278.1) with a E-value of 1E-33 and 33.62% identity. Three highly conserved splice sites were identified among the UF-TM2 genes of amphioxus and that of the hemichordate *Saccoglossus kowalevskii* (Fig. S23). As for UF-TM3, homologs were identified in tunicates and tandem duplications were observed among four amphioxus (Fig. S24).

**Table S1. Overview statistics of the genome assemblies and annotations**

For better distinction between two *B. belcheri* subspecies in this study, the Beihai amphioxus and the Xiamen amphioxus are tentatively named as *B. belcheri* beihai and *B. belcheri* xiamen, respectively.

| | ***B. belcheri*** **beihai** | ***B. belcheri*** **xiamen** | ***B. floridae*** | ***B. lanceolatum*** |
|---|---|---|---|---|
| Assembly statistics | | | | |
| **Assembly size (bp)** | 478,319,013 | 426,123,519 | 513,460,931 | 495,286,038 |
| **Ungapped length** | 477,574,676 | 420,596,363 | 487,128,935 | 474,945,525 |
| **Scaffold number** | 583 | 2,308 | 432 | 10,247 |
| **Scaffold N50 (bp)** | 4,185,906 | 2,325,619 | 25,441,410 | 1,295,816 |
| **Gap content** | 0.156% | 1.297% | 5.128% | 4.120% |
| **GC content** | 41.46% | 41.32% | 41.25% | 41.49% |
| **BUSCO completeness** | 97.2% | 97.2% | 95.7% | 95.8% |
| Complete and single-copy | 94.4% | 95.8% | 94.7% | 93.1% |
| Complete and duplicated | 2.8% | 1.4% | 1.0% | 2.7% |
| Fragmented | 0.2% | 0.7% | 1.2% | 2.5% |
| Missing | 2.6% | 2.1% | 3.1% | 1.7% |
| Annotation statistics | | | | |
| **Protein-coding genes** | 44,745 | 23,855 | 26,676 | 38,190 |
| **BUSCO completeness** | 95.2% | 98.7% | 96.6% | 91.9% |
| Complete and single-copy | 91.3% | 97.5% | 93.8% | 88.0% |
| Complete and duplicated | 3.9% | 1.2% | 2.8% | 3.9% |
| Fragmented | 2.5% | 0.6% | 1.7% | 4.3% |
| Missing | 2.3% | 0.7% | 1.7% | 3.8% |
| **Repeat content** | 37.21% | 35.97% | 36.12% | 33.69% |
| Interspersed repeats | 35.47% | 34.47% | 34.93% | 32.55% |
| Retrotransposons | 9.47% | 9.31% | 8.82% | 8.50% |
| DNA transposons | 13.26% | 13.09% | 15.00% | 11.97% |
| Simple repeats | 1.24% | 1.05% | 0.86% | 0.94% |

#### Table S2. NCBI GenBank accessions of genome assemblies

| Taxonomical groups | | Species name | Description | NCBI GenBank accession |
|---|---|---|---|---|
| Deuterostomia | Vertebrata | *Mus musculus* | House mouse | GCA_000001635.9 |
| | | *Gallus gallus* | Chicken | GCA_016699485.1 |
| | | *Xenopus tropicalis* | Tropical clawed frog | GCA_000004195.4 |
| | | *Danio rerio* | Zebrafish | GCA_000002035.4 |
| | | *Petromyzon marinus* | Sea lamprey | GCA_010993605.1 |
| | | *Eptatretus burgeri* | Inshore hagfish | GCA_900186335.2 |
| | Urochordata | *Ciona intestinalis* | Vase tunicate | GCA_000224145.2 |
| | | *Styela clava* | Asian tunicate | GCA_013122585.2 |
| | | *Phallusia fumigata* | Black sea squirt | GCA_008931825.1 |
| | | *Phallusia mammillata* | Warty sea squirt | GCA_003260075.1 |
| | | *Halocynthia aurantium* | Pacific sea peach | GCA_013436065.1 |
| | | *Halocynthia roretzi* | Sea pineapple | GCA_013436055.1 |
| | | *Aplidium turbinatum* | Colonial sea squirt | GCA_918807975.1 |
| | Cephalochordata | *Branchiostoma belcheri* beihai | Beihai amphioxus | GCA_025402995.1 |
| | | *Branchiostoma belcheri* xiamen | Xiamen amphioxus | GCA_001625305.1 |
| | | *Branchiostoma floridae* | Florida amphioxus | GCA_000003815.2 |
| | | *Branchiostoma lanceolatum* | European amphioxus | GCA_900088365.1 |
| | Echinodermata | *Acanthaster planci* | Crown-of-thorns starfish | GCA_001949145.1 |
| | | *Asterias rubens* | European starfish | GCA_902459465.3 |
| | | *Patiria miniata* | Bat starfish | GCA_015706575.1 |
| | | *Lytechinus variegatus* | Green sea urchin | GCA_018143015.1 |
| | | *Chiridota heheva* | Sea cucumber | GCA_020152595.1 |
| | Hemichordata | *Ptychodera flava* | Acorn worm | GCA_001465055.1 |
| | | *Saccoglossus kowalevskii* | Acorn worm | GCA_000003605.1 |
| **Protostomia** | | *Helobdella robusta* | Segmented worm | GCA_000326865.1 |
| | | *Drosophila melanogaster* | Fruit fly | GCA_000001215.4 |
| | | *Caenorhabditis elegans* | Roundworm | GCA_000002985.3 |
| **Cnidaria** | | *Dendronephthya gigantea* | Soft coral | GCA_004324835.1 |
| **Porifera** | | *Amphimedon queenslandica* | Sponge | GCA_016292275.1 |

**Table S3. Conservation analysis of the 28 genes of amphioxus with other Deuterostome species**

Conservation analysis of the 28 genes in Table 1 were searched in five major groups of Deuterostomes (except jawed vertebrates). Grey background means this gene novelty is conserved in the species and the star (*) indicates sharing conserved splice site with the genes of amphioxus.

| **Functional category** | **Abbrev.** | **Deuterostomia groups** | | | | |
|---|---|---|---|---|---|---|
| | | Jawless vertebrates | Urochordata | Cephalochordata | Echinodermata | Hemichordata |
| **Innate immunity** | LOX | * | | | | |
| | DBH | | | | | * |
| | RBL | | | | | |
| | TIR-a | | | | | |
| | TIR-b | | | | | |
| **Glycolysis** | ADH | | | | | |
| | FDX | | | | | |
| **Calcium balance** | VDCC | | | | * | * |
| | CML26 | | | | | |
| | EF-CaBP | | | | | |
| **Amino acid metabolism** | SCP24 | | * | | | |
| | GSA | | | | * | |
| | GNAT | | | | | |
| | DDC | | | | | * |
| | DOD | | | | | |
| | FR | | | | | |
| | DHFR | | | | | |
| **Other metabolic genes** | GH27 | | | | | |
| | MCO | | | | | |
| | anmK | | | | | |
| **Others** | TPR | | | | | |
| | ApoLp | | | | | |
| | HintN | | | | | |
| | SDR_a4 | | | | | |
| | CytP460 | | | | | |
| | UF-TM1 | | | | | |
| | UF-TM2 | | | | | * |
| | UF-TM3 | | | | | |

**Table S4. Protein sequences of 28 representative genes**

For the 28 genes (Table 1), the protein sequences of representative genes are listed as follows.

| | |
|---|---|
| LOX | MKAQLRDLISIARGVPARDPPGSPFGPDRDPKAARPNCQSRSALCRTLHGSRRETVDRALDGRHTLGFMTPAGLYTENTPSRPRFKCDSSLIKDLAFGKEI<br>TLTHTEVVPVSAEVYWSGFDDFFHLRSFLHANVETIKGDGGVGTVVEFDFPGGRAQEELVEKDEATKTWAVSMLGSNPVFKSYLVTFKVDEDTSETAK<br>VTVTAKGIFALEDYNQRRARIAFSKYMLRSHINKIMAAIADKKGDVLQFDVDVDCSFQKMWDTFRDWGNVSWMMGATGVQVLPDYQNTDLRRKVLF<br>GAGSVEERLVGATEKPASLTFEVFKNDAMRVLYFRKKLELVKISERKTKIKSTSYFVPQPGPSPKATLDALVVPSLKWIGNAFAEKPKSRPGPAEKRSIKT<br>SAIVTKLVDDCILPEGFLAEGAEVFLEHTEEMPISAEDYWSGFQDFVQLQLYMEGHEGVKVALGHVVFGQSNGEVGTVVSFYYDPTDLAEGQREMKLV<br>EKDDSTRTWRVQEAEPNTLYKSYEMTIKVTGTTNASVTVSARFVSAIKNDEERRDDIAYHKVHLLGATLRDVLNFVVAEKLGRKYQVDQEVDISREKL<br>WCIMGNFNDVTWIPNAKYAQVNGVSRQIMFSRDPTCHEGERTLDVRLVWYDSSDYLLVLEYTGAMLVTKFYREKIQLFPVGANRTRVAYECIVLPAKD<br>EPAEGFDKWLIPAISNRVTNLKRMFEDEEDEEDHGDQCTPPSGPPVFPRGAYSTVARGTVHAPVKEVWSAFRPFGKACLEYWKIYESMQIEGPGKDCVG<br>CVRSFVTSKTGSHIKERLEWRDDVNHVEIYSLLSMTPPPPVAMSNVYTTITMTAVGEKETEVKFECTFDVSVSFAVSRIQDNQKAAYMACITGLQQLFHS<br>EVGTLEVVVGSAADLARTDGWFDADPYVVLALNGGKPFTTRVCNGTQTPVWDERFAMSVTPRSRSVVFTVMDRDTVGQDDIMGTANVNLDELTSGQE<br>KRMTLDVQGGGTLNVRLLLKMHDKPKDQEDEGDKLIRSIALPFLAPVAQSELDAIKKEFTNLIMSFVGPGQKYELSYMTRLRTHPDVPLEEYPAACVPM<br>PTGEMFTPLKAGRLMQRLMEFISSQASAHYFAGNSPAVPGKLLGNHTRTTFVNGQIMVRLAQAKGIWDPWKANFSGYLPANERLIEEWQKDEEFCRQY<br>LQGINPMLVTVCKDQSQIPAEMLGLKGQGKTTLELMAENRLFIVDYAPMLGVPAVPGKFIYAPIVLMYKEELDGGKSRLNMLGIQLTRDKGNNEVYSPE<br>SAKTHPNKYMFAKMHVQSADNNVHQFLYHLGYTHLGMEPLVVAMYTHLPPDHPVHRLLLPHFKDTIGINYLARHSLVSRIFPITDPMFATSTVGGLVMF<br>LKEWRKYNFMDMAFPEELKRRGFDEAGTDGLKDYFYRDDGFKLWNIYKSYVTGMVNHAYADDQAVQADKALQEFCKMTAGPGQLRGFPREISDKKL<br>LIDCLTNIIFNVSAQHSAINFPQYDYYSFVPNRPAQLSMPMPDGPNDMLQSTVLEALPPPHFTALQVLLSYMLSMPSQTAITGVEAMKEVYPEVHETFNTQ<br>LKELSKEIQTRNEGLKTEGKAAYTYLDPENVAMSIDI |
| DBH | MVRNTYGKTAWGGAFLDALGRLERGKSYANTGKVLNVNINGPKVSAHVQGTAPRPYEVKIQFQLLDHAEVQRVYDVINENPLLYGQIINGDLPDELVD<br>TLEEADIDLLPASWYDMKARCKCPDEANPCKHMAAVYYLITSEIDKNPFTLFNLRGVDLMGYFNISPSDTDPGYPLPLNRPSSMQSTETEVNENHILPCPL<br>LKIPPMSDFILSCLKSNPPFSLELDYKCVMEEFYSYAAKQSTKVLDDLMNPGKAGSAHWLSELDYDEVQNIMFDYTGAEYAPQLQIHNPVWDNLSEHES<br>ENKEWEFIKRVFKDEMPSYDAAFIPDVLPLSRQQQDKIKDFKIIWRPAHTAELVAEQLDNLTKIYPEDNPCVEVDNMLLGGHSAALHVLSTLLTRFVTDL<br>HFMHKKSKNNPPPESLAFFRGSVYNIRSIAQQSVPRAIAKYFGDFDLIKTDLEVAVHISPLKTKGFKIKNEELPPPDHYAMELWVKMIDNDIARHKPIRRFL<br>MSNSWAKAQAMKLLATLHSYLPHVGRLLKEDSIALTAAELEDFILNTVDVFNNLGVKVILPKELRKILKPKAVVYAELEGSKNQQSFFSIGDLLTYNWKI<br>AIGDEYIAVHEFERLVHSGEGLVRFRDKYISVKPEEMTTILNNVKQEPPQPSPLDLLKESLLEDSKFHLSASLSSMVESVKNVEEHPIPHNLQATLRPYQVS<br>GYRWLVSNMQHGFGVILADDMGLGKTIQSIAAMLHLKNRGDLKDPVLLVVPTSVMSNWEREIQRFAPSLTVSRYYGAGRMLPSSQCHGNSDDLALQEY<br>ENQPDTPNGAGKRPSPGDGPKKKRARLSWDIPQTDIIITTYHIVRIDVDALAKIQYSMVLIDEAQYIKNCKSQTAKAVKRMKAKMHVALSGTPVENNLTE<br>LWSVFDFTFPKYLGTNKDFTQTYAKPIEVHRDPDRVEALRAVTRPFLLRRLKTDRSIIKDLPDKITTPKYNSLTAEQAALYESVVQNMVKKLAEAKEKKE<br>RTGVIFQTMTFLKQICNHPANFTKNSNRDIHASGKMKVLIDLLQPILQQGEKVLIFSQYVQMIKLMAHMVEGDLKVKPLIFEGCMTQPQRDEAITAFQTQP<br>HRQIMIVSLQAGGVGLNLTAANHVIHYDLWFNPAKENQATDRAFRIGQTKTVFVYRFISENTFEEKIDAMLEKKKDLSNLSVQAGETWIGNLNDDEITQL<br>FTRGSPWGNRFAQREARVVECASSYAVDPADAPPDSAVNAHIDPAHSSRISG |
| RBL | MPGATRFWHPLKTPRQNKNPCLTTSDAQSSSSVVKSVTVSRSAYINSIQTIAKYLVERSKESNTSMLAKRGRGAIIVVVDLIGQYRRCGTTGSTTVYAFTE<br>HSVARIRKLDYPVVKHTASAFSSSDVSSLQMPRFEYIRRSVLESRERFARFNRERGPREQQQEQAARPEVLAIARRPGFTGASAGLRLAVAIHLSWAEALQ<br>TALGSTVRSSTPPNRHGGYFYIMSRQTGMVLDVFEGNTAASTRLIVYTMKTEGPENQLWKYDSESGELVSRLNGYRVDVSGGYAFDGAKIINYPTNNAP<br>NQNWVFKPDGSIQSGLGDQWVLDVEGANTNAGTHVILYQRKDGGPSINQQWDIVPYVPTTGFYIRSALNDCYLQIADGSDKTSAPLCIGNKPGVPRADH<br>RYLLWKYNKEDRTICSELNDYAVDVQGGSAENSVKIIAYPCHEGANQQWTFEKSGTIVSGLGENWVLDDQASGGAGTPVILYTRHGGKNQQWYISED |
| TIR-a | MAHATPTASTGKGTKRKRDNGEGRASSSSETPKRVFISYSMDPYIDQSRTPLQRSIDVKEQRNKVKALADRLRNDGVDGWIDQYDEYNPPELWTRWME<br>EEIVKADYVLMICSPNYKECVMGERNEEAVAGFGVRFEGKVIYSLLSNPQNHGKFLPVFFGPIERDHVPDVFGGANFFGPIVTPPTLEGEYLKLLSKILGRN<br>PPGQGPPPVRQPAF |
| TIR-b | MASGDNKETIEDFVTDNYDHLSKRLQVEKLIPHFIQSRKLDLPDKQVIMSKVTTRGKAEALLDILIENGKCSPDEFVEILRNGDHEHVADQLRRTSTQNES<br>KEGPQVFICHAGPDKGRFVRPLVERLQDEGLPAEKLFYDEISLQPGDVIDDKIRATLSSPSLKLVVIVISRHVLNDRYWPKLEVELTLKANKKVLPIWLDQ<br>NEDGFTAFGKRLREYSPTLKGIVGRKVSPQRAGREIPGIAEEIVRKLEIA |
| ADH | MSSVPGIPAKMKGVVCYAPGDYRMEALDVPTVGPEEVLVKVTAVGICAGDAKCFAGAPLFWGDKDREPYCQPPIVPGHEFIGEVVALGPGAAEKFSLEL<br>GDQAIAESIVPCWKCRYCLRGQYHMCMPHDIFGFHQRTPGAMAGYMKYISDSIIHKVPKSVPPHHAAFIEPLACSIHAVERGEIQFRDVVVVSGCGPLGL<br>GMVAAAKQKNPAKLIALDLYDWKLEVAKKCGADVVLNPGNCDVIAEVKKLTDGYGCDVYIEATGSPVSVKQGLHMIAKLGTFVEFSVFKNETSVDWT<br>IIGDTKRYNTYPKAISMIAKKELPIEDIITQKLPLSDVVKGIDLVNSSSKSIKVVLIPE |
| FDX | MHLFVQSKRNYGLLGITGEETIRTIKERVFKVEGLPVDQQILYYSGKPLEDDATLIGSGIPDMGTLDVNVRTRGGATYKVTLKTPDGERVFDCPDDSYILD<br>AAEELGLDIPYSCRAGACSTCAGKIESGTVDQSDQSFLDDDQMANGYVLTCIACPTSDVVITTHQEEALY |
| VDCC | RNIYACSSPELEEESVCSQEYGASIFTGTGCATCTNRKLSVSGSYIRTPSREATCTDALNRDICLSQGLDTSFLDNWFKDEREPGVKWQFFASQTGLYRMFP<br>GVAQERCHDYDPRLRAWYVSATTGPKDVVIVIDSSDSMALHPWSSPLTLMELTKEAVNSVLWSLTHLDYVAVVSFATLAHQLTIQNKNTLVQATRDNI<br>QALTQEVKKVEPNGHTNFEAAFRTAFDILQRSEVVSRTAGCNKMVLFMTDGEPTRGEREPEQLTQIIRSLNQREDGSKVASIFTYSVGGLADTTITKIIACN<br>EGGVWSEVETAGKASLAEQLSQYYDYYSTLREGQGHVIWTEPYMDAFGAGEVVTAAKAVHVYNTSTVPQTFLGVVGIDVATSATERYTTSKQDLLDTL<br>RSRSSSCADLRESSLCELEQLRQKVYQQGNVRYENNSAKWCHLSSAQECTFPVTEPCENQPQLYRTCMNHSAVKVDYETTSCCGRYDQPCPSTTTRISSS<br>LHLLILPVTFVRSFLYSNILQPLL |
| CML26 | MMLFLALALTMATALPVPRKGEQYFDNLDINGDGIITLHEADHGLSVASIFSAMDADCDGFITRGQILFYMREFVEAFDTLDLNGDGFLTVQEAETATTIG<br>RVFAFFDTNADGQLERWEAERLINLIVLLQDAAAAQGTACATTPSPNHGQGH |
| EF-CaBP | MTPFLLLFVALATASPMKRQTTGLDANGDGLVSKDEVLNTMSLHEALVALDRDGDGFIYLPQIIELWGTGDKFYELNTDGDDHLTFREIENGMTLSDFY<br>DQFDFNGNGILTTTEAYQLHYIYNVIHNNVATDDPLDSNGDGKLSKLEITSHMRYREVLGALDVNGDGELTPLELTILLGGQTGQFVIELDTNNDGTLSIQ<br>EARNTSLRQIFMRLDVDDSGYLENGEGAGFYPVWNTIVVSGISDPITG |
| SCP24 | MVPYVCLLMIFLATCPGQAFSRPETQDSDRIESLPGLNATLPFSQYAGYITVNQSHGRRLFYWFVESQNDPERDPLVLWLNGGPGCSSFNGLFEENGPFSP<br>NPDGKTLDLNPYSWNRNASVIFLESPSGVGFSYSDTKADYTTGRDSLDFMLKFLEKYPQYQKNKFWITGESYAGHYVPNLASHIVSYNIVKPGSINLEGF<br>MVGNAWTDPDLDNAGATFFWWSHALISDRTYNNINKACNYSAIGPLLANGEQTLLSSSPDRLRDECEMLLAEAHTEMGNINIYNIYVDVCLRRRGGRQL<br>LSQLARSDSVLRKFAQRRLQSEDVGRKMYPCEDDYMEKYLNRPDVIATIHAATLPYKWTPCSTIVDYSRKDLLTSMLPVYKRLFSAGLRILVYSGDVDAI<br>VPVTGTRAWLKALKLTEVEGWSAWTASDQQVGGYSVVYDKLMFATVRNAGHEVPGYQPLRALDMFNRFLMNRRL |
| GSA | MDSETDGTPLGQAAQRSLAVFPAGSNGEFNLPKELTTVLSRGKDCYVWDVDGKRYLDFSMGWGTNLVGHAREEVVAAVTEQAANGSNFAYVNEKSL<br>QLAEEIKRLSPAVDYLRFCSSGTEATMYCERLARAFTGRSKVLKFEGAYHGANETGVTSLFPKGLLDYPTPDPSSAGIEHSVKDHVLVAPYNDLSMTTEIV<br>EQHKDDLAAVIVEPLHRCTPPMDGFLQGLRDLTKKHGVLLIFDEVVTGFRLAYGGAQEYYGVIPDLVAYGKALGGGYPIGVFGGIVNEDLLGEDRRYV<br>WTASSLGCNTISASAALAALQIYRRTGTYKQLHDIGKYVRSKMKECLEKNDIPGHVLFTEKLVYNYREQKLQENAARRRAMMVGLFKRGVFLNPMGTK<br>LYLSLKHTEDVCDQFVEIFDQTLQEIKRNEKSATTN |
| GNAT | MFEDSRATESETGGSPDTFRVRQVTSLSEVSLMLEWAAAEGWHPGVYDAESYYRQDPTGFYLAEVDGVPVGGVALVKYGENFAFGGNLIMKPAFRGK<br>GYARRMARVVMEASGMSDRNIGLESMVDLQDAYSRVLGVQFAWKNTRFRGVGSNTFTGVKQDDNPAALVSVKEVDFEYLCVFDESVFGTPRPSFLKS<br>WISQPGMVALAAVTTTPAGEDDEHSQETRIAGYGCVRLSVESYKDVNFSRRIGPIFAESAAVGSQLFAALVATIPPGVPYYIDVPETNESAMEVVRESGLE<br>TVSQTARMFTKGDGGVKCSMVYGITTLELG |
| DDC | MYVRISVDSRPHRDELQASYVRAGNELCDLKNRRRLVADVKNTFRVSASFGRLQKIAGDPENSRDARRNIGRTTAFSSEEVKKSTQEVKNLREKLIDLRE<br>KTGRLDLTESRRKELLDGVSNFADKALDKLQGSKTRIYTGAGDANNNNADSKGIRSSPIQDEPLDLDDILDLYGTHVESRGLRAGHGGHMAYVPNNGVF<br>PSALGDFLADTLCTYAGRAYESPGGVEMDNMLLEWLAGVFGYPKGFRGNLTSGGSAATVIAMATARDSRKGLRARDFHRVVVYVSELTHHCMRKALH<br>TIGLGEAIVREMPVDSRWKMDVQTLETTVEKDAQNGLIPFAVVATVGTTDVGSADPVDAIADVAERYGLWLHVDAAYGGFFALCEDTRPLFRGVERSD<br>SIIVDPHKGLNVPYGSGVVLVRNGRKLHLSNTAHKQGGYLYHPGSDDTELSPCDLSFELSTHFRGPRMWFPLKLFGVEAFRSMLTEKLQLATYFYHIIKD<br>VPGFEVLQAPELSVVAFRYTEPPSGMDANSFNKLLLEEMIQEGGVYLSSTTLKGTFYLRICVLGFRTHVEHVEICLEILQKSLRKLKTGMLS |

(to be continued)

| | |
|---|---|
| DOD | MEDVGLRLAVLCVVVVLCSAHPTKYPEVNLDVYRAKDTPPPFLSYHIHCMFVAYNNESVFAALKLFDRFIEHFNLTHTRPCEGLYHQGRLCAFDVDYTNTGPTDPFVSANWAIFVPLEWYAKTVPVVMQNRGDQDILVHPNSGAEVSDHTRWALWMGEKWPLNSASFTNQTVF |
| FR | MAPMMLVLLAAVAGVAVAGDVPCRPMQELYPSGKELCERMWDGAFVYETDLSRAYTMWFFERDNPNDKVSHKLGKTPPNKCYLQYLHKTVPGPEPDNFTECHPWKTRACCRQQTVSSPEKIIQNYGPEWRWDRCGKLSAACERFFVQEACFYECDPNAGLYRMYSDKDFDASNPRHNKWQIEGMPIRADYCDAWWAACRNDLFCALEQGNHYVCAAEYEKADGSQGYFYTLIKSIVSWFTQ |
| DHFR | MTTPSARIRIYSAMSVDGYIATEDGGVAWLEQFNTADTGYDDFFREISTVVMGRTTYEQVCGFDCPWPYAGKKTVLLTTKNLTNLTTEQTTCYSGDVRDLADDLRKATTAGDIWLVGGRMVTQAFLDARLVDELELYVMPVLLNRGIRMFEPSGGTSNDLTSLKLKESKTFPNGVVKLVYNKADVS |
| GH27 | MDGYSLMMNETAVLQIAAAQRSQLLYYGYDHLVLDAGWFWNITADAPILDDFGRPLPDPERFPSSKGTNGLEILARQVQDKGLKFGVWFLRGIPVQAVKDKLRIYGTQYTADQIALPSTQCSWDNMNYGINTSHPAAMYYYRSITDMFSSWGVDFVKADCFYGGEIHKDEVALFSQAISQTTRTMTLSYSPGNEANVTIAQWLVDNGYADMYRITADTWDSWTGADGIAKQFPAAAAHAKLIGVGDTYPDLDLLPLGHLGHLTGAHSPPRSTRLTMVEQKTAMTLWCMARSPLMFGGKLPTDDMTLGLLTHKELLRLHEFSMGNRPVYQSGDTVVWHAVSTKPVTLVEHYVAFFNLAVDHSLTVAVNFTDLSLPSGQLYSVIDTWDSHNLGLYKDSFSATVYARGSGLYRIIKDLTPPS |
| MCO | MNNPCKNFTVGNDLVRPDEHYSGNETRLNVTLTVDIGDVTIDWFTFPRRLYNGKMPGPTLRVRAGDTLNIKLINELGGDQEARPNNVPRHPNNTNLHVHGLHVSPLEPQDSPFTLVAPGESYQYQIEIPQDHASGTFWYHSHHHGATSFHMNGGMAGFLIVEDDPATMSDELDAISCPNNCQNDLPLLFGPWFKYAVPSPFLAPLSFAVLQDIWGDAVRLPVYNVTENTTLADWLMNPANGVDYFLTNGQLQPTITLKRREIKRFRMLTAGMWELLFITIEGSSCELQVLAWDGLYVDRPYTVNKVVLGPGQRLDLAVRCSQEGSFKMKSAPTDDDLLSVGPESVPYTGILATILVNDTINAIMDFPYELPSRPAYNADLRNLTEKQIQGRFVVEFGPGFEFNRQPYVSPTYYRYKAEVDTYQEWHILNTNPLKVHPFHPHVHHFQVMSYNKYEGPYGHLDVFAGLASRAYFDLQGKLCYRQYPGFDASAARPDRLPLVLTHAWNETWTDEKSIYNPVGQWRDTVIIPPLSNVTIRLQLHRYRGPLYVHCHTLFHSDHGMSGVVQVVGPGEPTGGAHVSAGGVWPGTCRRTDVYSPDTYGAGGVQVPSRAGPPQRLAAHRGWHVWAAASVMFLCLQTKLY |
| anmK | MEFTTKTTRALGLMSGTSMDGIDAALVDIIEHDNSRLEVRQVVAETYPYDSRLREDLGAVCSGKQLTVDQLCDLDDRVAEAFAETATTVMDREATKPTVIGSHGQTVFHRPPALTETGGTRLGHSVQLGRGAVIARRTGVVVVSDFRKADIEVGGQGAPLVPMVDWLLLADRRRTRTCQNIGGIGNVTYLPPNGNHDQSIAYSYREFLPKGPDEVLVSGGGSHNSFLMDRLREALHPVKVSTTEAVGINVDYKEAVAFAVLGYMRMREQPGNLPSVTGATRQVLLGEVSHPS |
| TPR | MADNVLVSSQRHYIQGRYNDAIKLLSQALKDKGRSLEPKDTAAIHCNLASCYLKLGLLRKVVSSSDAAIKVDASSLQAYLYKGKALHRLKKPDQAREVWESALSRYGDFQVLTEILDCIQCLEDEKSGNTQAQQSTNIKVNGDADSERSTTPQPMQEGTAESQQDNKKTTGGFLDPKSTEFQVARGFLLVNESNYTEAIKLFNTILQLHPMTYKAYVGRGTAYAMTRQLLKAEADFTQALTLEPKEAEIWKRRGQVRAALGKESQAIEDLNQAQSIAKDVDVDIQKEVGLIYQAKGDNVSALMHFKRAEKVLLGKQKQGDLDKSQGKTLWNAMGLCQNSLGYCMDAVASQQRALDLDPDFRESIVNMAQAYKDFGNFRTADELFTRALKLEPGCVTTLNLHGLCLYGAGEIRRSYDTFCKILEISPHSEDCLFMHGISAQGLGLFRKAAESFSRINGMNPSHWAYYTRELSLFQHHHLDMPMKDYNADIELNECFKEAWTKRLDPTHLTFDYCKQADFSPDIVDVDQNATLSNVARDIAKVAFPLGDKLQYHTPGFLVNRKQHRWFGLAMLDMAQTVRRVWRVMRKQDTGGSIIVTGRASSGCRTSHVFGWRDLFDIAVKWRQASEPNDPVWWVDLLNQETFAEGFGSQTPMILGQCHVVRYYPYWKKAFPLMKDLMKDQCTPSPAQLEQIKEAKTPHDLYVIRSHDFYVATPCHSQAVPGRIMEGTRLTIQKKTRYGYTCSIRTPGTPARWKQFDQEMGHAWEKLCEEAVGPQPDTRTVFELTATLAYYWYNFMPLSRGSAACGLLSLHAVLLAMDLEIGRPMPAGLQVDWEAIFLPTPRDFISHLSPNVKREIENNDLKFGGHQ |
| ApoLp | MGNSAGAKSSPIYLGGCDEPVLRPTRPAEKSKAKKKPDKNEKAKPTTVDNMVSLLNKRHKVYNIKVLSSHSADKNPPRAPDTVALKKTADGMQQLQVGQKTINTWSWDKSTNKLMWSDGDHAGHIAFPHEESARGYGTMSLGQTSYSVMATLSPVTYRCTLSSNAGAYVKQEGSALKLIWDESSSEWKNASWDDNVLSFTYGLEEQFFIGQKTYKVDINFEDLQTNVTYNPAEGDFSFVLTPEFKAVFQHENDTVPVPDDPRDDNKHVFPYLLQFTFNAEASEFTGAMLTKALDAATGEVYAVKAYVQEAEALTVSAPPPLNGQELLNMSQFKKDDDGRWYDAIQQKSMNDFYTILQYYMDADLRKKYMGMNNAPDLDPTIKQIADTDGNPSDEQRENGDDGNAKDWYKTLSVAYLAQALPTVWTNDSYAAKLNTIRAQKWLKTETSTSSVFNAQSPELYKTHWLRDMPRMGDFLVDQMQNNAEYASYIEQDKNSWIEEVESTVEDPTNQASMLSIIESLTTLGKEGKYWAYWLFRDITQPSALSMLQMLSVGGAASLDGSAFSRQIQQNCAILGILDESATFQEQYIRIIQIFQLTNILPQLIDFSGDMTNYQYAVEDILKGFEEQYINSPDPQMQEAAKEIQEKLNQKNLEDLLSELQKMASQFTGDYNFVKLMTYVDNVYAKGTLYLLGKLVASAAVCGAIMMFYFGVKNWKSLDGAKRAQVVSDGVLLFANVAAGLVKRGTALYEIWGTEAMTYKNVAKITFEGECQESIERASNGFSKWILKKKGEEAEVGVEATTEEAVVAGEEDMSLLETLLGSNLDEFMATRFGGAVAVLGLVFGAISLSKSKTDLEKAANSMGVLASTLQICATFGSWLIPEGAMVAGVECAAICSCLGALAVVAAIVGVILLIIELKKKTPDPLKDFATNQANDAGLYMPHGADIDSFQSYQPKGQLEKEGISLQMNGSSQQYLKFNTDGTLSLSPGTNGPDTVFYMTTDSYGRAKFIANLPNADNTAVVSKFLTVDGNGQLSGADQLGGDSANQQLWVAECQGSVQRTDGHLTSASFKFYSFSMFKAKNKKYYLGGSGGAPEVSESAQQPWTVSMDSTKPVGLRMDDITLFTFQMDQKFTPTLLNPGSDPKNWSIRPDLPSFMQLDQATGAVSQKSGVAPEITASNTYTLTVQNDLGSTSTKFVFKVGEYPDGA |
| HintN | MAAIQDPAVEKLFGLLPTPGGNNSARAALVSKLKGSTPGRHTKAFGKKGDTFRKIFFPNYKAAPYEGDTSLTKLKNWWADFSVAVLCQQGMYNLTKDLRKQLKKDDINNAVKSKNSELKTHCMAFYAKVFRDTFAKAQYNSIKNKTSAKAQYKAALTSDAWIMAKRMEASEGMWTDAAWELYHHWVKLHLLGASNAEIDGVIKQLSSKDLMIPPEVGAGKWTSYTAWMDPAAITWKDIQGDAAKGILKSERMPSYSPYGGSSSMKEENSFEFTAKGQPGRGYRHTGGGGGGSCFTGDTKVLMANGTLLPIRSVEVGDEVFTLQGPRRVAVISMPTRKQRHLFSINGHSFRFTETHPFVTPAGLGGGMDVHGAAFAAVSPRKLANLVPTLSRLGIVKMEQGLTVVSLQNEQPHPTPVHLIEEHPLKKPESEEDLLYDLILEPTEAGFPHYVVGDDSILVLVASEVPRIELAPFAAITILKALQEAADHIEAHPLPVNRGEPPSRVPHVFHVLSPTKAMVSNMDAMAINSKTCARQAESCDVSDMLSVFVKEDTASEQTSYRRELGDAYEQIAALHGEEIDAIVDLGFRSIGQEEDDASQWMLSVSLLDVIFHDLPKHSPSQRLAVNIEARTREKTDKVWCEESSEKISSNFARHFRQVFYFPDVDIDDGCVTDIHFQVHDADDSHPLPYSASVVVPWKLEHNYRMFEAILHDEDDQEIGLLRFDVRLLSSNAMQEEQRRSETWEDADKQAYALALGKCYGEHMKVQLPNVLSHGLVAVTKRPWFKRSCFSM |
| SDR_a4 | MEKVLIIGTGFVGSSLAASLKSAGVQVYCTRRQRGEEGTGTSQFVLEEEETWRNLPEEVDGVVWTLPARPLSQVQRFYDVYLQSILDSGKPVIVYGTTGRYHVEEEHQWVDEKTRYKQTESGREEGEDFLQEKGGTVLVLAGIWGESKNGQTRGPKAWIQKGYIEAVDKFLNLSHVDDIVKATEVVMERPELKGQAINVCDGNPMLWRDIILGLGYELPPMTTTMTAISSRPSKRVSNNLLQSIMGKDYKFQQVMDPVDLAVD |
| CytP460 | MISSGFGRVVAFTGAAGAVVGVVAGFRRRQAEFQAAALREFSRPSYNPPPAAYTGPVFKPRLDFPTRANPNNETFPWLDIDFRTEPEKYLYTVRDYCLEGNVECDFKVDQNQTRDWYHVPWMSSQPTGREPVHGLTMEKSSKQGMLSESQRREVQNWAVAFYNSPGAHAVGQVWSLPWQPAQDGVTFPEGTVGFKPLFTQATAEEVPFLKGSPVWKAAIAKTPREPGDRGPPVETRLIQMDVAVRDDRADIGWVFGTFVYHSFQCDDAPWRRLVPVCLQWGNDPDLTQQRYQEGARPVQTWNNPRIRQLGILSGSRPYLGWLGRANGPVDNFKSCCASCHSTATVPDKDNSVPRGVPPKNATADQTLQWFRNLAPGQAFQKDGKSLDYSLQLSSSISQYNQWKRSYLKNTLFGRLKDWWQEANPFQTVPPSGKDD |
| UF-TM1 | MRPKPILPRIDDCECTPSVQHLFWRHYLLQSPMYYIRWIYAALYLLYLLFVMRVPTDRDIVGYIENTTMAMLIRPATDGKSGEYEVTVRDCKLRALGGYKLKSMSLRYKGGKSGVEVLCFTRNGVKIDNRYQIFSTIYFYHIHSIHTKSHLFSNSLVRHIVDNDVKILQESSYTSIPLHYGLLHSSVCALAWDGNVSRYLGYGHVGTRESLVEESRNMSTLAGHQTMEHWKSHGKDSFAGKLYRSRLALQNVMERHKIDPKLLDPLLTTPLFSLHPWDTECSIYQAFNRSMFRILITQPNLNPLAPNTIRSINKPFYHDLYRELRKIDPKMADVVTASVMF |
| UF-TM2 | MKGCLSALFVSLCVVCASGTSAPHTTEASLVSKLVSGIPTDRQPTVHVGHGHNLTTETASLPTASSAETTTDRPVSTTTKPPLQTSTSKTGSADASGPGSGLGVVRHHRPLTLFGQLGAPLASAPQTRDPSCPPVNGSEPPQLVTSDGFYKKVTEDRIEILATDVALDETSALFDPCKDFYLETVSLTIKGHLRWRGRNVYISAVNLTVAGDSAAVDVSYIAHGQPDWDTPAQSGTAQGQAGQDGQAGGAGQRAGNATLYVGFLFGGRLEVLARGGSGGQGQAGGDGHVGAAAQPNQDRDACCHGRSTAGTTGGKGGDGGKPGQSGDGGPGGVITIRLLNKTWEDLDIHNLQALQLPTADVSGGPAGPQAAAGKPGPGGPGGGGSGTHCYNAGHWYDHHWRCEGPYAPPGPQGPPGQASVDVIDFEDISEHISPEMVQMTFRQGNRYYKEGDVEAARNVYLWVYFTMQNSSRPELRGHAREAMIYLMQIQHGLDFWAHAQNNIPMNSYSSIYAYLSTDIVPYAIDVEKDYNLYFAEATSEEQKMVVAKSSMDHNKFTITVLEKQKMEKLDELTQIQSELRDLEQQQAVRLARAQDATAECQDAIEADLVHKATAIHFSDILKIAGAAIKIYTSIYTGGLTLGSGINDIISALEAGKKLEGLDFLDQLKGVKTIIKNICGIWDKDKGIIKSDLKDIEEQYDKIKDTLDHKGQENQQKIVVDGKKFDEQFDKWLYLAQCRDAKPRIQAYLNVTQAVNDKLVHHDAVVIKIHSLDAEIQVLQQQNARLRSETAAVFDPTIISYAMAIGRVYVQLKDSIVDQMKRLQEAYNYQFLERRPFSYDDSRMAMIEAWLANNRISMLQRMEKIGNDVQVFNADSSPMTNTIVLRRPDLADDFAQFDNGTELPFVITGAEDLLMPMTHVHMTDVKAWIPGVRTEDRHVKVWLKRYGTSLVYDQQGGMWTFTHAPRTMLFHYNTGSLQASSDYVTLSPIGPWTLAVPREYNPGVDVSRVREIHLQLTLTFLPCAQPVCPPRTRQVNFTDAGAGSTAEFRVQDTAQVVGAGSAAWVAVLVVLAVLFCGLVVGIVYVRKRRSGQSPPGHSETTPLTANTAV |
| UF-TM3 | MAQANALFWRQVRPLQTWRLAAAATTSLRLRSGHRRWMSTDLVTTKDKEWFHWKDLPWQAISIGVGIGTLLYQTRKNKELELRRNNLEYVNSQLSQLYGPLYGNRLANQRSYKEVLQGHDNLVKFLRVAERKWRNPETGEEGVRLLTRWRKFLFYVMHPLDLKAEEIIRDNAHLFEHGVEEAVLFRKFVFHVNYQKMIVANWQEKGEVLGNKEVFEEKDFSRENNAGKSDEQTFKMLVQVVDHVKKTYEKLVERKKSLVQEIKERGGH |

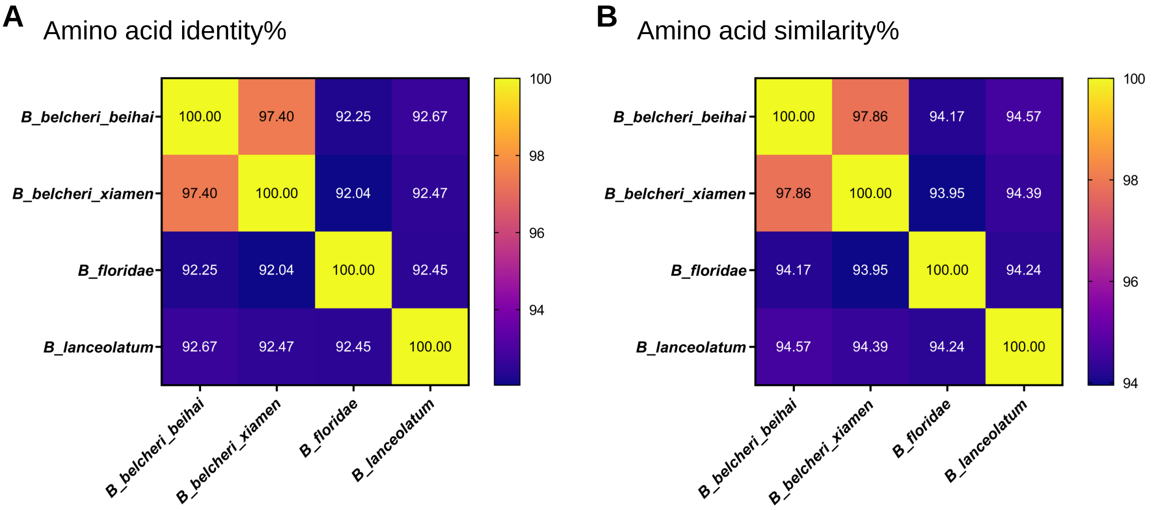


**Fig. S1. Similarity matrices of four amphioxus genomes**

One-versus-one matrices of the amino acid identity **(A)** and similarity **(B)** percentages of four amphioxus genomes. The genome assemblies of four amphioxus were assessed by BUSCO v3.1.0 with database metazoa_odb9. Then 295,426 conserved residues in 834 overlapped BUSCOs of four amphioxus were used to generate the similarity matrices. The identity and similarity percentages were calculated by an online tool SIAS (Sequence Identity And Similarity, http://imed.med.ucm.es/Tools/sias.html) in default parameters and the heatmaps were finally edited in GraphPad Prism v9.0.0.

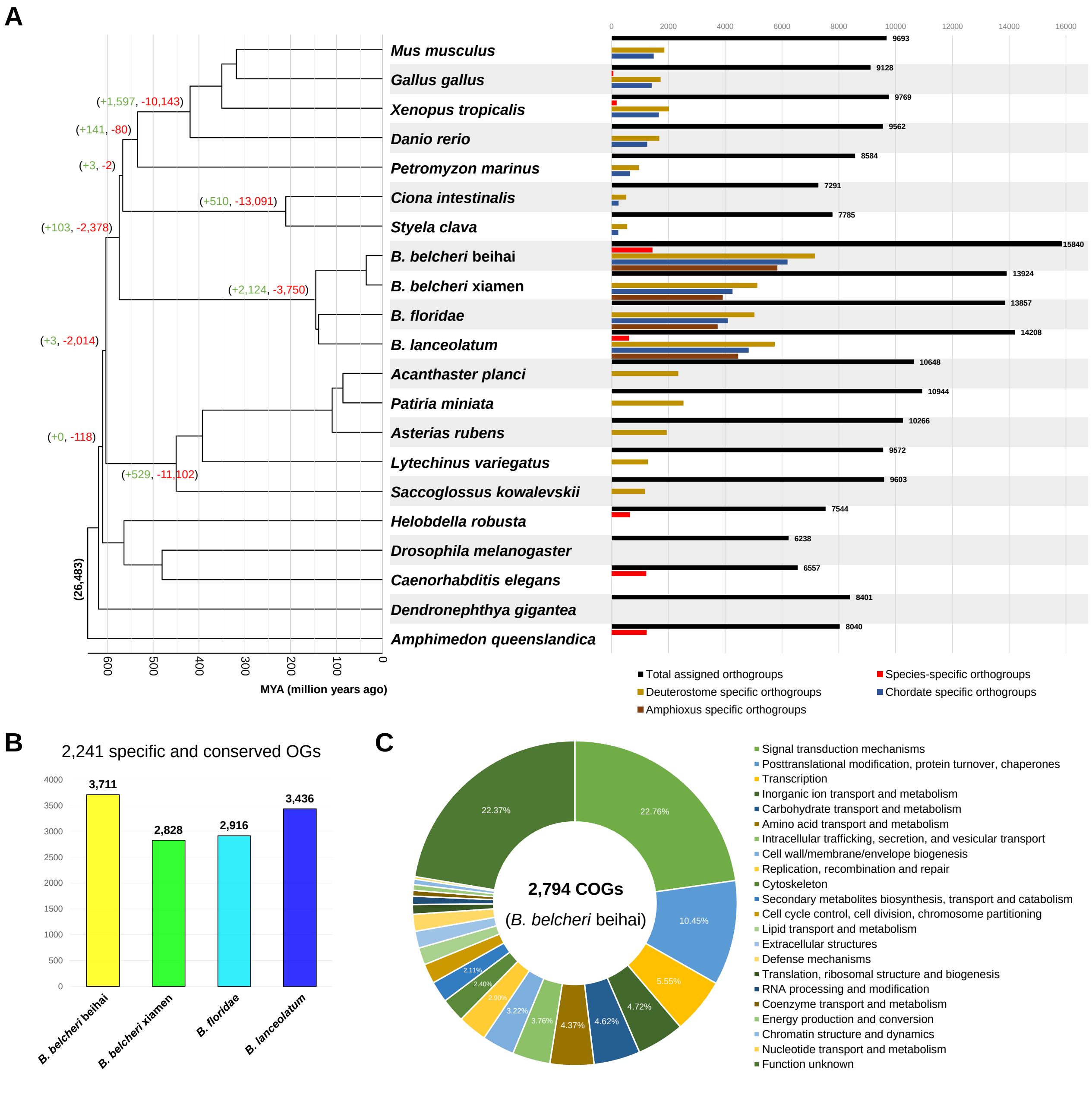


**Fig. S2. Phylogenomic orthology analysis of major deuterostome groups**
**(A)** The protein sequences of 21 annotated genomes were assigned into orthogroups (OGs) by OrthoFinder v2.5.4. The number of specific orthogroups were listed at different levels. The ultrametric time tree was constructed by MCMCTree in PAML v4.9j and final edited by the online tool iTOL. The numbers in brackets on nodes indicate the number of orthogroups under expansion (+ and green) or contraction (- and red). **(B)** Number of Genes contained in the 2,241 specific and conserved orthogroups of amphioxus. The 2,241 orthogroups were specific in four amphioxus and contained at least one gene of all the four annotated amphioxus genomes. **(C)** Functional clusters of orthologous groups (COGs) of the 3,711 genes of *B. belcheri* beihai in the 2,241 conserved OGs specific in the four amphioxus genomes. The COG category was annotated by eggnog-mapper v2.1.5.

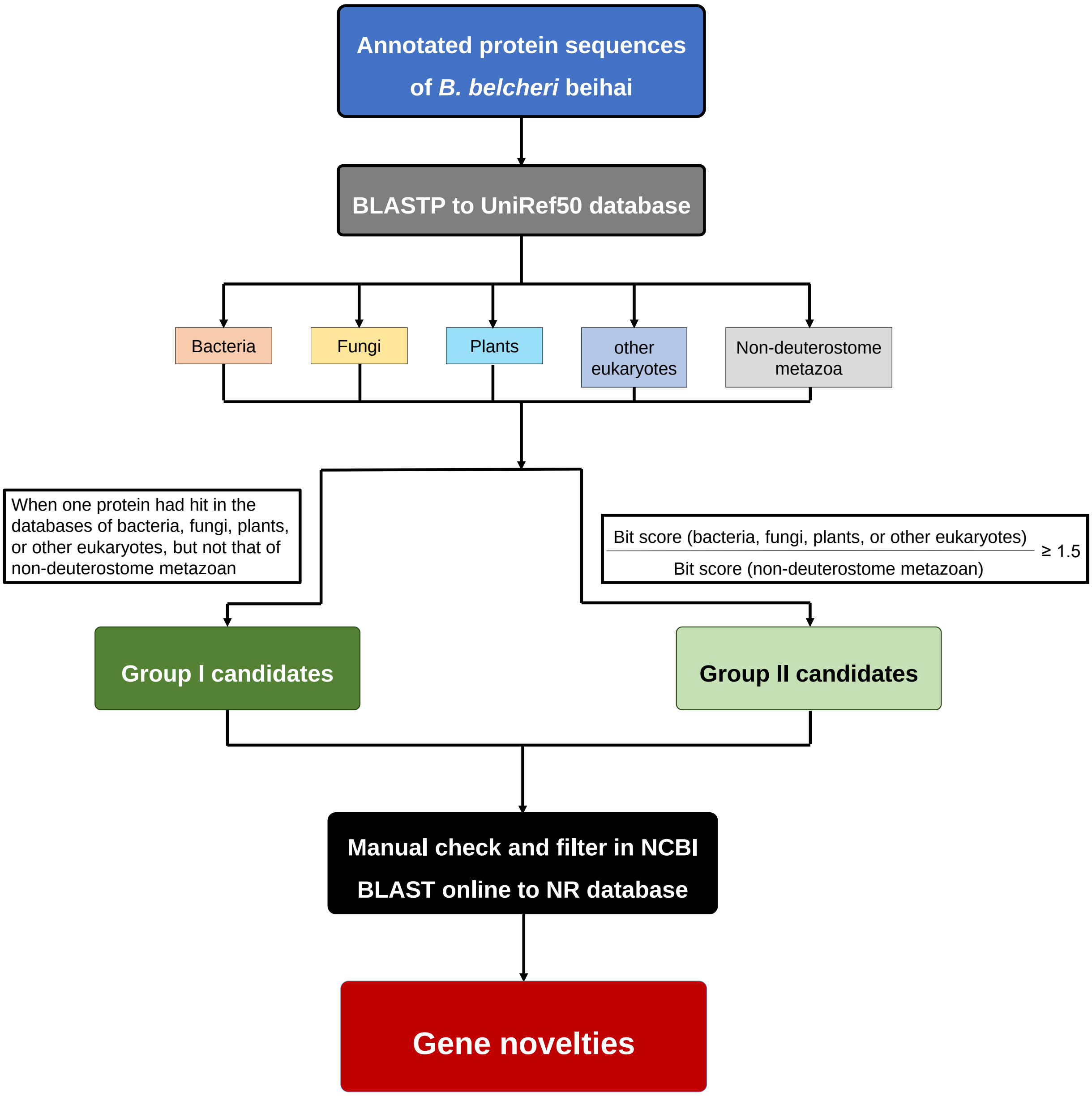


**Fig. S3. The identification workflow of the gene novelties in amphioxus**
To identify gene novelties in amphioxus, we analyzed 44,745 annotated protein sequences from the genome assembly of *B. belcheri beihai* against the UniRef50 database, spanning five distinct taxonomic groups, using BLASTP at E-value cutoff of 1e-6. The five taxonomic groups are non-deuterostome metazoa, bacteria, fungi, plants, and other eukaryotes (excluding fungi, plants, and metazoa). Following the established criteria, we identified two groups of gene novelty candidates, which were then meticulously checked and filtered using the NCBI BLAST online tool against the NR database. This identification method was adapted from the strategy previously applied to detect horizontal gene transfer events (Xiong et al., 2022). During the manual checking and filtering, we first eliminated candidates whose unexpected sequence similarities could be attributed to low-quality regions, such as repetitive sequences. Next, to prevent overlap with prior studies on gene novelties in stem chordates (Holland et al., 2008) and deuterostomes (Simakov et al., 2015), we excluded candidate genes that exhibited high similarity to genes found in jawed vertebrates. The final list of gene novelties was subject to further comparative analysis.

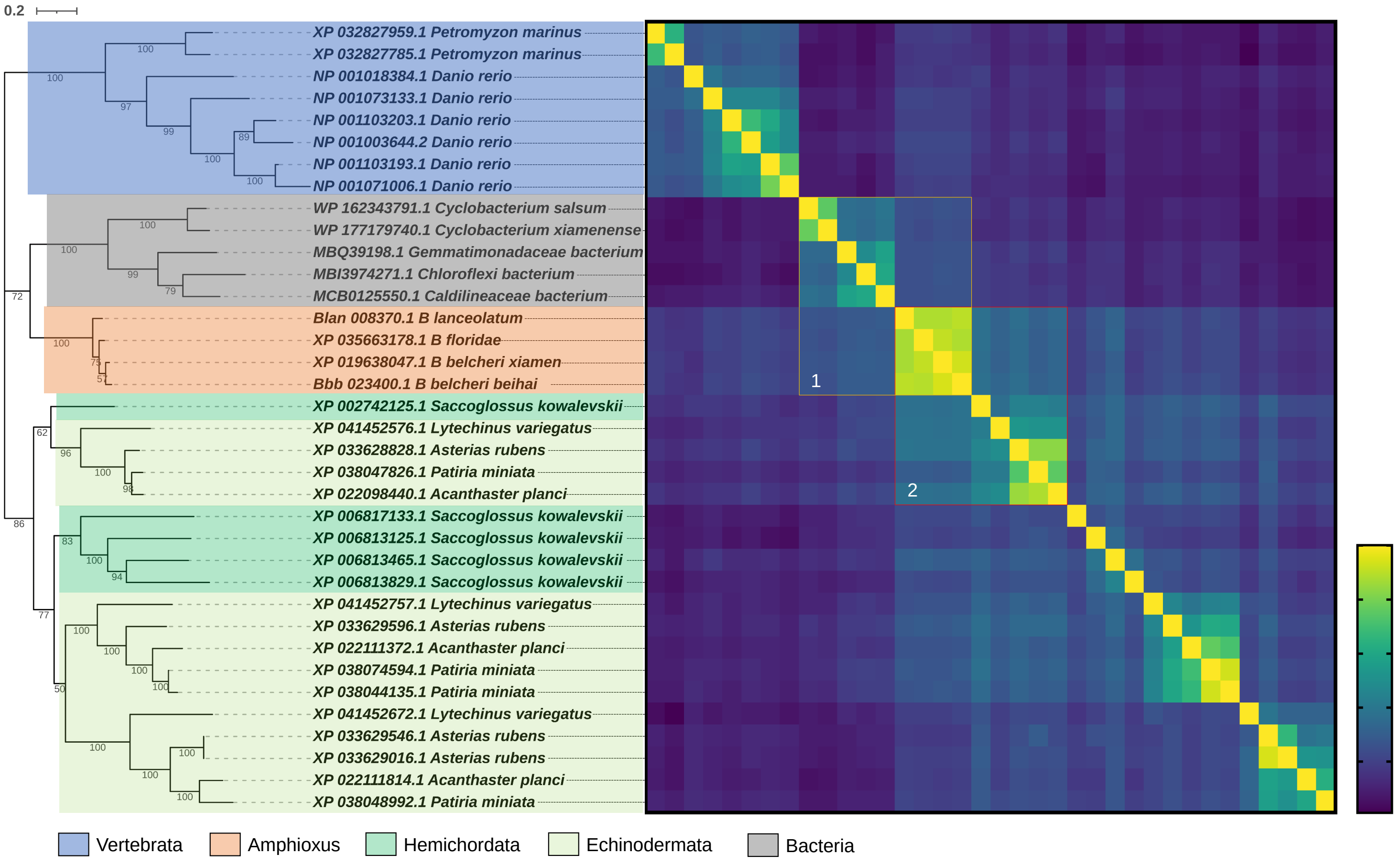


**Fig. S4. Phylogenetic analysis of the sialidases of amphioxus**

A single-copy sialidase of four amphioxus were phylogenetically analyzed with the best-match proteins in other taxonomic groups. Except two genes of *B. lanceolatum* and *B. belcheri* beihai were newly annotated in this study, all genes could be found in NCBI NR database according to the leave ID of this phylogenetic tree. The heatmap described the similarity matrix (BLOSUM62) generated by an online tool SIAS (Sequence Identity And Similarity, http://imed.med.ucm.es/Tools/sias.html) in default parameters and was finally edited in GraphPad Prism v9.0.0. Two high-similarity blocks containing the sialidases of amphioxus were highlighted and labeled as 1 and 2.

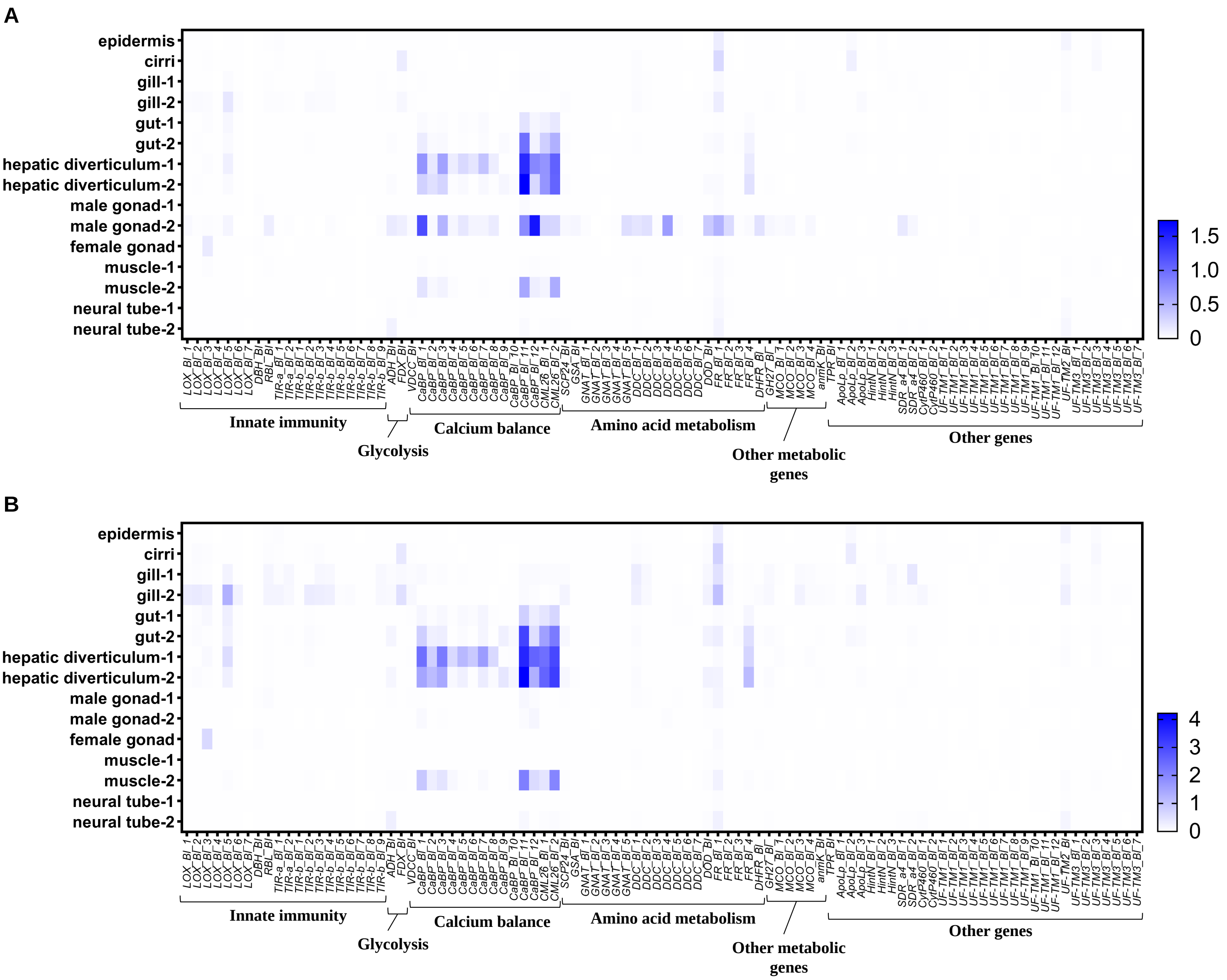


**Fig. S5. Heatmap of expression level of the novel genes in different tissues of *B. lanceolatum***

The transcripts per million (TPM) values of the novel genes (Table 1) were firstly divided by that of the Glyceraldehyde 3-phosphate dehydrogenase (*GAPDH*) or elongation factor 1 alpha (*EF1A*) to generate the normalized TPM values, and then the log2 (1+ normalized TPM) values were calculated and presented in the two heatmaps, **(A)** and **(B)** for normalization by *GAPDH* and *EF1A*, respectively. The expression level of the genes (Table 1) in different tissues of *B. lanceolatum* was generated by the program Salmon v0.12.0 with the options '--type quasi -k 31' using transcriptome data (Supplementary Note 3.2). The heatmaps were finally edited in GraphPad Prism v9.0.0.

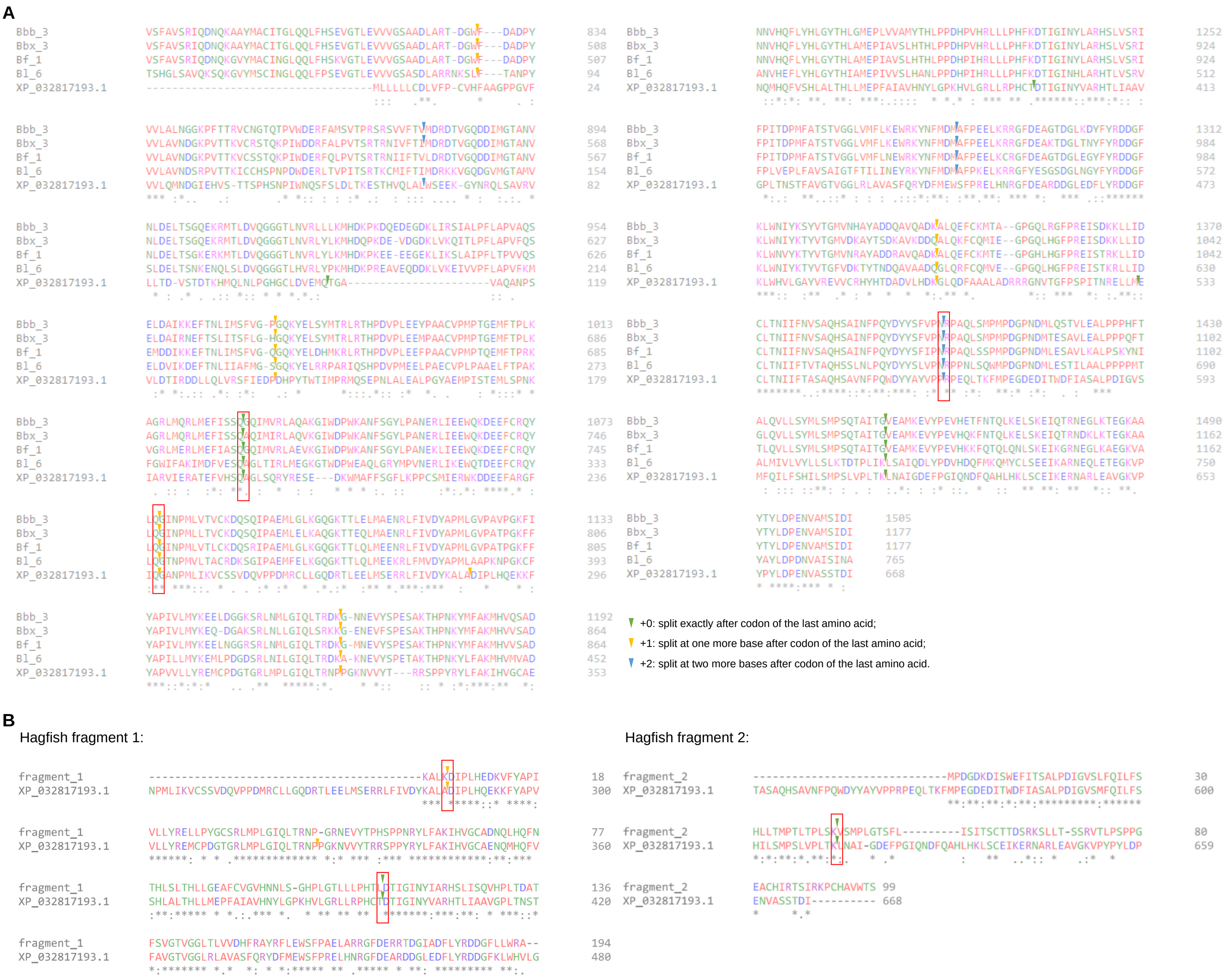


**Fig. S6. Splice site alignment of the LOX genes**

The protein sequences of LOX genes from four amphioxus, the sea lamprey *Petromyzon marinus* (GenBank accession: XP_032817193.1) and the hagfish *Eptatretus burger* (manually curated fragments with transcriptome reads) were aligned by the online tool Clustal Omega. The splice sites (locations of exon–intron boundaries) were marked by colored triangles, of which the conserved sites were defined with at least one of two flanking amino acid identical and highlighted in red squares.

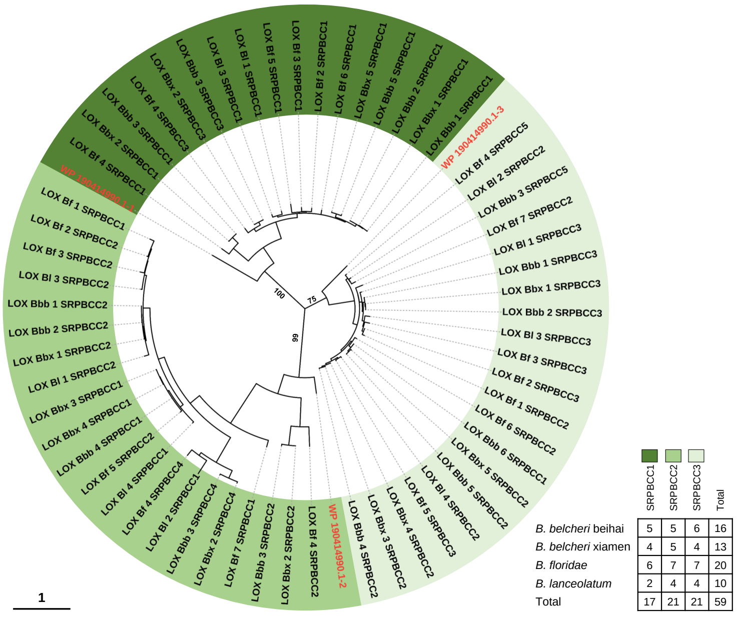


| | SRPBCC1 | SRPBCC2 | SRPBCC3 | Total |
|---|---|---|---|---|
| *B. belcheri* beihai | 5 | 5 | 6 | 16 |
| *B. belcheri* xiamen | 4 | 5 | 4 | 13 |
| *B. floridae* | 6 | 7 | 7 | 20 |
| *B. lanceolatum* | 2 | 4 | 4 | 10 |
| Total | 17 | 21 | 21 | 59 |

**Fig. S7. Phylogenetic analysis of SRPBCC domains in the LOX genes of amphioxus**
SRPBCC domains (Fig. 2B) were numbered from N to C terminal in 1-5. Totally 59 SRPBCC domains: 17 SRPBCC1 (Bet_v1-like), 21 SRPBCC2 (PYR_PYL_RCAR_like) and 21 SRPBCC3 (PYR_PYL_RCAR_like). Three SRPBCC domains (from N to C terminal in 1-3) of the cyanobacterium, *Coleofasciculus sp. FACHB-125* (GenBank accession: WP_190414990.1) were used as outgroups.

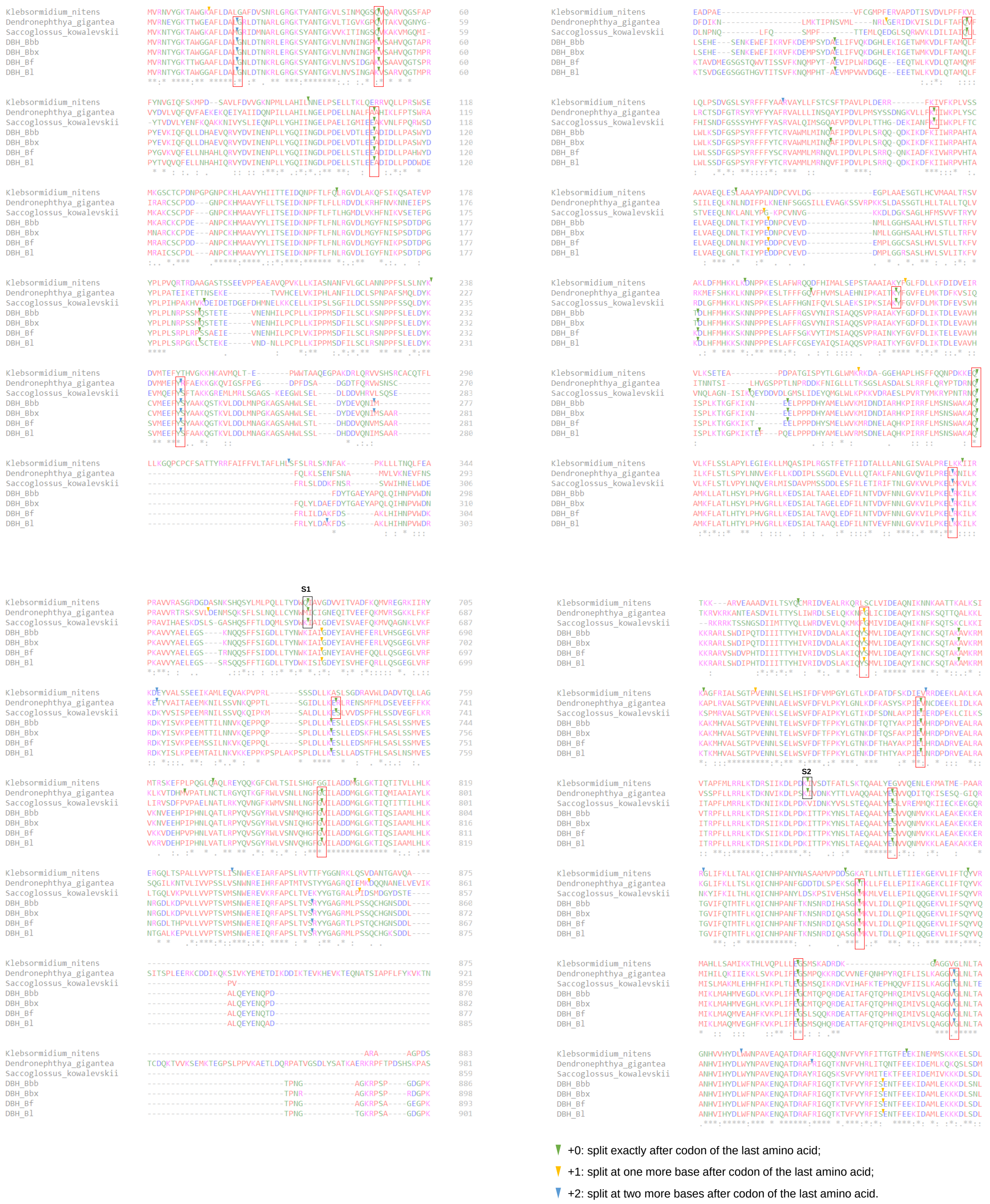


**Fig. S8. Splice site alignment of the DBH genes**

The protein sequences of DBH genes from four amphioxus, the charophytic algae, *Klebsormidium nitens* (GenBank accession: GAQ83283.1), the soft coral *Dendronephthya gigantea* (GenBank accession: XP_028406807.1) and the hemichordate *Saccoglossus kowalevskii* (GenBank accession: XP_002737002.1) were aligned by the online tool Clustal Omega. Their splice sites (locations of exon–intron boundaries) were marked, of which the conserved sites were defined with at least one of two flanking amino acid identical and highlighted in red squares.

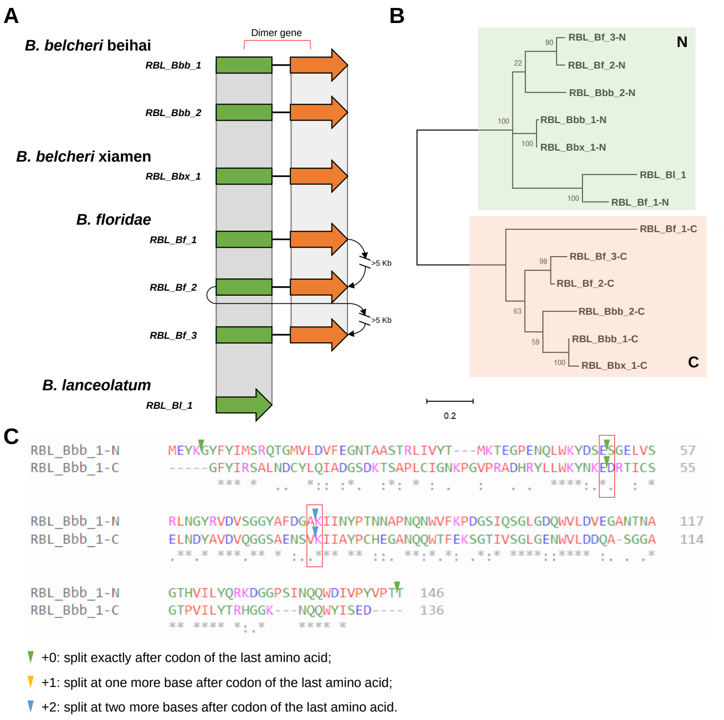


**Fig. S9. Ricin B-like lectins of amphioxus**

**(A)** Alignment of N and C terminal domains of dimer ricin B-like lectin genes. The distances between proximally arrayed genes of *B. floridae* were 5-10 Kb (labeled as >5 Kb) and other gene distances were all below 5 Kb. **(B)** Phylogenetic analysis of N and C domains of ricin B-like lectin genes. **(C)** Protein sequence alignment of the N and C terminal domains of *RBL_Bbb_1*. Two conserved splice sites were marked in red squares.

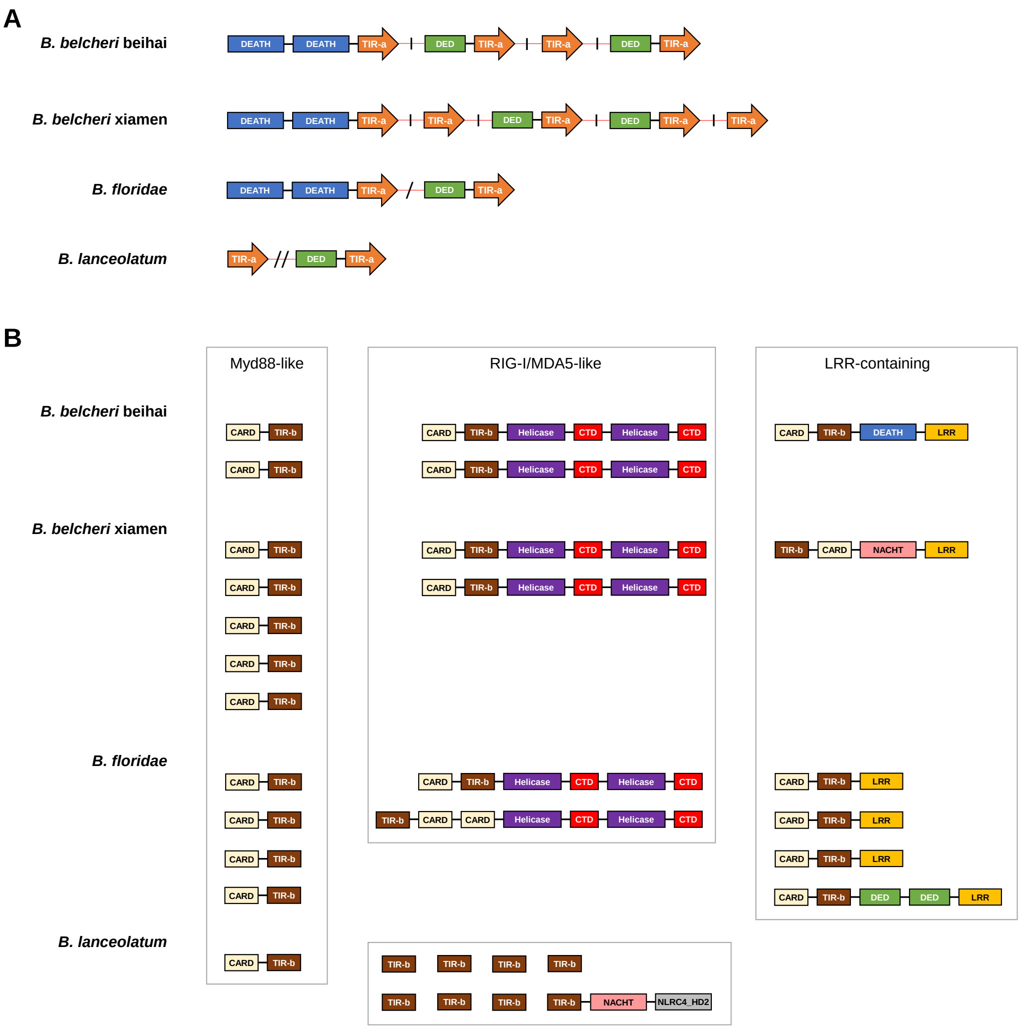


**Fig. S10. Schematic summary and domain structure of two groups of TIR-containing genes of amphioxus**

**(A)** All TIR-a domain-containing proteins were analyzed in gene synteny and domain structure. A vertical bar (|) between genes represents distance of below 5 Kb, a slash (/) represents 5-10 Kb and two slashes represent 10-15 Kb. **(B)** Partial TIR-b domain-containing proteins were analyzed in domain structure. Abbreviation: TIR, Toll/interleukin-1 receptor main; CARD, caspase activation and recruitment domain; Helicase, DEAD/DEAH box helicase; CTD, C-terminal domain; DEATH, death domain; DED, death effector domain; LRR, leucine-rich repeats; NLRC4_HD2, NLRC4 helical domain HD2.

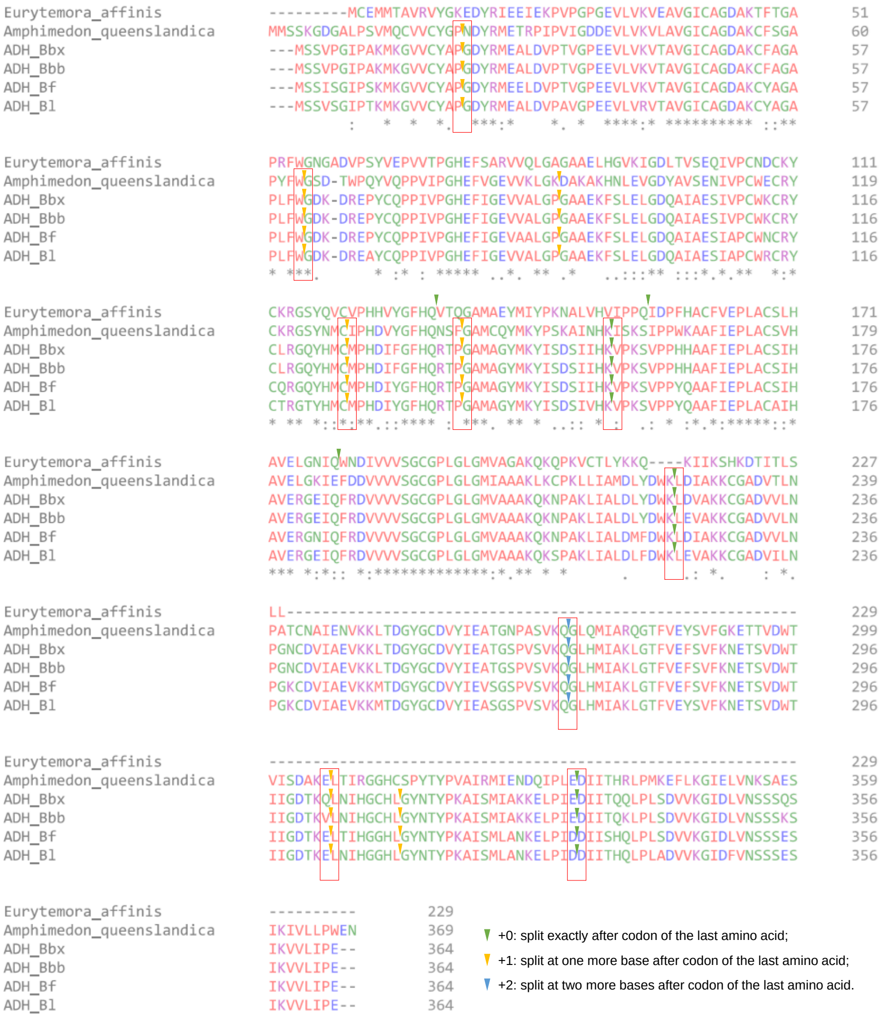


**Fig. S11. Splice site alignment of the ADH genes**

The protein sequences of alcohol dehydrogenase (ADH) genes from amphioxus, the sponge *Amphimedon queenslandica* (GenBank accession: XP_011406967.2) and the copepod *Eurytemora affinis* (UniProt accession: UPI000C762174) were aligned by the online tool Clustal Omega. Their splice sites (locations of exon–intron boundaries) were marked, of which the conserved sites were defined with at least one of two flanking amino acid identical and highlighted in red squares.

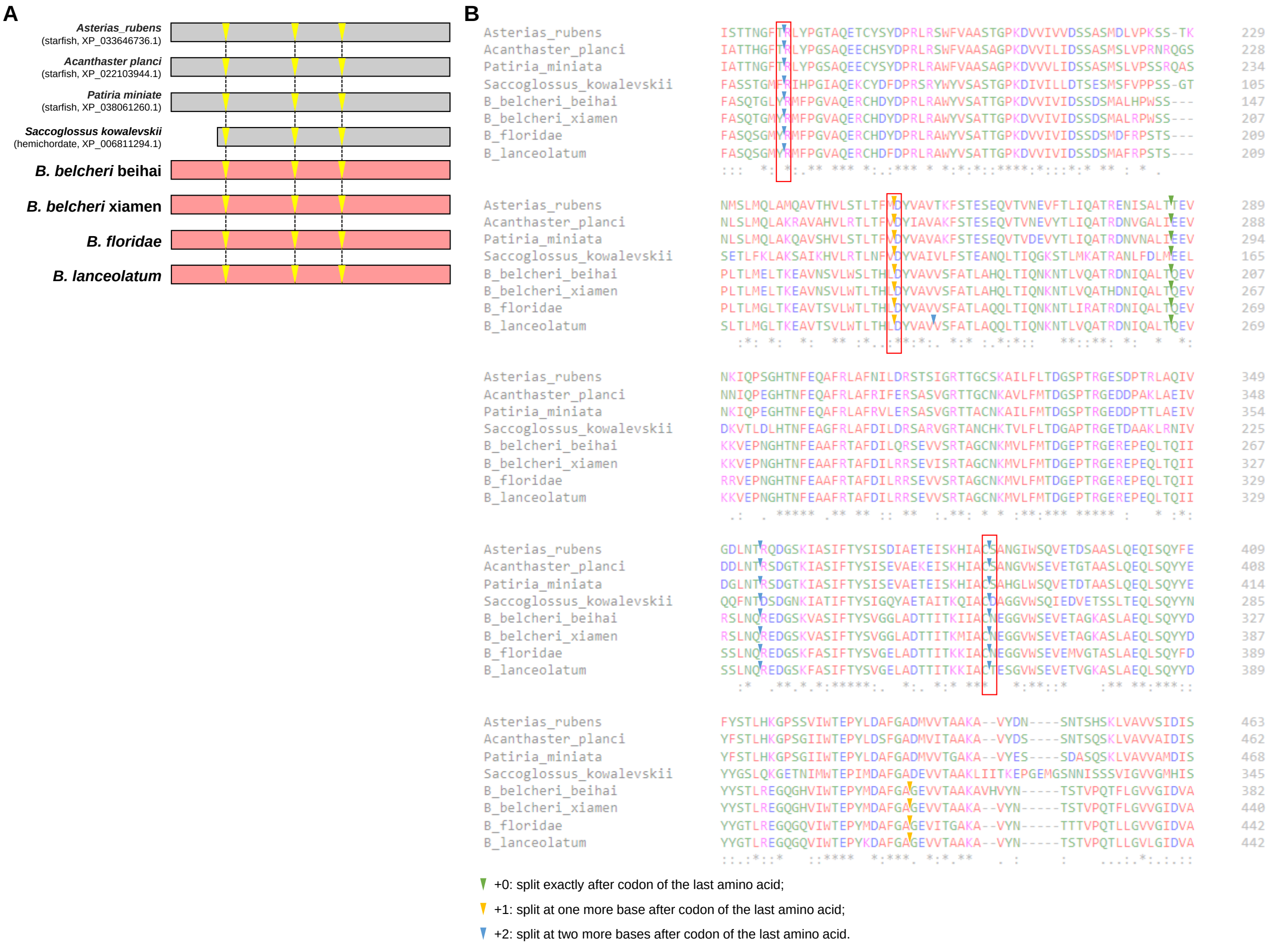


**Fig. S12. Splice site alignment of the VDCC genes**

**(A)** Alignment of splice sites of the VDCC genes of amphioxus, the hemichordate *Saccoglossus kowalevskii* (GenBank accession: XP_006811294.1) and three starfishes (GenBank accessions: XP_033646736.1, XP_022103944.1 and XP_038061260.1) **(B)** The protein sequences were aligned by the online tool Clustal Omega. Their splice sites (locations of exon–intron boundaries) were marked, of which the conserved sites were defined with at least one of two flanking amino acid identical and highlighted in red squares.

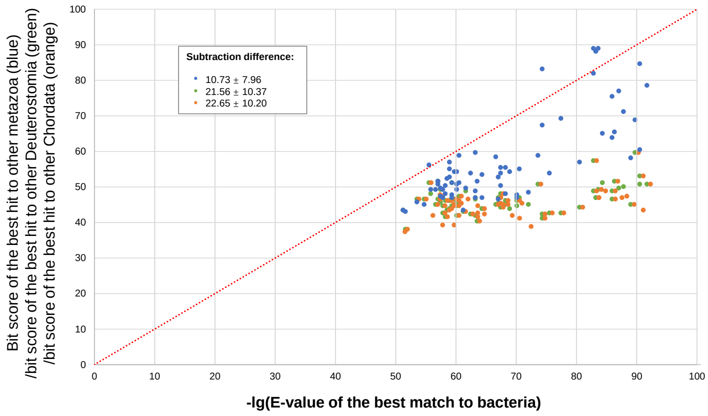


**Fig. S13. Bit scores comparison of the EF-CaBPs in different taxonomic groups**

Of a group of EF-CaBP genes of amphioxus, bit scores of the best hits in the UniRef50 database other metazoan, other Deuterostomia and other Chordata (excluding Amphioxiformes respectively) were compared with those to bacteria in dot plot, and marked in blue, green and orange dots, respectively. Only five blue dots located at upper left of the red dotted line (y=x). The subtraction differences were summarized in the black box (mean ± standard deviation).

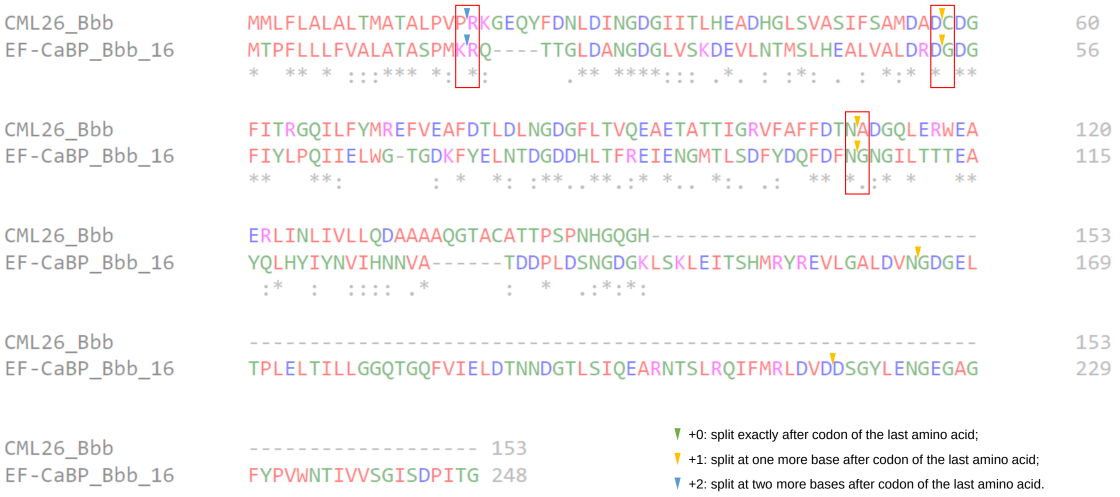


**Fig. S14. Splice site alignment of the CML26 and the EF-CaBP of *B. belcheri* beihai**
The protein sequences of the calmodulin-like protein 26 (CML26) gene, *CML26_Bbb*, and the EF-CaBP gene, *EF-CaBP_Bbb_16* were aligned by the online tool Clustal Omega. Three conserved splice sites were marked in red squares.

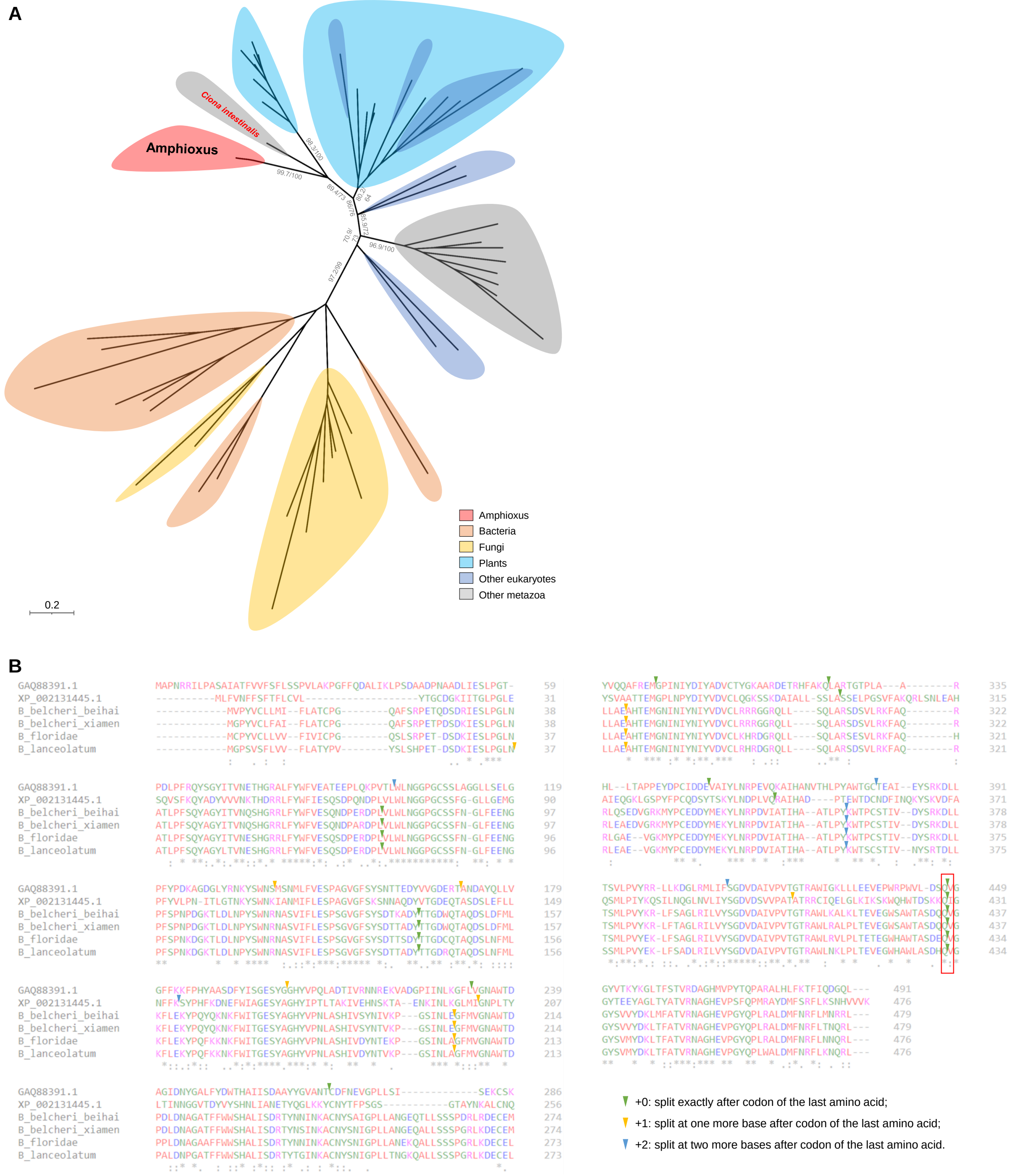


**Fig. S15. Comparative analysis of the SCP24 in amphioxus**

**(A)** Phylogenetic analysis of the serine carboxypeptidase 24 (SCP24) genes. The amphioxus SCP24 genes in amphioxus and their closest genes in the UniRef50 database from other taxonomic categories, including bacteria, fungi, plants, other eukaryotes (excluding metazoa, plants, and fungi) and other metazoa (excluding Amphioxiformes) were collected for phylogenetic analysis. In this maximum likelihood phylogenomic tree, SH-aLRT and ultrafast bootstrap support values are given in that order on the branches. **(B)** Splice site alignment of the SCP24 genes. The protein sequences of SCP24 genes from amphioxus, the tunicate *Ciona intestinalis* (GenBank accession: XP_002131445.1) and the charophyte alga *Klebsormidium nitens* (GenBank accession: GAQ88391.1) were aligned by the online tool Clustal Omega. Their splice sites (locations of exon–intron boundaries) were marked, of which the conserved site was defined with at least one of two flanking amino acid identical and highlighted in red squares.

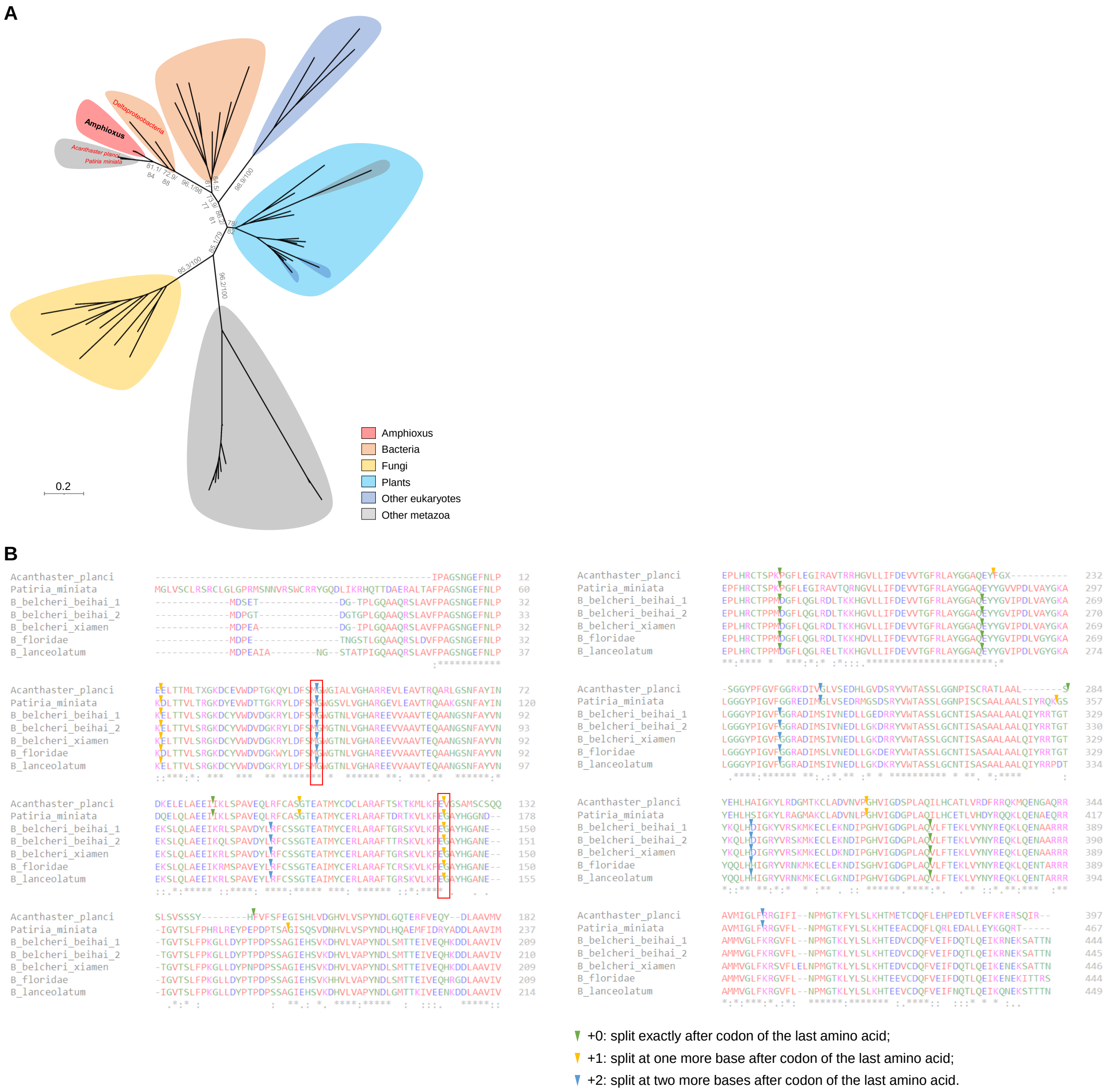


**Fig. S16. Comparative analysis of the GSAs in amphioxus**

**(A)** Phylogenetic analysis of the glutamate-1-semialdehyde aminotransferases (GSAs). The amphioxus GSA and their closest genes in the UniRef50 database from other taxonomic categories, including bacteria, fungi, plants, other eukaryotes (excluding metazoa, plants, and fungi) and other metazoa (excluding Amphioxiformes) were collected for phylogenetic analysis. In this maximum likelihood phylogenomic tree, SH-aLRT and ultrafast bootstrap support values are given in that order on the branches. **(B)** Splice site alignment of the GSA genes. The protein sequences of GSA genes from amphioxus and two starfishes (GenBank accession: GAQ88391.1 for *Acanthaster planci* and XP_038059025.1 for *Patiria miniata*) were aligned by the online tool Clustal Omega. Their splice sites (locations of exon–intron boundaries) were marked, of which the conserved site was highlighted in red squares.

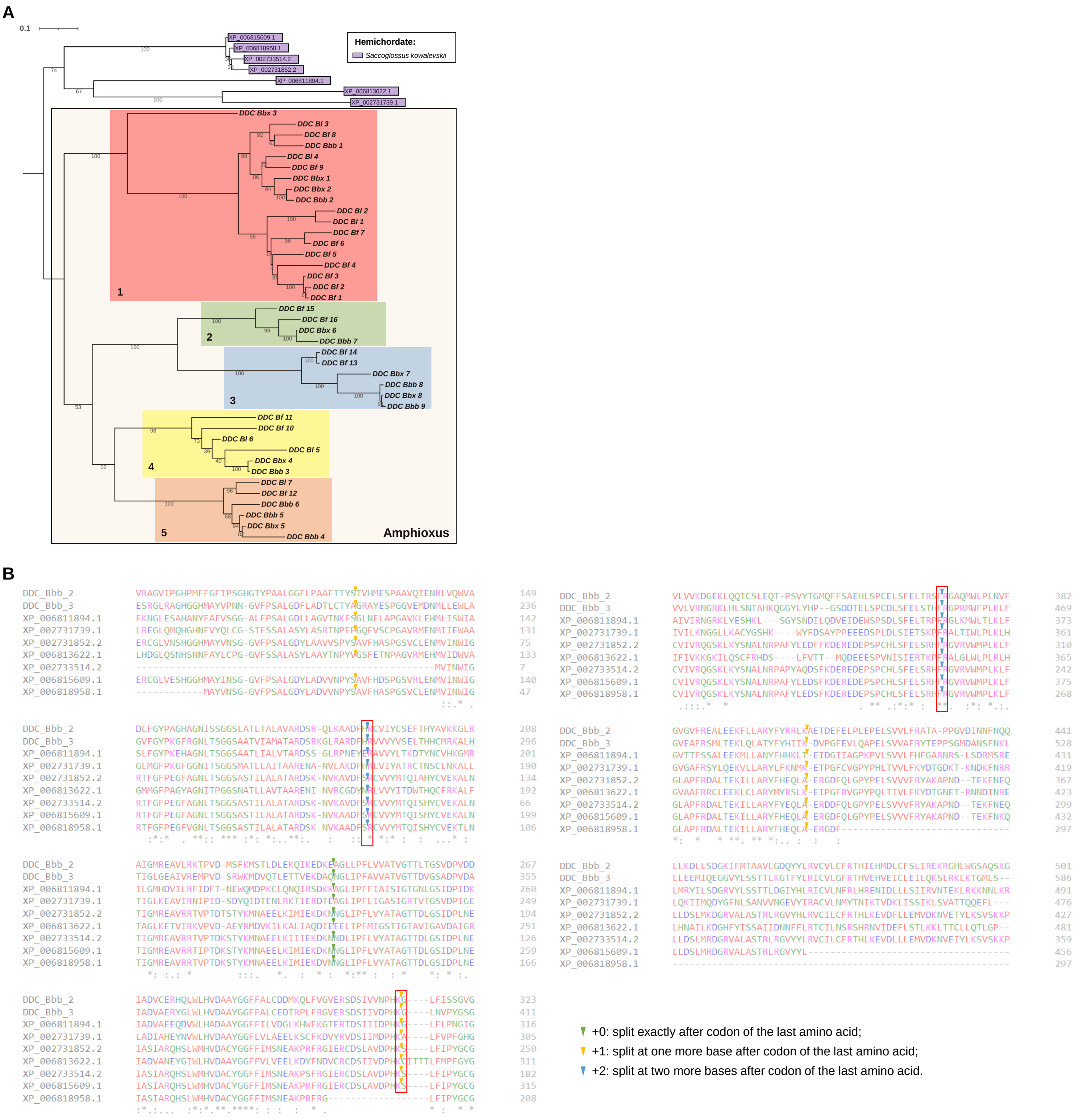


**Fig. S17. The duplicated DDCs in amphioxus**

**(A)** Phylogenetic analysis of the DOPA decarboxylases (DDCs) of amphioxus and the hemichordate *Saccoglossus kowalevskii*. Five clusters of DDCs in amphioxus were classified and marked in different colors. **(B)** Splice site alignment of the DDC genes. The protein sequences of DDCs from amphioxus (represented by *DDC_Bbb_2* and *DDC_Bbb_3*) and the hemichordate were aligned by the online tool Clustal Omega. Their splice sites (locations of exon–intron boundaries) were marked, of which the conserved site was highlighted in red squares.

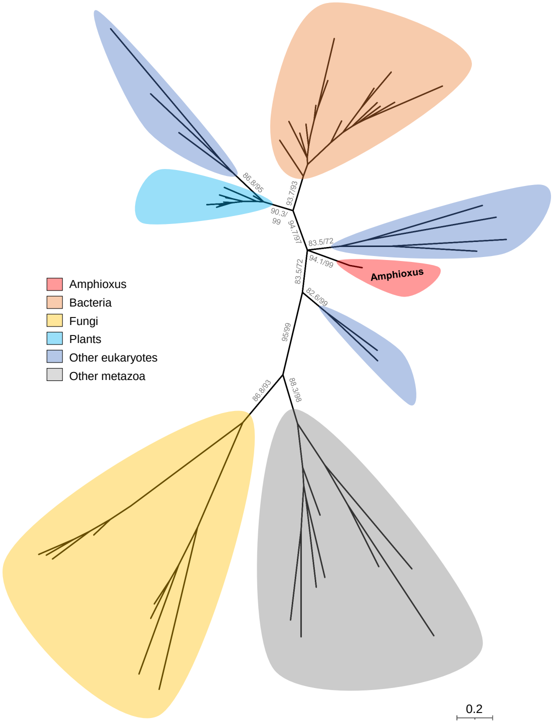


**Fig. S18. Comparative analysis of the GH27s of amphioxus**
The glycoside hydrolase family 27 (GH27) genes gained of four amphioxus and their closest genes in the UniRef50 database from other taxonomic categories, including bacteria, fungi, plants, other eukaryotes (excluding metazoa, plants, and fungi) and other metazoa (excluding Amphioxiformes) were collected for phylogenetic analysis. In this maximum likelihood phylogenomic tree, SH-aLRT and ultrafast bootstrap support values are given in that order on the branches.

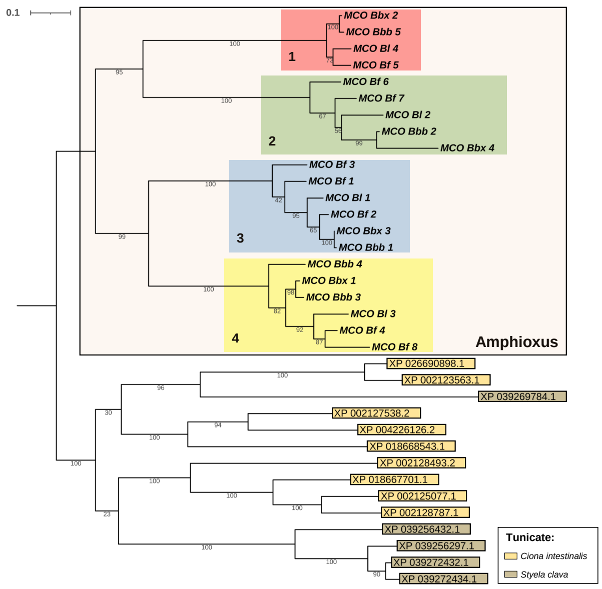


**Fig. S19. The duplicated MCOs in amphioxus**

Phylogenetic analysis of the multicopper oxidases (MCOs) of amphioxus and two tunicates. Four clusters of MCOs in amphioxus were classified and marked in different colors. No conserved splice site was identified.

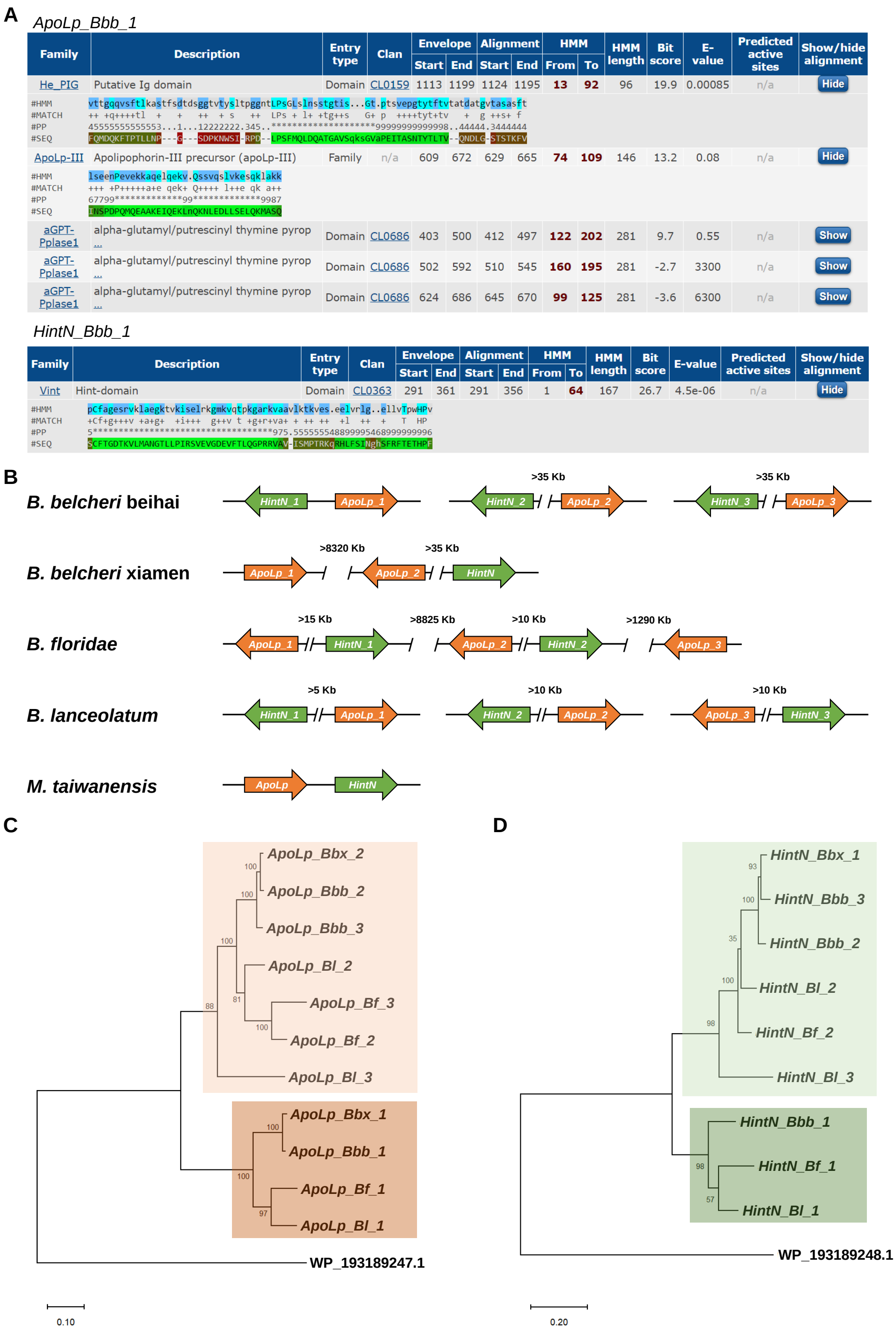


**Fig. S20. ApoLps and HintNs in four amphioxus genomes**

**(A)** Pfam search results of *ApoLp_Bbb_1* and *HintN_Bbb_1*. **(B)** Gene synteny alignment of ApoLps and HintNs in four amphioxus genomes. The distance precision was set as 5 Kb, e.g., >35 Kb stand for 35-40 Kb. Phylogenetic tree of all ApoLps **(C)** and HintNs **(D)** in four amphioxus genomes.

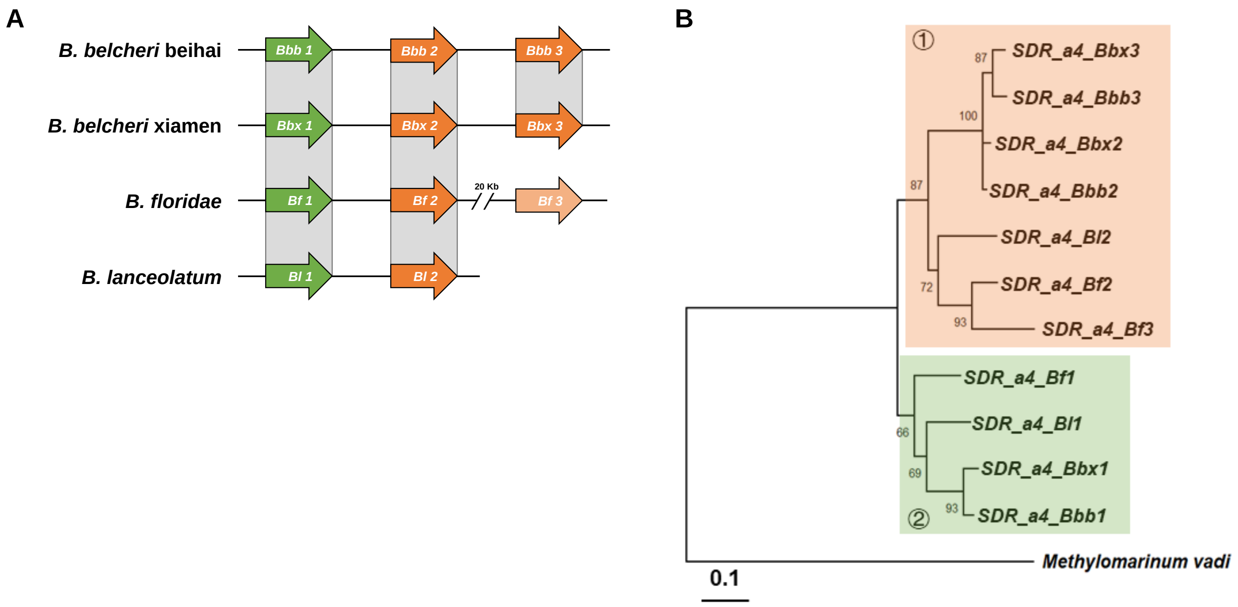


**Fig. S21. SDR_a4 of four amphioxus**

**(A)** Gene synteny alignment of atypical short-chain dehydrogenase/reductase subgroup 4 (SDR_a4) genes in four amphioxus. Except the Bf_2 and Bf_3 genes were 20-25 Kb apart; all other gene distances were below 5 Kb. **(B)** Phylogenetic analysis of the SDR_a4 genes in amphioxus. Two distinct clusters were divided. The closest gene in UniRef50 database (UPI0004DF68CE) of the marine bacterium *Methylomarinum vadi*, was used as the outgroup.

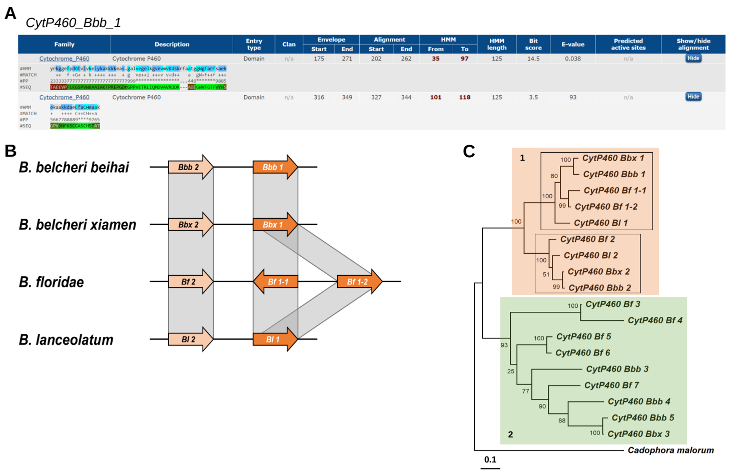


**Fig. S22. Comparative analysis of CytP460-like genes of amphioxus**
**(A)** Pfam search results of *CytP460_Bbb_1* . **(B)** Partial gene synteny alignment of CytP460-like genes. **(C)** Phylogenetic analysis of CytP460-like genes in amphioxus. Two clusters were divided and gene synteny of CytP460 genes in the cluster 1 was aligned in **(B)**.

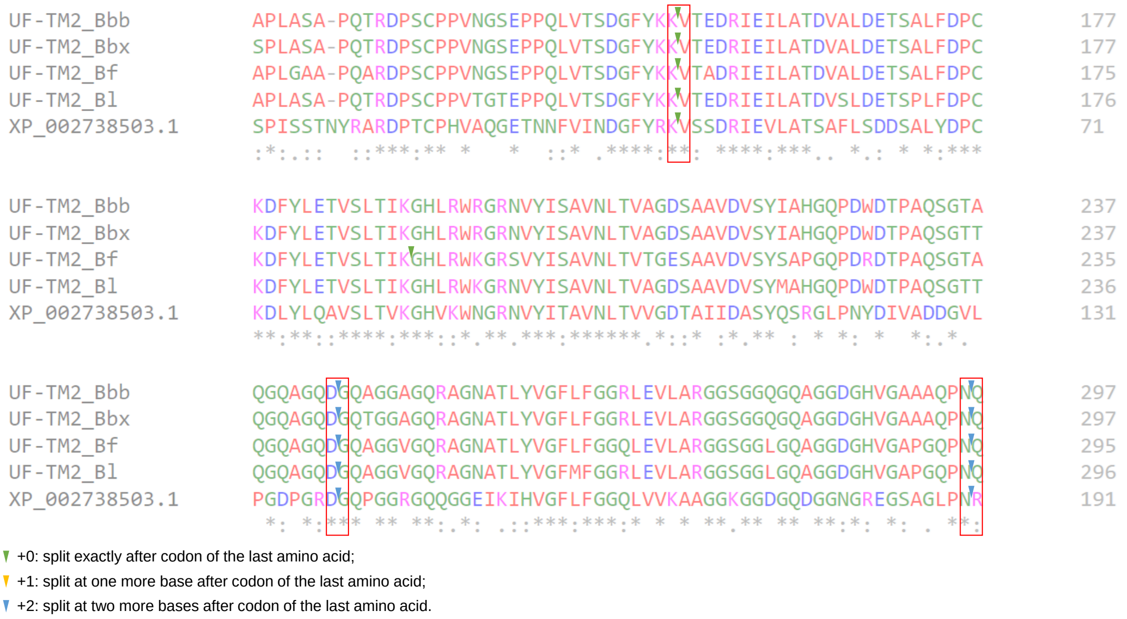


**Fig. S23. Partial sequence alignment of the transmembrane protein 2 of unknown function (UF-TM2) genes**

The protein sequences were aligned by the online tool Clustal Omega. XP_002738503.1 is the GenBank accession of the homologous protein from the hemichordate *Saccoglossus kowalevskii*. Three conserved splice sites were marked in red squares.

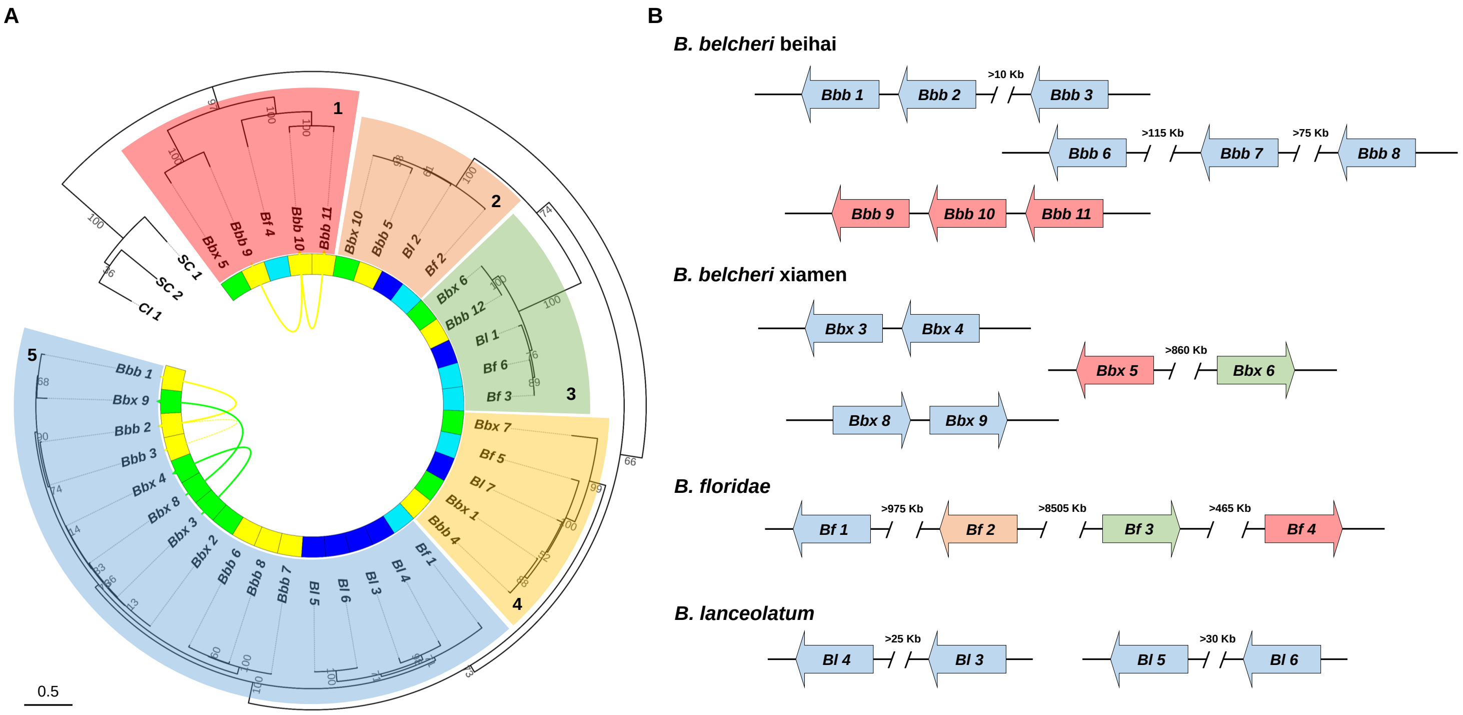


**Fig. S24. Comparative analysis of the transmembrane protein 3 of unknown function (UF-TM3) genes**

**(A)** Phylogenetic analysis of the UF-TM3 genes of amphioxus and two tunicates. SC1 and SC2, XP_039255317.1 and XP_039274118.1 of *Styela clava* respectively; CI1, XP_002128529.1 of *Ciona intestinalis*. Five clusters of DDC genes in amphioxus were classified and marked in different colors. **(B)** Gene synteny alignment of the UF-TM3 genes in four amphioxus genomes. The distance precision was set as 5 Kb, e.g., >75 Kb stand for 75-80 Kb.

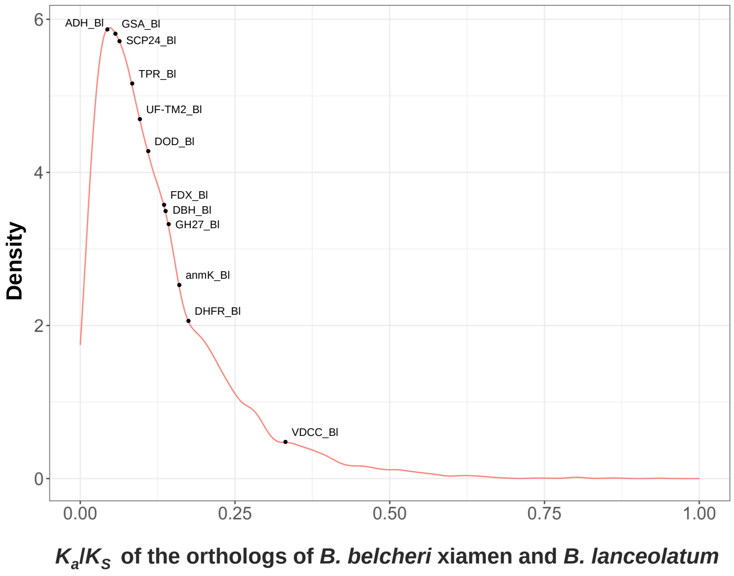


**Fig. S25. Selection pressure analysis of the orthologs of *B. belcheri* xiamen and *B. lanceolatum***
The nonsynonymous-to-synonymous substitution ratio (*Ka*/*Ks*) was calculated by the wgd pipeline and the CodeML program from the PAML package v4.9. The probability density distribution of *Ka*/*Ks* values for 6,667 one-to-one orthologs in the genomes of *B. belcheri* xiamen and *B. lanceolatum* was generated using ggplot2 in R and is displayed in the fitted curve. 12 one-to-one novel genes (except RBL that failed in this analysis, Table 1) are highlighted with black dots alongside *B. lanceolatum* genes (corresponding to those shown in Fig. S5), indicating their respective *Ka*/*Ks* values. All the *Ka*/*Ks* values for the 12 novel genes were below 0.34, with the highest value being 0.33 (VDCC_Bl). This pattern of low *Ka*/*Ks* values indicates a strong likelihood of negative selection, or purifying selection, acting on these genes.

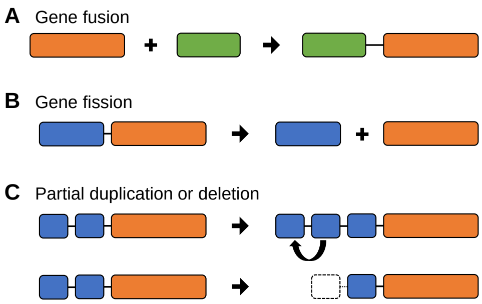


**Fig. S26. Models of novel gene generation**
Three models of novel gene generation after acquiring new genes of amphioxus were graphically illustrated. **(A)** Gene fusion, including two models fusing with an endogenous gene (or domain). **(B)** Gene fission. Two function domains in a novel gene were split into two genes. **(C)** Partial duplication or deletion. A novel gene containing multi-copy domains adds a domain via partial duplication or deletes a domain via partial deletion.

Marlétaz, F., Firbas, P.N., Maeso, I., Tena, J.J., Bogdanovic, O., Perry, M., Wyatt, C.D.R., de la Calle-Mustienes, E., Bertrand, S., Burguera, D., Acemel, R.D., van Heeringen, S.J., Naranjo, S., Herrera-Ubeda, C., Skvortsova, K., Jimenez-Gancedo, S., Aldea, D., Marquez, Y., Buono, L., Kozmikova, I., Permanyer, J., Louis, A., Albuixech-Crespo, B., Le Petillon, Y., Leon, A., Subirana, L., Balwierz, P.J., Duckett, P.E., Farahani, E., Aury, J.-M., Mangenot, S., Wincker, P.,

Albalat, R., Benito-Gutiérrez, È., Cañestro, C., Castro, F., D'Aniello, S., Ferrier, D.E.K., Huang, S., Laudet, V., Marais, G.A.B., Pontarotti, P., Schubert, M., Seitz, H., Somorjai, I., Takahashi, T., Mirabeau, O., Xu, A., Yu, J.-K., Carninci, P., Martinez-Morales, J.R., Crollius, H.R., Kozmik, Z., Weirauch, M.T., Garcia-Fernàndez, J., Lister, R., Lenhard, B., Holland, P.W.H., Escriva, H., Gómez-Skarmeta, J.L., Irimia, M., 2018. Amphioxus functional genomics and the origins of vertebrate gene regulation. Nature 564, 64-70.

Noon, J.B., Baum, T.J., 2016. Horizontal gene transfer of acetyltransferases, invertases and chorismate mutases from different bacteria to diverse recipients. BMC Evolutionary Biology 16, 74.

Pawson, T., Nash, P., 2003. Assembly of cell regulatory systems through protein interaction domains. Science 300, 445-452.

Putnam, N.H., Butts, T., Ferrier, D.E.K., Furlong, R.F., Hellsten, U., Kawashima, T., Robinson-Rechavi, M., Shoguchi, E., Terry, A., Yu, J.-K., Benito-Gutiérrez, E.l., Dubchak, I., Garcia-Fernàndez, J., Gibson-Brown, J.J., Grigoriev, I.V., Horton, A.C., de Jong, P.J., Jurka, J., Kapitonov, V.V., Kohara, Y., Kuroki, Y., Lindquist, E., Lucas, S., Osoegawa, K., Pennacchio, L.A., Salamov, A.A., Satou, Y., Sauka-Spengler, T., Schmutz, J., Shin-I, T., Toyoda, A., Bronner-Fraser, M., Fujiyama, A., Holland, L.Z., Holland, P.W.H., Satoh, N., Rokhsar, D.S., 2008. The amphioxus genome and the evolution of the chordate karyotype. Nature 453, 1064-1071.

Rotskaya, U.N., Kryukov, V.Y., Kosman, E., Tyurin, M., Glupov, V.V., 2021. Identification of the Ricin-B-Lectin LdRBLk in the Colorado Potato Beetle and an Analysis of Its Expression in Response to Fungal Infections. J Fungi (Basel) 7.

Schnell, J.R., Dyson, H.J., Wright, P.E., 2004. Structure, dynamics, and catalytic function of dihydrofolate reductase. Annu Rev Biophys Biomol Struct 33, 119-140.

Simakov, O., Kawashima, T., Marlétaz, F., Jenkins, J., Koyanagi, R., Mitros, T., Hisata, K., Bredeson, J., Shoguchi, E., Gyoja, F., Yue, J.-X., Chen, Y.-C., Freeman, R.M., Sasaki, A., Hikosaka-Katayama, T., Sato, A., Fujie, M., Baughman, K.W., Levine, J., Gonzalez, P., Cameron, C., Fritzenwanker, J.H., Pani, A.M., Goto, H., Kanda, M., Arakaki, N., Yamasaki, S., Qu, J., Cree, A., Ding, Y., Dinh, H.H., Dugan, S., Holder, M., Jhangiani, S.N., Kovar, C.L., Lee, S.L., Lewis, L.R., Morton, D., Nazareth, L.V., Okwuonu, G., Santibanez, J., Chen, R., Richards, S., Muzny, D.M., Gillis, A., Peshkin, L., Wu, M., Humphreys, T., Su, Y.-H., Putnam, N.H., Schmutz, J., Fujiyama, A., Yu, J.-K., Tagawa, K., Worley, K.C., Gibbs, R.A., Kirschner, M.W., Lowe, C.J., Satoh, N., Rokhsar, D.S., Gerhart, J., 2015. Hemichordate genomes and deuterostome origins. Nature 527, 459-465.

Simão, F.A., Waterhouse, R.M., Ioannidis, P., Kriventseva, E.V., Zdobnov, E.M., 2015. BUSCO: assessing genome assembly and annotation completeness with single-copy orthologs. Bioinformatics 31, 3210-3212.

Stamatakis, A., 2014. RAxML version 8: a tool for phylogenetic analysis and post-analysis of large phylogenies. Bioinformatics 30, 1312-1313.

Uehara, T., Suefuji, K., Valbuena, N., Meehan, B., Donegan, M., Park, J.T., 2005. Recycling of the anhydro-N-acetylmuramic acid derived from cell wall murein involves a two-step conversion to N-acetylglucosamine-phosphate. J Bacteriol 187, 3643-3649.

Wang, C.F., Sun, W., Zhang, Z., 2019. Functional characterization of the horizontally transferred 4,5-DOPA extradiol dioxygenase gene in the domestic silkworm, Bombyx mori. Insect Mol Biol 28, 409-419.

Wibowo, A.S., Singh, M., Reeder, K.M., Carter, J.J., Kovach, A.R., Meng, W., Ratnam, M., Zhang, F., Dann, C.E., 3rd, 2013. Structures of human folate receptors reveal biological trafficking states and diversity in folate and antifolate recognition. Proc Natl Acad Sci U S A 110, 15180-15188.

Xu, X., Zhang, L., Zhao, W., Fu, L., Han, Y., Wang, K., Yan, L., Li, Y., Zhang, X.-H., Min, D.-H., 2021. Genome-wide analysis of the serine carboxypeptidase-like protein family in Triticum aestivum reveals TaSCPL184-6D is involved in abiotic stress response. BMC Genomics 22, 350.

Yang, K.Y., Chen, Y., Zhang, Z., Ng, P.K.-S., Zhou, W.J., Zhang, Y., Liu, M., Chen, J., Mao, B., Tsui, S.K.-W., 2016.

Transcriptome analysis of different developmental stages of amphioxus reveals dynamic changes of distinct classes of genes during development. Scientific Reports 6, 23195.
Yang, Z., 2007. PAML 4: phylogenetic analysis by maximum likelihood. Molecular biology and evolution 24, 1586-1591.
Zhang, Q.-L., Zhu, Q.-H., Liao, X., Wang, X.-Q., Chen, T., Xu, H.-T., Wang, J., Yuan, M.-L., Chen, J.-Y., 2016. Selection of reliable reference genes for normalization of quantitative RT-PCR from different developmental stages and tissues in amphioxus. Scientific Reports 6, 37549.
